# A Human Accelerated Region Drives Opposing Heterochronic Changes in Craniofacial and Limb Development

**DOI:** 10.64898/2026.09.08.750211

**Authors:** Yu Ji, Matheo Morales, Marybeth Baumgartner, James P. Noonan

## Abstract

Many human developmental processes proceed over a prolonged timescale compared to other primates, a phenomenon known as heterochrony. Human accelerated regions (HARs), which encode transcriptional enhancers with human-specific activity, have been implicated in the evolution of novel human traits. However, their contributions to changes in developmental timing remain unknown. Using single-nucleus RNA sequencing and developmental trajectory analyses in a genetically humanized mouse model of the HAR *HACNS1*, we show that *HACNS1* drives heterochronic shifts in opposite directions during craniofacial and limb development. In pharyngeal arches, *HACNS1* delays chondrocyte differentiation and the expression of genes involved in cartilage and skeletal maturation. In contrast, in the limb buds, *HACNS1* accelerates chondrogenesis and promotes earlier expression of differentiation-associated genes. Consistent with these transcriptomic shifts, SOX9-expressing pre-cartilaginous domains are more diffuse in the pharyngeal arches, but the condensed domains are expanded in the limb buds. Gene regulatory network inference suggests that *HACNS1*-driven heterochronic shifts are due to changes in the expression of its target gene *Gbx2* and resulting downstream effects on the regulatory networks through which *Gbx2* functions. Our findings demonstrate that a single human-specific gene regulatory change can alter developmental timing, providing a potential mechanism for how uniquely human genetic changes reshaped conserved developmental programs.

## Introduction

Development proceeds through the tightly regulated spatiotemporal progression of cell fate specification, maturation, organogenesis, and morphogenesis. Changes in the timing of development across species, termed heterochrony, are a fundamental mechanism driving evolutionary innovation^1–3^. The evolution of many uniquely human traits involved heterochronic shifts in the emergence, rate, or duration of developmental events^4,5^. Insight into the genetic mechanisms that shift developmental timing *in vivo* remains limited. Recent studies have shown that human-specific gene duplicates drive changes in developmental rates, particularly in the brain. For example, the human-specific gene duplicate *SRGAP2C* delays synaptic development^6,7^. However, whether human-specific gene regulatory changes also drive heterochronic shifts during development is unknown.

More than 20,000 human-specific genetic changes that potentially altered gene expression have been identified over the past two decades^8^. One prominent class, termed Human Accelerated Regions (HARs), consists of genomic elements highly conserved across many species but that show a significant excess of human-specific sequence changes^9–12^. Many HARs have been shown to exhibit human-specific transcriptional enhancer activity during development, suggesting they contributed to the evolution of uniquely human traits by altering gene expression^13–21^. The first HAR demonstrated to encode human-specific enhancer activity is *HACNS1* (also known as *HAR2*). In a transgenic mouse enhancer assay, *HACNS1* exhibited human-specific activity in developing limb buds and in the mandibular processes (Md) of the first pharyngeal arch and the second pharyngeal arch (PA2) compared with its chimpanzee ortholog^13^. A subsequent study using an *HACNS1* humanized mouse model showed that *HACNS1* maintained its human-specific enhancer activity in the mouse context and altered the expression level and spatial distribution of the nearby gene *Gbx2* in the limb and the pharyngeal arches (PAs), although no overt morphological phenotype was observed^22^.

*Gbx2* encodes a transcription factor that functions in multiple tissues during development. In the PAs, *Gbx2* is expressed in the proximal mesenchyme and is required for proper neural crest patterning, with loss of *Gbx2* leading to defects in pharyngeal arch-derived structures and craniofacial abnormalities. *Gbx2*-derived cells contribute to all major lineages of the mandible^23–25^. *Gbx2* is transiently expressed in the posterior-middle region of the limb bud at embryonic day 9.5 (E9.5) and becomes nearly undetectable by E11.5, but its role in limb development is unknown^26^. How *HACNS1*-driven changes in *Gbx2* expression might have altered downstream gene regulatory networks and the development of the PAs and limb buds remains unclear.

Here we characterized *HACNS1* enhancer activity during early chondrogenic progression in both humanized mouse craniofacial and limb mesenchyme and found that *HACNS1* shifted the timing of chondrogenic fate commitment and mesenchymal condensation. Analyses of developmental trajectories inferred from longitudinal single-nucleus transcriptome data revealed opposite heterochronic effects across tissues. *HACNS1* delayed chondrogenic fate commitment in the Md and PA2 while it accelerated these programs in the limb buds, revealing tissue-specific changes in developmental timing. Consistent with these gene-expression shifts, SOX9-positive pre-cartilaginous condensation domains at E11.5 showed divergent morphologies: more diffuse in the Md and PA2, but the condensed domains were expanded in the limb buds compared to wild type, supporting heterochronic changes in chondrocyte differentiation. SOX9 is a master regulator of chondrogenesis and an early marker of pre-cartilaginous mesenchymal condensations^27,28^, and changes in SOX9*-*positive domains provide a morphological indicator of altered chondrogenic timing. Together, these results implicate *HACNS1-*mediated modification of the *Gbx2* regulatory pathway as a mechanism through which a uniquely human genetic change could differentially modulate the course of chondrocyte differentiation during human evolution.

## Results

### Characterizing the temporal pattern of *HACNS1* enhancer activity during mouse embryonic development

Previous studies using mouse transgenic enhancer assays and genetically modified mouse models demonstrated that *HACNS1* exhibited human-specific enhancer activity in the E11.5 Md, PA2, and limb buds, with *HACNS1* notably showing increased levels of the enhancer activity-associated histone modification H3K27ac in the limb compared to the chimpanzee and mouse orthologs^13,22^. However, the developmental time span during which *HACNS1* may be active in these tissues has not been characterized, which is an essential first step towards understanding its effects on developmental rates. We performed assays for transposase-accessible chromatin using high-throughput sequencing (ATAC-seq) on Md, PA2, and limb bud tissues collected at one-day intervals from E9.5 to E12.5 from both the previously described *HACNS1* humanized mouse line and a control line in which the endogenous locus was replaced with the chimpanzee ortholog. The time period we examined includes key morphogenetic events, including PA patterning and the emergence of Meckel’s cartilage, which provides a transient blueprint that organizes subsequent mandibular morphogenesis^29^, as well as limb bud outgrowth and segmentation^30^. We did not collect data for the hindlimb at E9.5, as it has yet to emerge, or the PAs at E12.5, when the mandibular process is one structure.

Our results indicated that both *HACNS1* and its chimpanzee ortholog exhibited significant open chromatin accessibility in the Md, PA2, and forelimb as early as E9.5. Across all tissues examined, chromatin accessibility at *HACNS1* peaked at E10.5, declined starting at E11.5, and was substantially reduced by E12.5 (Fig. 1A, B; Fig. S1). Compared with the chimpanzee ortholog, the human *HACNS1* ortholog exhibited increased chromatin accessibility at earlier time points. Specifically, at E9.5, accessibility at *HACNS1* was significantly higher in PA2 and the forelimb (Fig. 1B; Fig. S1). Our data indicate that both human and chimpanzee orthologs remain accessible from E9.5 to E12.5 but with reduced accessibility at later stages, suggesting their enhancer activity is strongest earlier in development. We note that *HACNS1* did not show significantly increased accessibility compared to the chimpanzee ortholog at every time point and in every tissue. However, this does not imply the orthologs necessarily have similar enhancer activity, as previous studies have shown that human *HACNS1* drives stronger enhancer activity than the chimpanzee ortholog at E11.5^22^.

**Fig. 1.**
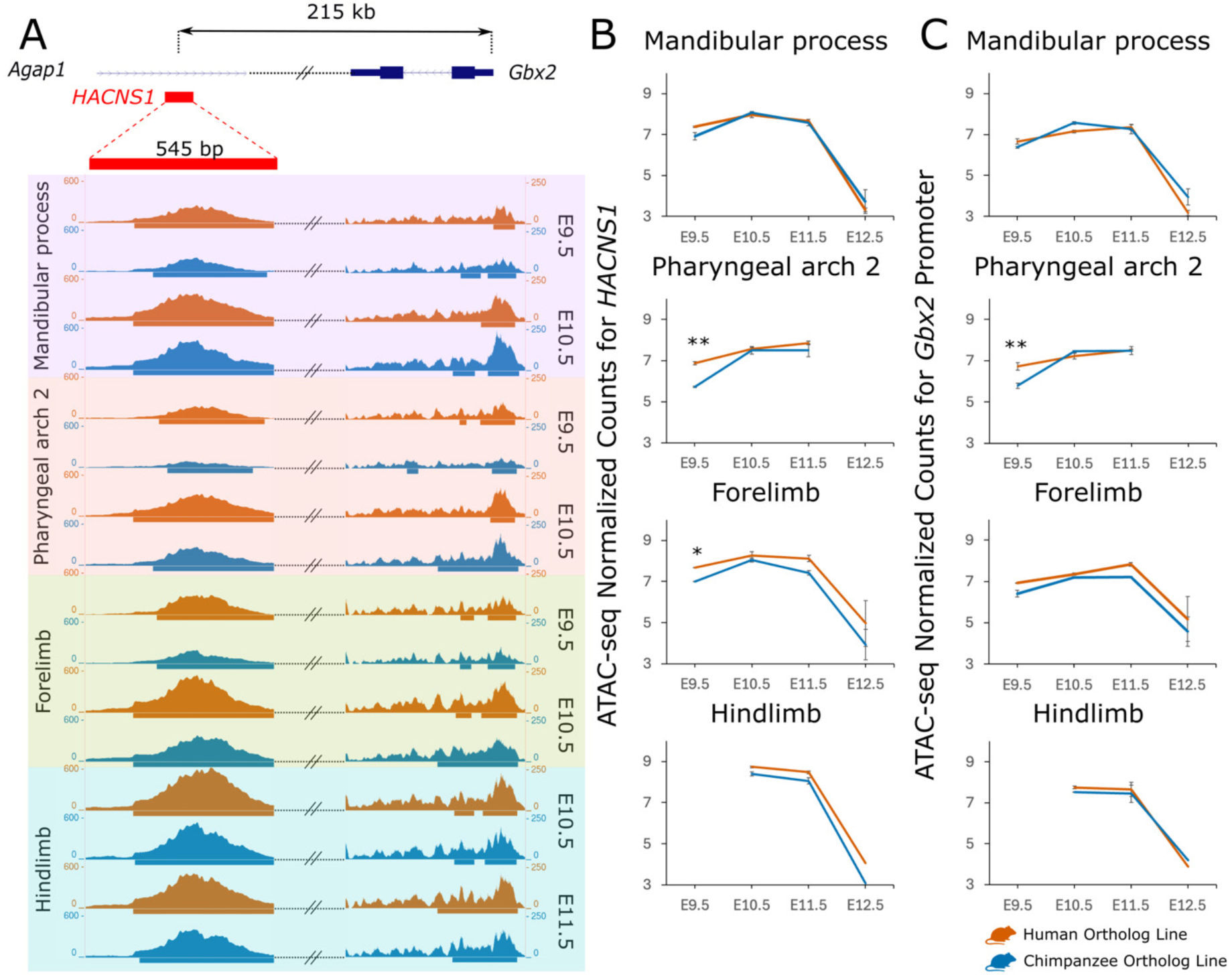
Increased chromatin accessibility at *HACNS1* in humanized mouse embryos. **A:** ATAC-seq signal at the *HACNS1* locus (left) and the *Gbx2* promoter (right) in the *HACNS1* humanized line (dark orange) and a line in which the chimpanzee ortholog has precisely replaced the endogenous mouse ortholog (blue) in the mandibular process (lavender shading), second pharyngeal arch (peach), forelimb (light green), and hindlimb (cyan) at the indicated time points. The genomic location of *HACNS1* (in red) in the humanized line relative to *Gbx2* (in blue) and the ATAC-seq signal at each time point is indicated at the top of the panel. Peak calls showing significantly accessible regions are indicated by bars below each signal track. See Methods for details on the genome assembly used for alignment and peak calling. ATAC-seq signals obtained for all tissues and time points are shown in Supplementary Fig. 1. **B, C:** Comparison of normalized ATAC-seq signal at the *HACNS1* locus **(B)** and *Gbx2* promoter **(C)** between the *HACNS1* humanized line (dark orange) and the chimpanzee ortholog line (blue). Significant differences in accessibility between genotypes were identified using DESeq2 implemented in DiffBind^31,32^ (*FDR < 0.05; **FDR < 0.01).

To test whether the human-specific earlier accessibility of *HACNS1* is associated with enhanced regulatory activity, we next examined chromatin accessibility at the promoter of *Gbx2*, previously identified as a regulatory target of *HACNS1* in the Md, PA2, and limb buds^22^. Similar to *HACNS1*, the *Gbx2* promoter is accessible at E9.5 and reaches peak accessibility at E10.5-E11.5 in the Md, PA2, and forelimb. Accessibility is reduced by E12.5 in both the Md and limb buds. Compared to the chimpanzee ortholog line, the *Gbx2* promoter shows significantly higher accessibility in *HACNS1* humanized mouse embryos at E9.5 in PA2 (Fig. 1C, Fig. S1), supporting that the earlier chromatin accessibility of human *HACNS1* is associated with earlier upregulation of its downstream target gene. Together, these results suggest that *HACNS1* functions as a temporally specific enhancer active from E9.5 to E11.5, with the human ortholog exhibiting stronger enhancer activity than the chimpanzee ortholog. This finding is consistent with the previously reported increase in *HACNS1* enhancer activity and *Gbx2* expression levels at E11.5 in the *HACNS1* humanized mouse model^22^.

### Single-nucleus transcriptome profiling of *HACNS1* humanized and chimpanzee ortholog mouse embryonic pharyngeal arches and limb buds

To assess the impact of *HACNS1* on gene expression and development, we performed single-nucleus RNA sequencing (snRNA-seq) in *HACNS1* humanized and chimpanzee ortholog mouse embryonic Md, PA2, and limb buds at 1-day intervals from E9.5 to E12.5. At each time point, we harvested at least eight embryos and collected the Md, PA2, forelimb, and hindlimb from the same embryo. In total, we generated 28 snRNA-seq datasets from *HACNS1* humanized and chimpanzee ortholog mouse embryos (Fig. 2A). After quality control filtering (Methods), 299,553 nuclei were retained for downstream analyses (Fig. S2; Table S1). Following normalization using SCTransform, integration of datasets across genotypes using Harmony, and visualization of the integrated data using uniform manifold approximation and projection (UMAP), we identified cell clusters for all tissues at each time point using the Louvain algorithm. Cell clusters were annotated based on the expression of known cell type markers (Figs. S3-12; Table S2). In the Md and PA2, we identified major cell types including neural crest-derived mesenchymal cells (*Dlx5*, *Prrx1*), epithelial cells (*Epcam*, *Cdh1*), oral epithelium (*Fgf8*, *Vgll2*), pharyngeal cleft cells (*Wnt6*, *Tfap2b*), mesoderm-derived cells (*Tbx1*, *Pax7*) and endothelial cells (*Cdh5*, *Cldn5*) (Fig. S3, 4)^24^. In the limb buds, we identified mesenchymal cells (*Twist1*, *Prrx1*), epithelial cells (*Epcam*, *Cdh1*), and apical ectodermal ridge cells (*Fgf8*) (Fig.S7, 8)^33,34^.

**Fig. 2.**
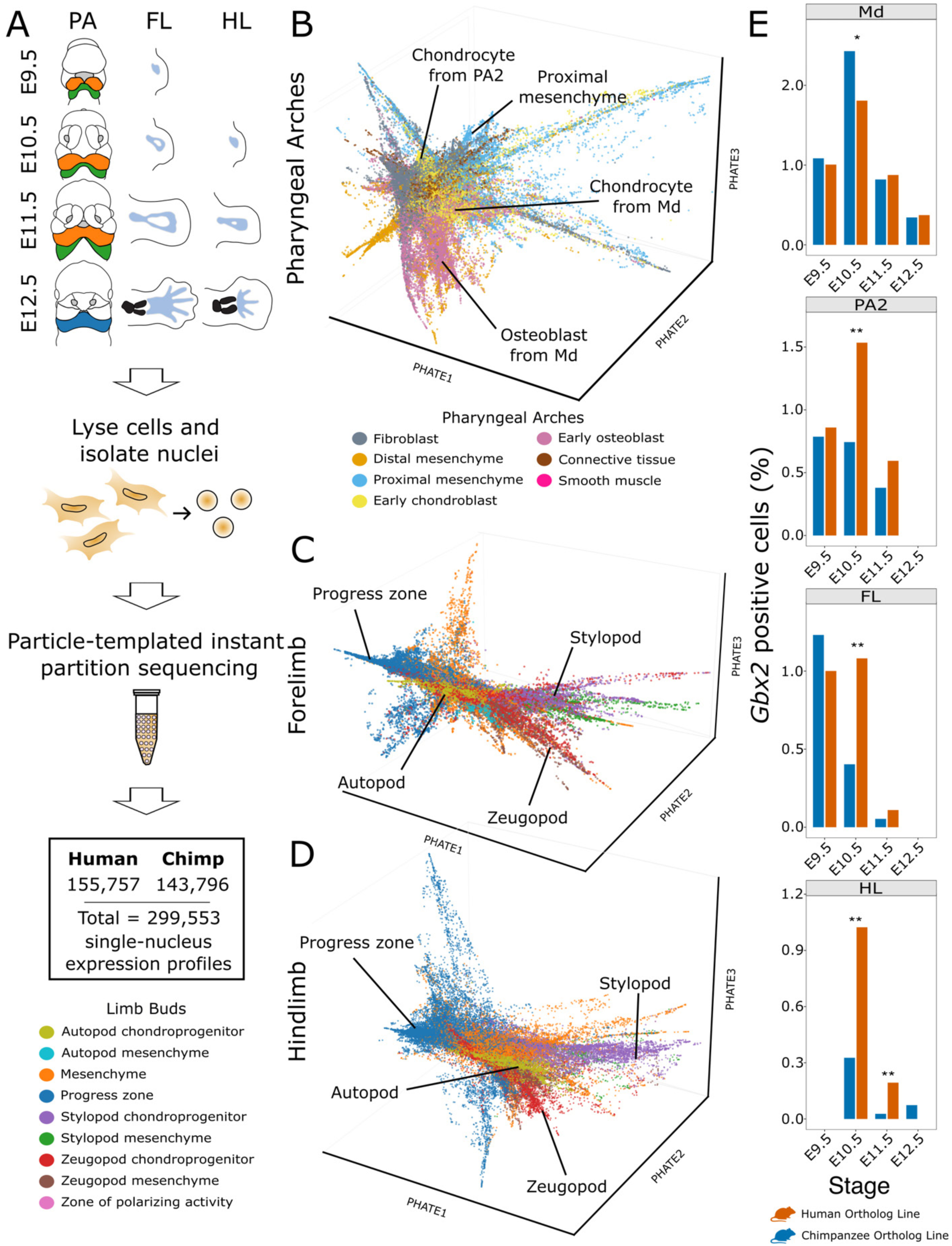
Single-nucleus transcriptomics reveals mesenchymal developmental lineages in pharyngeal arches and limb buds of *HACNS1* humanized and chimpanzee ortholog mouse embryos. A: Experimental design and tissue collection scheme for snRNA-seq. Embryos were collected from E9.5 to E12.5, with dissected tissues indicated by color: Md and PA2 from E9.5 to E11.5 are shown in orange and green, and Md at E12.5 is shown in blue. Forelimbs were collected from E9.5 to E12.5, and hindlimbs were collected from E10.5 to E12.5. The workflow for nuclei isolation, particle-templated instant partition (PIP) sequencing, and the number of high-quality nuclei obtained from each genotype are shown below. **B-D:** Three-dimensional PHATE embeddings of mesenchymal cells from the pharyngeal arches **(B)**, forelimb **(C)**, and hindlimb **(D)**. Cells are colored by broadly defined cell-type clusters for visualization as indicated in Fig 2A. Text labels on each plot point to the locations of major lineages in each tissue. **(E)** Bar plots show the percentage of *Gbx2*-positive nuclei in each indicated tissue and genotype. Differences between genotypes were assessed using two-tailed Fisher’s exact test, with P values adjusted for multiple testing using the Benjamini-Hochberg method^42^. * p_adj < 0.05; ** p_adj < 0.01.

A previous study found that *HACNS1* upregulated *Gbx2* expression in chondrogenic mesenchyme^22^. Therefore, we focused our subsequent analyses on mesenchymal cell populations across all datasets. In the Md and PA2, single-nucleus transcriptomic profiles clearly resolved fate-undecided mesenchymal populations patterned along both the proximal-distal and cranial-caudal axes, particularly at early developmental stages such as E9.5. These axial domains are evident from the expression of known marker genes across the embeddings, which show expression gradients revealing distinct proximal-distal and cranial-caudal patterning identities (Fig. S5A-G; Fig. S6A-F). From E10.5 to E12.5, these patterned mesenchymal populations progressively give rise to fate-committed cell types, including pre-chondrocytes (*Sox9*, *Sox5*, *Sox6*), pro-osteoblasts (*Runx2*), connective tissue cells (*Nr5a2*), and fibroblasts (*Dlk1*) (Fig. S5H; Fig. S6G)^35,36,24,37^, indicating that the major axes of PA patterning and differentiation are captured in the data.

We also observed a similar resolution of axial patterning in the limb bud datasets. Single-nucleus profiles delineated clear proximal-distal and anterior-posterior domains defined by established marker genes, with undifferentiated mesenchymal stem cells enriched in the distal progress zone at E9.5 (*Fgf10*, *Msx1*) (Fig. S9-12). From E10.5 onward, chondrogenic lineages emerged in a spatially ordered manner along the distal-proximal axis. *Sox9* marked the emergence of stylopod, zeugopod, and autopod chondroblast populations at E10.5 and E11.5. By E12.5, the osteoblast marker *Runx2* was expressed in the proximal stylopod domain with concomitant reduction in *Sox9* expression in this region, highlighting the transition from chondrogenesis to osteogenesis (Fig. S10-12)^38,39^. These results demonstrate that the principal axes and lineage hierarchies of limb patterning are faithfully captured by our single-nucleus transcriptomic profiles.

We next sought to infer mesenchymal developmental trajectories in the Md, PA2, and limb buds using our snRNA-seq data to identify potential changes driven by *HACNS1*. We first integrated mesenchymal cell populations across developmental stages using Harmony. We analyzed forelimb and hindlimb mesenchymal cells separately. However, we integrated mesenchymal cells from the Md and PA2 across all time points for downstream analyses, as progenitors in both arches can contribute to the same downstream anatomical structures, making it difficult to infer developmental trajectories for each arch separately. For example, although Meckel’s cartilage and Reichert’s cartilage are derived from the first and second pharyngeal arches, respectively, multiple studies have shown that mixed neural crest cell populations associated with both arches also contribute to anatomical derivatives such as middle ear ossicular elements^40^.

To visualize the developmental dynamics of these integrated datasets and capture continuous developmental trajectories, we then applied PHATE^41^ to the Harmony-corrected low-dimensional representation (Fig. 2B-D). This approach allowed us to detect transcriptional transitions among mesenchymal cells while preserving the broader organization of developmental cell states across stages. In the PAs, including the Md and PA2, cells in the PHATE embedding segregated primarily by tissue identity, while developmental stage accounted for much of the variation within each tissue (Fig. S13A). Genotype had little apparent effect on cell distribution, supporting that overall mesenchymal cell-type composition is similar in the *HACNS1* humanized and chimpanzee ortholog lines (Fig. S13A). The position of cells in the PHATE embedding was consistent with their relative maturation state and degree of fate commitment (Fig. 2B; Fig. S13). Proximal mesenchymal cells clustered on one side of the embedding. In contrast, fate-committed cells occupied the opposite side and branched into distinct lineages, including chondroblast, osteoblast, connective tissue, and fibroblast lineages. We also found that *Gbx2* was expressed in the proximal mesenchyme of the Md and PA2 (Fig. S13A). At E10.5, the Md and PA2 in *HACNS1* humanized embryos contain a significantly higher percentage of *Gbx2*-positive cells than the corresponding tissues in chimpanzee control embryos (Fig. 2E, top; Fig. S13A; Movie S1), consistent with *HACNS1* driving increased *Gbx2* expression.

In both forelimb and hindlimb, cells displayed similar overall distributions, with developmental stage accounting for the majority of observed variation. Cells did not separate by genotype (Fig. S13B, C). The position of cells in the limb PHATE embeddings was also consistent with progressive maturation and fate commitment (Fig. 2C, D). Cells at one end of the embedding predominantly corresponded to undifferentiated progress zone cells. At the opposite end, the chondroblast population branched into three major anatomical lineages: the stylopod, zeugopod, and autopod, which give rise to the proximal, intermediate, and distal segments of the limb, respectively. In the limb buds, *Gbx2* is expressed in the progress zone (Fig. S13B, C). At E10.5 in the forelimb and at E10.5-E11.5 in the hindlimb, *HACNS1* humanized embryos contain a significantly higher number of *Gbx2*-positive cells than chimpanzee control embryos (Fig. 2E, bottom; Fig. S13B, C; Movie S2, 3). Together, these results indicate that the known cellular identities and spatial patterning of PAs and limb buds are captured in our snRNA-seq data, providing the means to understand how *HACNS1* may have altered the timing and trajectory of chondroblast differentiation.

### Inferring mesenchymal developmental trajectories in the pharyngeal arches and limb buds

After identifying the major mesenchymal cell populations and their broad organization from undifferentiated to fate-committed states, we next inferred the developmental trajectories of differentiating mesenchymal cells within the *HACNS1* humanized and chimpanzee ortholog line PAs and limb buds from our snRNA-seq datasets. This allowed us to reconstruct the process of skeletal differentiation and link cells to specific developmental transitions. A challenge in our analysis is that mesenchymal populations in the PAs and limb buds are organized both by spatial patterning and developmental stage as they progress toward distinct skeletal fates. To extract these complex developmental processes, we developed an integrated pipeline that combines URD, PHATE, and Slingshot (Fig. 3A)^43,44,41^. URD reconstructs developmental trajectories by using simulated random walks on a cell-cell transition graph to estimate the likelihood that individual cells connect early progenitor states to specified terminal cell fates. We first applied URD to the integrated mesenchymal cell populations to identify cells with high predicted connectivity between early mesenchymal progenitors and later skeletal states (Fig. S14A). We selected osteoblasts or chondrocytes from E12.5 samples as terminal states. These populations represent the most differentiated skeletal cell states captured in our snRNA-seq time series and provide appropriate terminal points for reconstructing progression from early mesenchyme toward chondrogenic and osteogenic fates^45,46^. We defined developmental starting points as early mesenchymal progenitor populations: proximal mesenchyme at E9.5 in the Md and PA2, which contributes to all major mandibular lineages, and progress zone cells at E9.5 in the forelimb or E10.5 in the hindlimb, which represent distal undifferentiated mesoderm that give rise to skeletal elements along the proximal-distal axis. We then used URD to identify mesenchymal cells with a high likelihood of progressing toward skeletal lineages, which were retained for subsequent trajectory and pseudotime analyses (Fig. S14A).

**Fig. 3.**
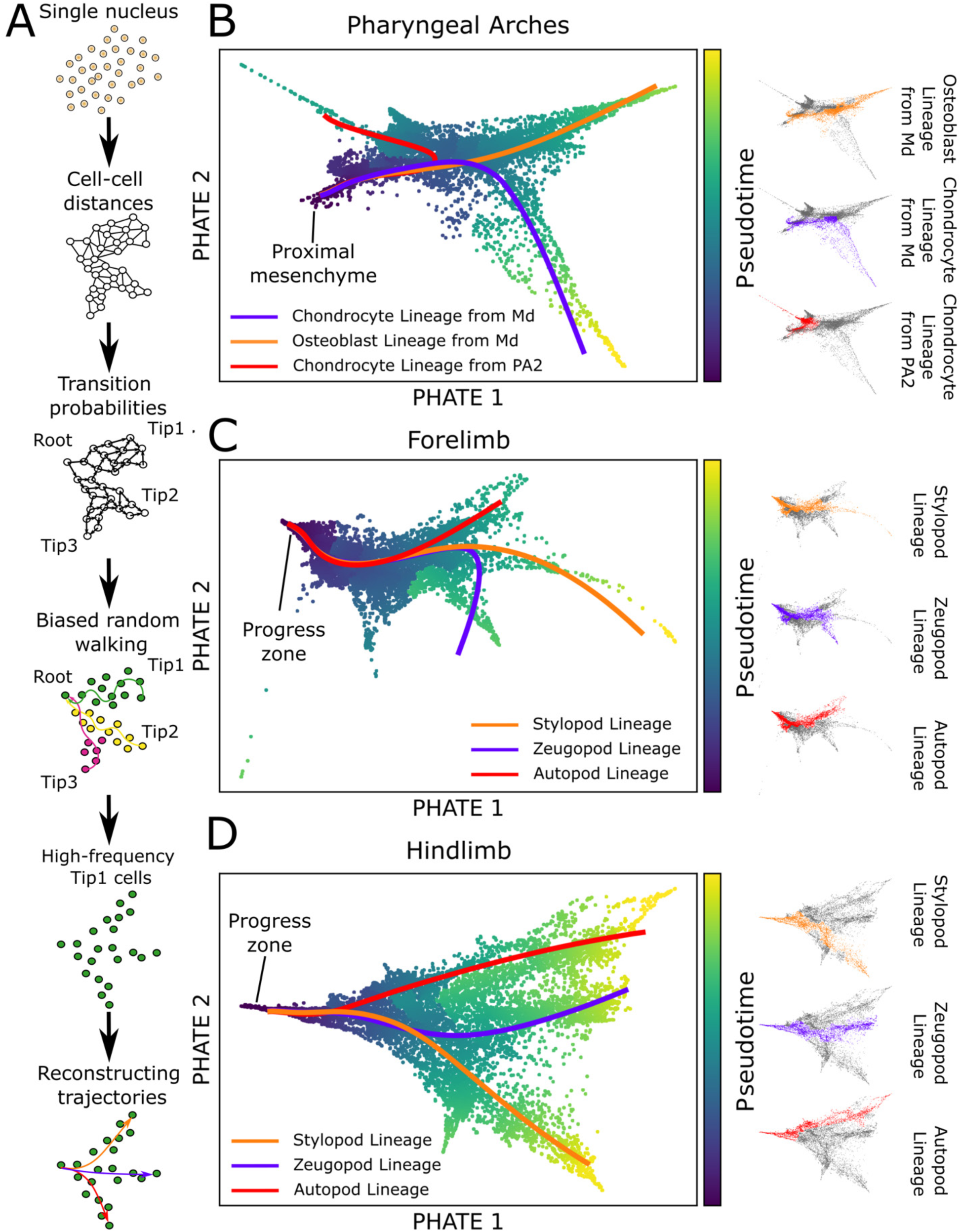
Reconstruction of mesenchymal developmental trajectories in the pharyngeal arches and limb buds of *HACNS1* humanized and chimpanzee ortholog mouse embryos. A: Schematic of the workflow used to reconstruct mesenchymal differentiation trajectories from snRNA-seq data. See Methods for details. B-D: *Left*. PHATE embeddings of mesenchymal cells identified by URD as associated with chondroblast or osteoblast differentiation in the pharyngeal arches **(B)**, forelimb **(C)**, and hindlimb **(D)**. Cells are colored by pseudotime values as shown on the scale to the right of each panel. Bold lines indicate lineage-specific trajectories resolved by Slingshot. The location of undifferentiated mesenchymal cells (the proximal mesenchyme in the PAs and the progress zone in the forelimb and hindlimb) is labeled in each panel. *Right*. Each panel shows cells in each embedding colored by their assigned trajectories, with cells not assigned to the indicated trajectories shown in gray.

Although URD identified mesenchymal cells associated with skeletal lineages, it did not resolve anatomically distinct populations. Because cartilage and skeletal populations in different PA domains and limb segments follow separate developmental trajectories, we used PHATE to further distinguish these populations (Fig. 3; Fig. S14). The PHATE embedding revealed clear separation between skeletal populations derived from the mandibular process and PA2, as well as between the stylopod, zeugopod, and autopod populations in the limb buds (Fig. S14B). Finally, we applied Slingshot to these PHATE embeddings to define lineage-specific trajectories and rank the maturation of cells along the trajectories by pseudotime, with increasing pseudotime values corresponding to progressive maturation along each trajectory (Fig. 3B-D). This analysis revealed regionally specific trajectories within Md and PA2 as well as segment-specific trajectories in the limb buds. Across all tissues, cells from earlier developmental stages were assigned lower pseudotime values, whereas cells from later stages were assigned higher pseudotime values: E9.5 cells were located at the beginning of each inferred trajectory and E12.5 cells at the end. Consistent with this temporal ordering, cells were also ordered by their degree of maturation: Undifferentiated mesenchymal stem cells were assigned lower pseudotime values, while fate-determined cell populations were assigned higher pseudotime values. These results indicate that the inferred trajectories accurately recapitulate the known mesenchymal differentiation process. We did not observe cells separating along pseudotime by genotype, indicating that the overall trajectory of chondrogenic fate commitment is similar between genotypes (Fig. S14B).

To further evaluate whether our inferred trajectories captured *bona fide* developmental processes, we examined the expression of genes known to be involved in bone formation along pseudotime. In the Md and PA2, genes that regulate mesenchymal fate determination and neural crest stemness, including *Id1*, *Dlx1*, and *Prrx1*^47–51^, were highly expressed at the beginning of the trajectories and progressively decreased along pseudotime (Fig. S15A). In contrast, genes associated with bone and cartilage formation, including collagen genes such as *Col1a2* and *Col5a2*, and the osteogenic transcription factor *Runx2*, were expressed toward the end of the trajectories (Fig. S15A). Similarly, in the limb buds, genes that maintain the undifferentiated progress zone, such as *Fgf10*, were expressed at the beginning of the trajectories and decreased toward the end, whereas *Sox9*, a key regulator of chondrogenic differentiation, showed a steady increase along pseudotime (Fig. S15B, C).

Because genes that exhibit coordinated expression changes across development are likely to participate in related biological processes, we next used Monocle3 to identify modules of genes showing similar changes in their expression along each trajectory as a function of pseudotime^52^ (Methods). Gene Ontology (GO) enrichment analysis of these modules revealed that terms associated with skeletal system development, cartilage condensation, and extracellular matrix organization were consistently enriched in modules whose expression peaks at the end of each trajectory (Fig. S16-18; Tables S3, S4). Together, these results support that we successfully reconstructed the developmental trajectories of early osteoblasts and chondrocytes in both the Md, PA2, and limb buds.

### Transcriptional trajectory alignments reveal heterochronic shifts in pharyngeal arch and limb bud development

After reconstructing mesenchymal differentiation trajectories, we next investigated whether *HACNS1* altered the timing and distribution of gene expression along these trajectories. We aligned cells assigned to each trajectory we examined from *HACNS1* humanized and chimpanzee control mice using cellAlign^53^ (Fig. 4). Although Md and PA2 were analyzed together to identify cell types, cells assigned to trajectories called by Slingshot within each region were extracted and aligned separately for this analysis (Fig. 3B). We then aligned the genotype-specific pseudotime trajectories using the top 3000 highly variable genes identified by SCTransform, enabling us to compare developmental timing across genotypes. If these lineages developed at similar rates in the *HACNS1* humanized mouse and chimpanzee control lines, we anticipated their transcriptional trajectories would align closely with minimal pseudotemporal shift. Consistent with this expectation, cells from both genotypes were largely ordered by developmental time point along all of the trajectories (Fig. 4). Trajectories from the Md and forelimb aligned closely between *HACNS1* humanized and chimpanzee control lines (Fig. 4A, B, D, E, F). Notably, although *HACNS1* exhibits increased accessibility in the forelimb at E9.5, this difference does not lead to a clear heterochronic shift at the transcriptional level.

**Fig. 4.**
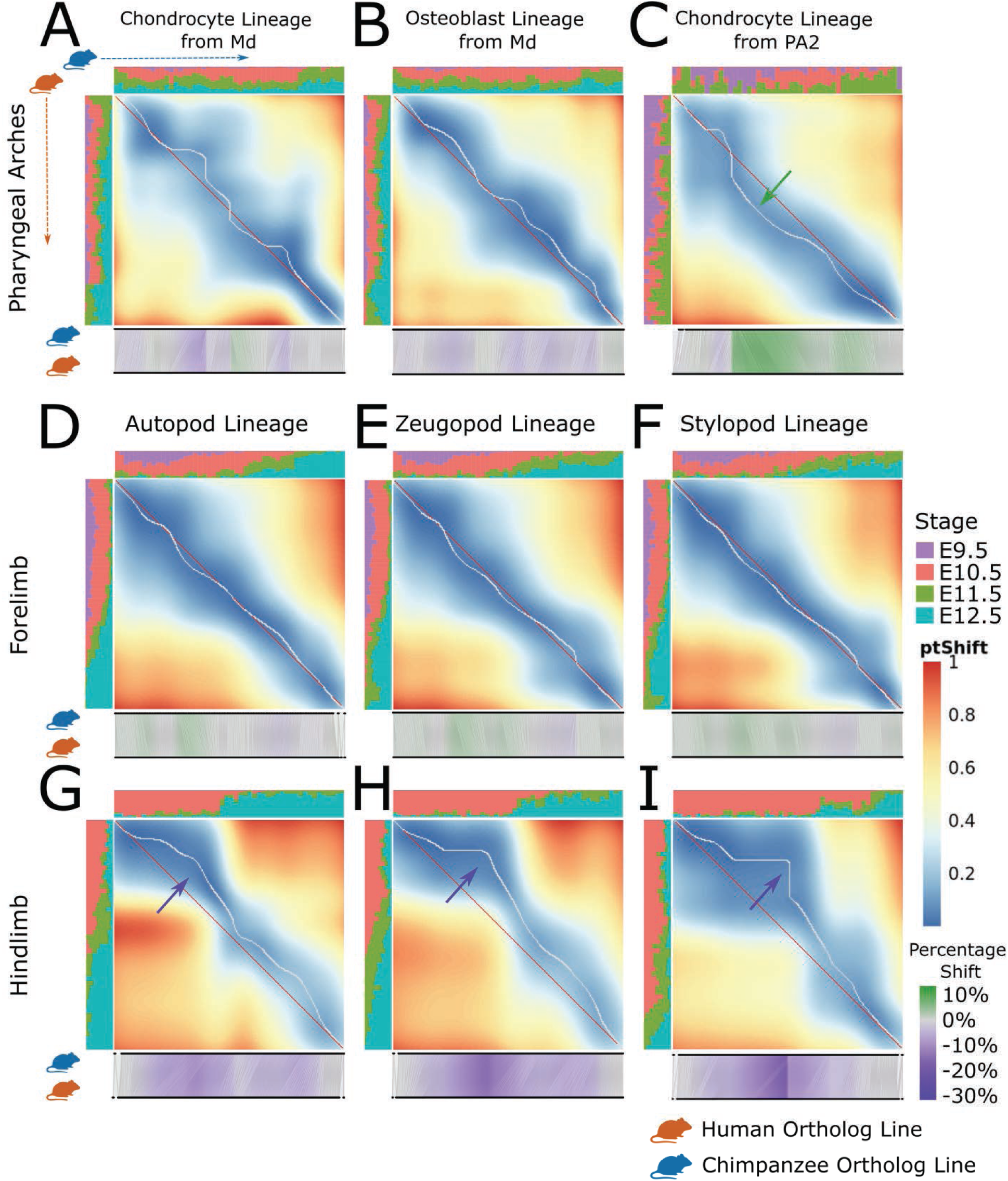
Inferred developmental trajectory alignments reveal heterochronic shifts in pharyngeal arch and limb bud development in *HACNS1* humanized mice. Pseudotime alignment of inferred developmental trajectories between *HACNS1* humanized and chimpanzee ortholog lines for the pharyngeal arches **(A-C)**, forelimb **(D-F)**, and hindlimb **(G-I)**. The middle panels show the degree of transcriptional similarity over pseudotime in *HACNS1* humanized (ordered top to bottom) and chimpanzee ortholog cells (ordered left to right). Blue shading indicates high similarity while red shading indicates low similarity according to the scale at the right (ptShift). The white curve on each heatmap is a dynamic time warping (DTW)-based map between the *HACNS1* humanized and chimpanzee ortholog gene expression profiles compared to an ideal one-to-one mapping across species (shown as a red line). Histograms above and to the left of each heatmap show the proportion of cells from each time point for the chimpanzee ortholog and *HACNS1* humanized lines, respectively. Cells from each time point are labeled according to the legend at the right of the figure. The panel below each heatmap shows pseudotime shift maps obtained by dividing each trajectory into 200 pseudocells (black circles) between each genotype based on gene expression similarity; aligned pseudocells are connected by a line. Regions where pseudocells from the *HACNS1* line align to chimpanzee ortholog pseudocells located later in pseudotime (suggesting accelerated development) are shown in purple while those that align to pseudocells located earlier (suggesting delayed development) are shown in green. The degree of shift is shown based on the scale at the bottom right. Arrows on the heatmaps indicate regions of the pseudotime alignment exhibiting the greatest shifts. See Methods for more details.

In contrast, cells from PA2 in the *HACNS1* humanized line align with cells from the chimpanzee control line that are located at earlier positions along pseudotime, suggesting they are in a less mature transcriptional state and that mesenchymal progenitors in the humanized embryos retain earlier developmental gene expression profiles for a prolonged period (Fig. 4C). Conversely, cells from all three hindlimb lineages, including the autopod, zeugopod, and stylopod lineages in the *HACNS1* humanized line align with cells from the chimpanzee control line that are located at later positions along pseudotime, supporting that *HACNS1* drives accelerated developmental progression in the hindlimb (Fig. 4G-I). In both PA2 and the hindlimb, the largest shifts occur around E10.5, corresponding to the developmental stage at which *HACNS1* shows peak chromatin accessibility, suggesting it is at its highest level of enhancer activity.

To determine the biological processes and pathways implicated in these *HACNS1*-induced heterochronic shifts, we took advantage of the gene modules described above. These modules classify genes with coordinated developmental expression patterns and related biological functions. We used cellAlign^53^ and tradeSeq^54^ to identify genes within the gene modules (Fig. S16-18) that exhibited significant shifts along pseudotime between genotypes (Fig. 5). We found that the majority of shifted genes belong to gene modules that are upregulated toward the end of inferred developmental trajectories, particularly in PA2 and the hindlimb (Fig. 5; Fig. S19-21; Table S5), which display the largest heterochronic shifts between *HACNS1* humanized and chimpanzee ortholog cells. In PA2, genes within one gene module, termed Module 1 in Fig. 5A (see also Fig. S16E, F), showed increased expression toward the end of the pseudotime trajectory in both genotypes, consistent with their association with mesenchymal differentiation. Within this gene module, 71 genes showed significant shifts along pseudotime in the *HACNS1* humanized line, suggesting their expression is delayed compared to the chimpanzee ortholog line (Fig. 5A). To further characterize the temporal patterns of these shifts, we further grouped the shifted genes in Module 1 based on the similarity of their pseudotime shift profiles, which separated them into two clusters (cluster 1 and cluster 2 in Fig. 5E). Notably, all genes in both clusters exhibited delayed increases in expression in the *HACNS1* humanized line relative to the chimpanzee ortholog line (Fig. 5E). In both clusters, the shifts emerged early along the trajectory, reached a maximum at intermediate pseudotime values, and then converged toward comparable levels between genotypes at the end of the trajectory (Fig. 5E, top panel). GO enrichment analysis of the shifted genes revealed enrichment for processes related to canonical Wnt signaling, regulation of cell migration, and cartilage and bone development (Fig. 5B). Genes such as *Sox5*, *Col5a2*, *Mecom*, and *Ror1* showed higher expression earlier in the chimpanzee ortholog line compared to the *HACNS1* humanized line (Fig. 5E). These genes play key roles in early neural crest development and PA morphogenesis, including regulation of cranial neural crest cell (CNCC) development (*Mecom*, *Ror1*)^55,56^, regulation of mesenchymal condensation (*Fn1*)^57^, and control of cartilage development (*Sox5*, *Pcdh7, Col5a2*)^58–60^. Together, the delayed activation of this late-acting, differentiation-associated gene set suggests *HACNS1* drives a delay in chondroblast development in PA2.

**Fig. 5.**
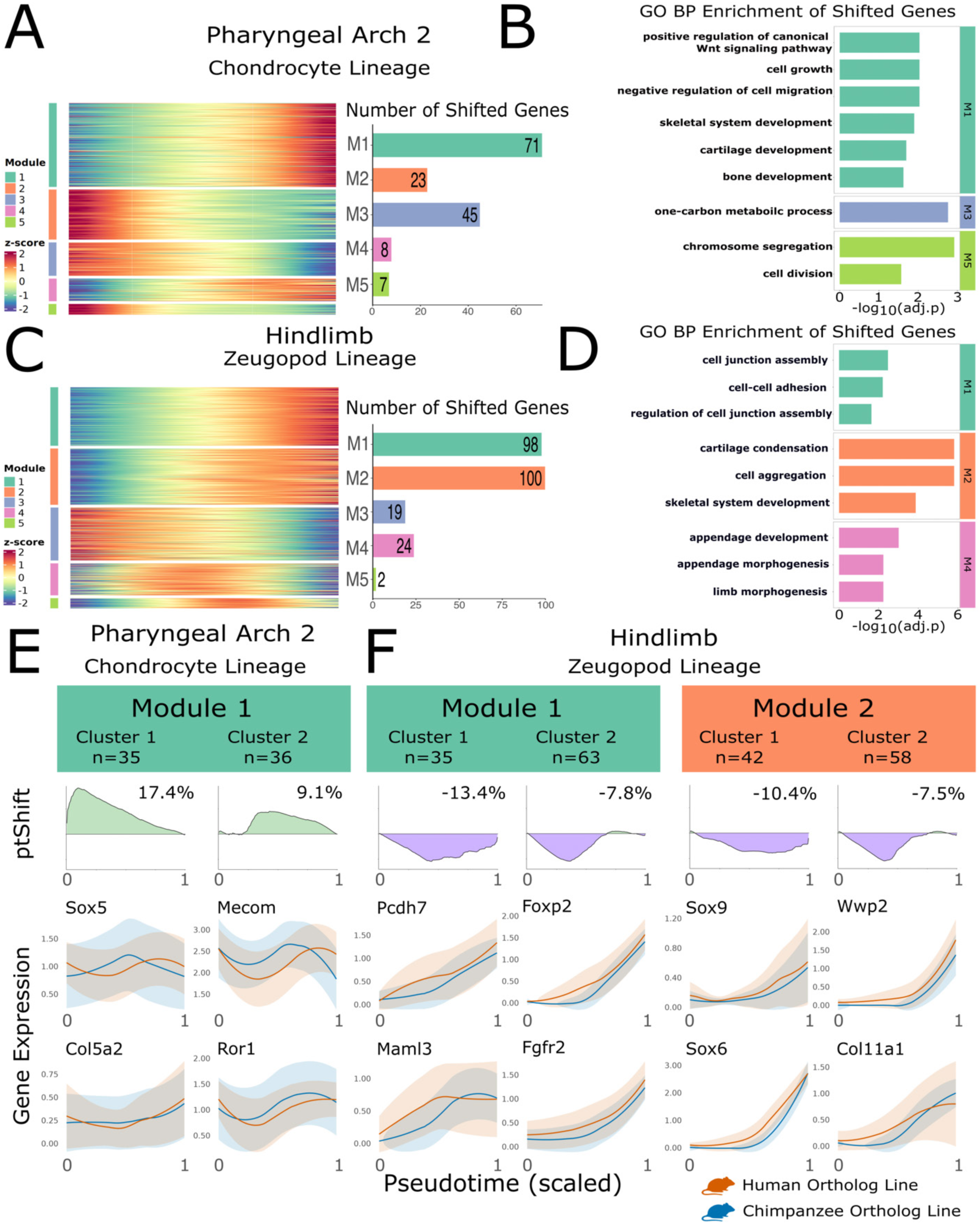
Genes involved in skeletal development show heterochronic shifts in their expression in *HACNS1* humanized mouse pharyngeal arch and hindlimb. A, C: Left panels: Heatmaps showing genes with dynamic expression patterns during development of the PA2 chondrocyte lineage **(A)** or the hindlimb zeugopod lineage **(C)**. Cells are ordered by pseudotime. Gene expression values are scaled using z-scores, with red indicating relatively high expression and blue indicating relatively low expression (Methods). Gene modules are labeled by color throughout the figure according to the legend at the left of each heatmap. Right panels: The number of significantly shifted genes within each module. **B, D:** GO enrichment analysis of shifted genes for each module in the PA2 chondrocyte lineage **(B)** or the hindlimb zeugopod lineage **(D)**. P values were corrected for multiple testing using the Benjamini-Hochberg method. **E, F:** Gene expression shifts along pseudotime for each gene module from the PA2 chondrocyte lineage **(E)** or the hindlimb zeugopod lineage **(F)**. Top panels: Pseudotime shift (ptShift) profiles along the developmental trajectory for each gene cluster within the indicated module. Positive values indicate relative developmental delay (green), whereas negative values indicate relative developmental acceleration (purple) in the *HACNS1* humanized line compared to the chimpanzee ortholog line. The mean pseudotime shift for each cluster is indicated in the top right corner of each plot. Bottom panel: Expression of representative genes from each cluster along each developmental trajectory. Lines depict the average gene expression along pseudotime in each genotype, and shaded areas indicate the standard deviation.

We next focused on the limb bud datasets. In the limb buds, particularly in the hindlimb, the majority of shifted genes belong to gene modules whose expression increases toward later stages of each developmental trajectory (Fig. S20A-C, S21A-C). For example, in the hindlimb zeugopod trajectory, 198 genes showed shifts suggesting they were expressed earlier in the *HACNS1* humanized line compared to the chimpanzee ortholog line (Module 1 and Module 2 in Fig. 5C), which together account for the largest fraction of pseudotime-shifted genes in this trajectory. Both gene modules are enriched in genes involved in mesenchymal cell differentiation and cartilage development (Fig. 5C-D; Fig. S18C, D). We also observed shifts suggesting earlier gene expression in a set of genes showing peak expression in the middle of the developmental trajectory (Module 4 in Fig. 5C). Module 4 includes genes involved in appendage development and limb morphogenesis (Fig. 5D; Fig. S18C, D). In contrast to PA2, the shifted genes within these modules show earlier increases in expression in the *HACNS1* humanized line relative to the chimpanzee ortholog line (Fig. 5F). Similarly, compared with the chimpanzee ortholog line, genes within Module 4 also reach peak expression earlier in the *HACNS1* humanized line (Fig. S21B).

To further characterize the temporal patterns of these shifts, we grouped shifted genes within Module 1 and Module 2 separately based on the similarity of their pseudotime shift profiles as described for the PA2 data above. In each gene module, the shifted genes were partitioned into two clusters. As observed in PA2, the gene expression shifts emerged near the beginning of the trajectory, reached a maximum at intermediate pseudotime values, and converged toward comparable levels by the end of each trajectory (Fig. 5F, top panel). The overall temporal patterns of pseudotime shifts were highly similar across the two modules. GO enrichment analysis revealed that shifted genes in Module 1 were enriched for processes related to cell junction assembly, whereas those in Module 2 were associated with cartilage condensation and skeletal system development (Fig. 5D). Consistent with these functional annotations, genes such as *Pcdh7*, *Maml3*, *Foxp2*, *Fgfr2*, *Sox6*, *Sox9*, *Wwp2*, and *Col11a1* showed earlier increases in expression in *HACNS1* humanized embryos (Fig. 5F). These genes contribute to cartilage differentiation (*Sox6*, *Sox9*)^58,28^, regulation of signaling pathways critical for mesenchymal differentiation and skeletal patterning, including Notch (*Maml3*)^61^, coordination of osteogenesis and chondrocyte maturation (*Foxp2*)^62^, and cartilage development (*Pcdh7*, *Col11a1*, *Wwp2*)^63,59,64^. Together, the earlier upregulation of this late-acting, differentiation-associated gene set is consistent with an acceleration of limb development observed in *HACNS1* humanized embryos.

Although PA2 and the hindlimb showed the strongest heterochronic shifts involving the largest number of shifted genes, we also detected smaller sets of shifted genes in the Md and forelimb (Fig. S19-21; Table S5). In the Md, genes including *Sox5* and *Col5a2* showed earlier expression in the chimpanzee ortholog line and GO enrichment analysis indicated that these shifted genes were associated with cartilage condensation and cell aggregation (Tables S5, S6). In the forelimb, the shifts were more subtle compared to the hindlimb, but genes including *Pcdh7* and *Runx2* still showed earlier expression in the *HACNS1* humanized line (Table S5). These results suggest that *HACNS1* may also affect developmental timing in the Md and forelimb, but with weaker or more restricted effects than those observed in PA2 and the hindlimb.

Together, our results suggest that the heterochronic shifts we observed in both PA2 and the hindlimb mainly reflect changes in the timing of mesenchymal progenitor fate determination: Fate determination is delayed in PA2 but accelerated in the hindlimb. Previous studies^22^ and our snRNA-seq data show that *HACNS1*-driven expansion of *Gbx2* expression occurs primarily in the proximal regions of the pharyngeal arches and the distal regions of the limb buds (Fig. S13), which correspond to early positions along each developmental trajectory. This spatial and temporal pattern suggests that *HACNS1*-mediated *Gbx2* upregulation likely acts early in progenitor populations and subsequently produces downstream effects on skeletal differentiation, even after *Gbx2* itself is no longer expressed.

To determine if the heterochronic shifts in transcription we observed in the Md, PA2, and limb buds corresponded to changes in the timing of cartilage differentiation and development, we performed immunofluorescence (IF) staining for SOX9, a transcription factor essential for cartilage development, in E10.5, E11.5, and E12.5 *HACNS1* humanized mouse embryos, chimpanzee ortholog embryos, and wild-type mouse embryos. At E10.5 and E12.5, we did not observe qualitative differences in the spatial distribution of SOX9 expression between *HACNS1* humanized embryos and the control groups (Fig. S22, 24; Movies S4-12; 22-27). In contrast, at E11.5, SOX9 expression exhibited clear qualitative differences in its spatial organization across the Md, PA2, and limb buds (Fig. 6; Movies S13-21).

**Fig. 6.**
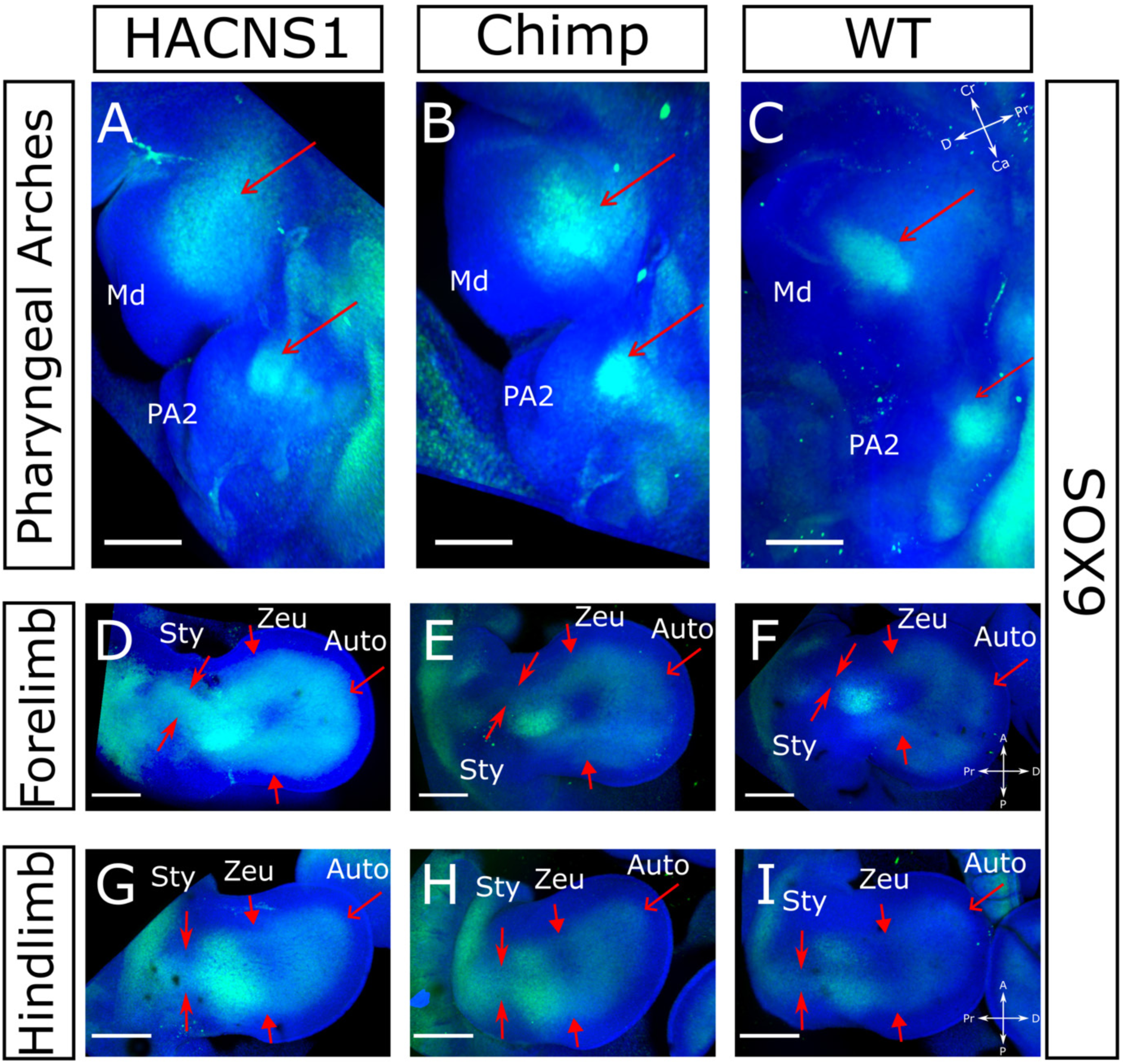
Heterochronic changes in SOX9 expression in *HACNS1* E11.5 humanized mice. Representative immunofluorescence staining for SOX9 in E11.5 *HACNS1* humanized, chimpanzee ortholog, and wild-type control embryos. SOX9 expression domains in the pharyngeal arches **(A-C)**, forelimb **(D-F)**, and hindlimb **(G-I)**. Images were collected from clarified embryos; see Methods for details. Red arrows indicate regions where SOX9 expression qualitatively differs between *HACNS1* humanized embryos and control embryos; the approximate location of each limb segment is also indicated. SOX9 is shown in green, and DAPI nuclear staining is shown in blue; Scale bar, 300 μm; Cr, Cranial; Ca, Caudal; D, Distal; Pr, Proximal; A, Anterior; P, Posterior; Sty, Stylopod; Zeu, Zeugopod; Auto, Autopod; all replicates (n=5) are shown in Fig. S23 and Supplemental Movies S4-S27.

SOX9 expression in *HACNS1* humanized embryos was more diffuse and spatially dispersed in the E11.5 Md and PA2 compared to wild-type embryos, particularly at the proximal end (red arrows in Fig. 6A-C), which represents an earlier stage of mesenchymal development. This finding suggests pre-cartilaginous domains in the Md and PA2 exhibit reduced condensation in the *HACNS1* line (Fig. 6A-C; Fig. S23; Movies S13, 16, 19). Chimpanzee ortholog embryos showed an intermediate pattern, with SOX9 expression appearing less compact than in wild-type embryos but more restricted than in *HACNS1* humanized embryos. Mesenchymal condensation marks the initial step of cartilage formation, during which mesenchymal progenitors aggregate into compact SOX9-positive pre-cartilaginous domains before differentiating into chondrocytes^35,65^. The more diffuse and spatially dispersed SOX9 expression we observed in the Md and PA2 of *HACNS1* humanized embryos suggests delayed condensation of pre-cartilaginous domains compared to wild-type embryos.

During limb development, mesenchymal differentiation proceeds in a proximal-to-distal sequence, with the stylopod differentiating first, followed by the zeugopod and then the autopod. Consistent with this developmental progression, at E11.5, wild-type forelimbs and hindlimbs showed SOX9-positive condensations corresponding to the locations of the stylopod and zeugopod. However, SOX9 expression in the location of the autopod was limited, as the autopod is just beginning to emerge at this time point (Fig. 6D-I). In *HACNS1* humanized embryos, the SOX9-positive condensation domains corresponding to the stylopod and zeugopod (Fig. 6D-I) appeared expanded compared to both chimpanzee ortholog embryos and wild-type embryos. In the autopod, *HACNS1* humanized embryos exhibited a more prominent SOX9-positive condensation domain than wild-type embryos, whereas chimpanzee ortholog embryos showed intermediate expression (Fig. 6D-I). In contrast to the delayed condensation observed in the proximal domain of the Md and PA2, the expansion of SOX9 expression in all three limb segments, particularly in the autopod, suggests cartilage differentiation is accelerated in the limb buds of *HACNS1* humanized embryos.

### Inferring tissue-specific *Gbx2*-associated regulatory networks involved in mesenchymal fate decisions in the pharyngeal arches and limb buds

To gain insight into the regulatory pathways implicated in *HACNS1*-mediated changes in mesenchymal differentiation in the PAs and limb buds, we inferred gene regulatory networks (GRNs) at E10.5 for all tissues in our study (Methods). We focused our GRN analyses on E10.5 for two reasons. First, *HACNS1* exhibits peak chromatin accessibility at this stage, suggesting that it has also reached its highest level of enhancer activity. Second, both our data (Fig. S13) and a previous study^22^ indicate that *Gbx2* expression is localized to undifferentiated mesenchymal cell populations. Although our trajectory alignments showed that *Sox9* and most other shifted genes are upregulated later along the developmental trajectories we identified, and our immunofluorescence data show SOX9 expression shifts after E10.5, our goal was to identify direct downstream GRNs of *Gbx2*. We therefore prioritized E10.5, when both *HACNS1* activity and *Gbx2* expression are highest.

To construct the GRNs, we integrated our ATAC-seq and snRNA-seq datasets with published histone H3K27 acetylation (H3K27ac) ChIP-seq data from E10.5 Md, PA2, forelimb, and hindlimb^66,67^ and promoter capture Hi-C (PCHi-C) data from E10.5 Md and PA2^68^ (Fig. 7A). Briefly, active promoters and enhancers were identified using ATAC-seq and H3K27ac ChIP-seq data, and distal regulatory elements were assigned to target genes using promoter capture Hi-C. Transcription factor binding at these cis-regulatory elements was predicted from ATAC-seq footprinting and TF motif analysis. To reduce false-positive regulatory interactions, we further incorporated snRNA-seq-based co-expression information and retained only transcription factor-target gene pairs supported by both predicted binding evidence and coordinated expression (Methods). The resulting GRNs represent tissue-specific inferred regulatory interactions downstream of *Gbx2,* with genes categorized by their shortest downstream path from *Gbx2* in the network. Direct predicted *Gbx2* targets were defined as one-step downstream genes, whereas more distal network genes were assigned to subsequent steps (Table S7).

**Fig. 7.**
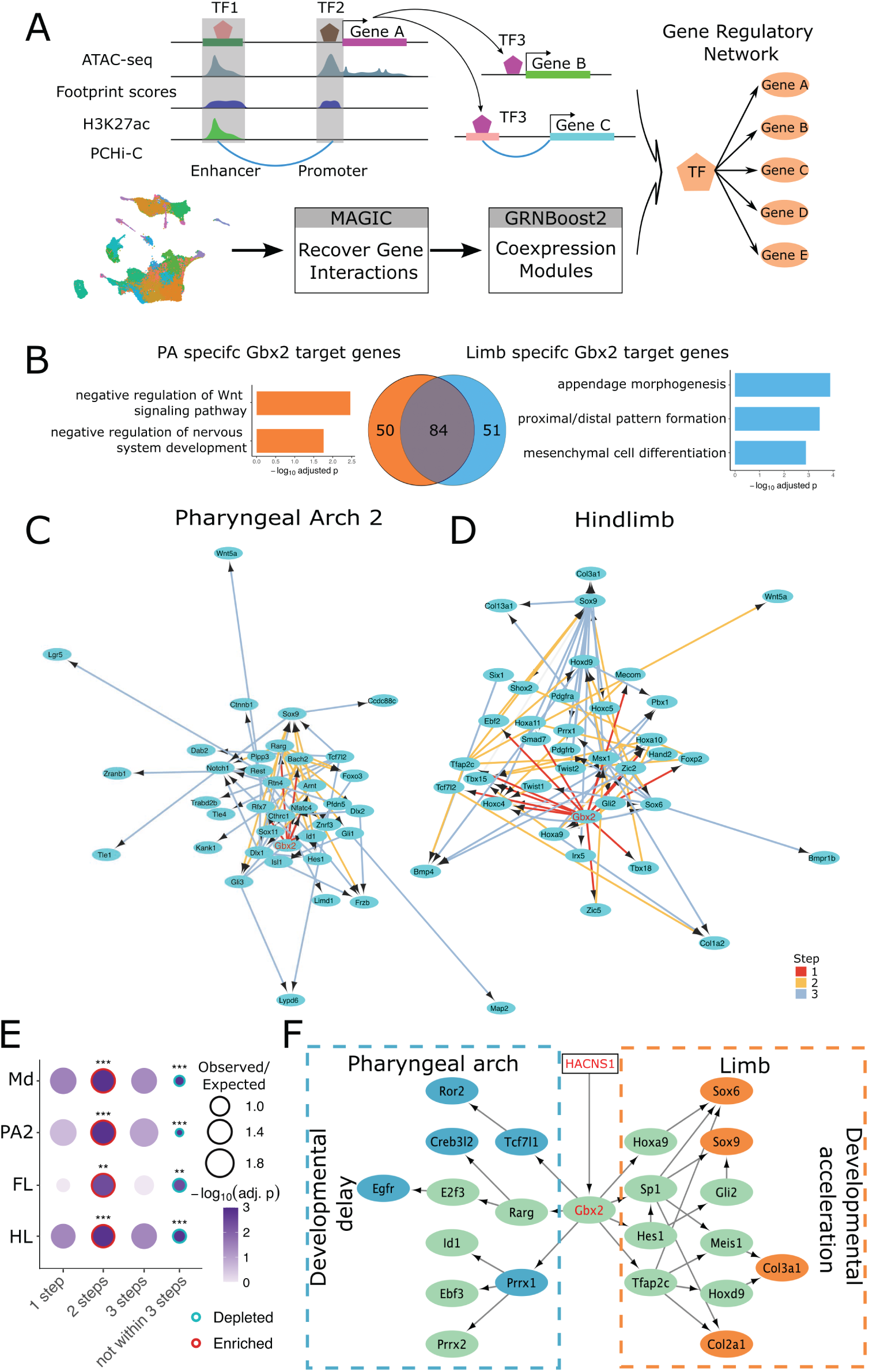
*Gbx2*-driven tissue-specific gene regulatory networks in the pharyngeal arches and limb buds. **A:** Workflow used to reconstruct gene regulatory networks. **B:** Tissue-specific candidate *Gbx2* target genes. Middle panel: Venn diagram shows candidate *Gbx2* target genes identified in pharyngeal arches and limb buds. GO enrichment analysis of pharyngeal arch-specific (Left panel) or limb-specific (Right panel) *Gbx2* target genes. **C, D:** *Gbx2* downstream gene regulatory networks in PA2 **(C)** and hindlimb **(D)**. Edge color indicates the number of steps away from *Gbx2*. Edge length reflects the transcription factor importance score for its target gene, with shorter edges indicating stronger regulatory influence. **E:** Dot plot showing the enrichment of shifted genes by the shortest distance from *Gbx2*. The size of the dots is scaled according to the observed-to-expected ratio calculated from permutation testing (Methods). The color of the dots indicates the adjusted p-value. The color of the dot outline indicates whether shifted genes are enriched or depleted in each step, with red indicating enrichment and cyan indicating depletion. **adjusted p < 0.01; ***adjusted p < 0.001. **F:** Model of *Gbx2-*regulated tissue-specific GRNs controlling mesenchymal cell development in pharyngeal arches and limb buds. Rectangular nodes indicate enhancers, and oval nodes indicate genes. Blue nodes indicate genes whose expression is shifted earlier in chimpanzee ortholog PAs compared to *HACNS1* humanized PAs (suggesting *HACNS1* PA development is delayed), and orange nodes indicate genes shifted earlier in *HACNS1* humanized limb, suggesting *HACNS1* limb development is accelerated. Green nodes represent genes that do not show significant shifts.

We then focused on inferred *Gbx2* target genes in each tissue from tissue-specific GRNs based on ATAC-seq footprint scores at *Gbx2* motif sites. At E10.5, aggregated ATAC-seq footprints across all tissues centered on *Gbx2* motif sites display a characteristic central depletion flanked by accessible chromatin, consistent with putative GBX2 binding (Fig. S25A-D). This pattern is observed in both the *HACNS1* humanized and chimpanzee ortholog lines. However, the depth of the central depletion around the *Gbx2* motif is greater in the *HACNS1* humanized line, suggesting increased GBX2 binding, particularly in PA2 and the hindlimb (Fig. S25E-H).

We next compared inferred *Gbx2* target genes across tissues and identified 50 and 51 tissue-specific *Gbx2* target genes in the PAs and limb buds, respectively (Fig. 7B). GO enrichment analysis revealed that PA-specific *Gbx2* target genes are enriched for negative regulation of Wnt signaling and negative regulation of neurogenesis (Fig. 7B; Table S8). Reconstruction of the GRNs in the Md (Fig. S26A) and PA2 (Fig. 7C) further showed that *Gbx2* may regulate genes involved in Wnt signaling (*Ctnnb1*, *Frzb*, *Wnt5a*) and components of the Notch and Id pathways (*Notch1*, *Id1*). In addition, *Gbx2* is also predicted to directly or indirectly regulate key developmental transcription factors, including *Rarg*, *Dlx1, Dlx2*, *Id1*, *Hes1*, *Rest*, and *Gli1*. Together, these results suggest that *Gbx2* may maintain the stemness of neural crest-derived mesenchymal progenitor cells through the activation of *Id1* and *Notch*-*Hes1* signaling, induction of *Dlx*-dependent craniofacial mesenchymal programs, and negative regulation of Wnt signaling, a regulatory network known to preserve progenitor states and restrain premature differentiation in neural crest-derived lineages^69,70^.

In contrast, limb-specific *Gbx2* target genes are enriched for functions related to limb development, mesenchymal cell differentiation, and skeletal system development (Fig. 7B; Table S8). Reconstruction of the GRNs in the limb buds (Fig. 7D; Fig. S26B) showed that *Gbx2* may directly regulate HOX genes (*Hoxa9*, *Hoxa10*, *Hoxc5*) and limb identity regulators (*Prrx1*, *Tbx15*, *Tbx18*, *Shox2*). *Gbx2* may also indirectly act upstream of key chondrogenic factors such as *Sox9, Bmp4,* and *Col3a1*. Compared with the PAs, the limb network suggests that *Gbx2* is more strongly associated with programs regulating skeletal and appendage morphogenesis.

After reconstructing the GRNs, we next asked whether genes showing heterochronic shifts were included within tissue-specific *Gbx2*-associated networks. For each tissue, we counted the shortest distance from *Gbx2* to each shifted gene and calculated the proportion of shifted genes located one, two, or three steps from *Gbx2* in the network, or that were not within three steps of *Gbx2* (Fig. S25J). Across all tissues, shifted genes were significantly enriched two steps downstream of *Gbx2* and depleted among genes not located within three steps of *Gbx2* (Fig. 7E; Fig. S25J), supporting a connection between *HACNS1*-mediated changes in *Gbx2* expression and downstream shifts in gene expression.

Based on these analyses, we hypothesize that *HACNS1*-mediated changes in *Gbx2* expression act early in mesenchymal progenitor populations and influence the timing of mesenchymal differentiation through tissue-specific regulatory networks (Fig. 7F). In the Md and PA2, *Gbx2* participates in GRNs associated with delayed progression of neural crest-derived mesenchyme toward chondrogenic differentiation. In the limb buds, the *Gbx2*-associated network includes limb patterning and chondrogenic regulators, consistent with accelerated skeletal differentiation.

## Discussion

Among the many characteristics that distinguish humans from other primates, two are especially prominent: speech and bipedal locomotion. The evolution of these derived features depended on modifications to early developmental programs that pattern the PAs and limb buds, leading to human-specific anatomical changes in the face and limbs. One mechanism by which such evolutionary changes can arise is heterochrony, in which shifts in developmental timing alter the progression of tissue patterning and differentiation, ultimately contributing to differences in morphology^71,72,1,73^. The mandible arises from neural crest-derived mesenchyme within the first and second PAs, and limb skeletal elements emerge through tightly regulated developmental programs in the limb buds. Changes in the timing of these processes could have important downstream consequences for human craniofacial and limb evolution.

A fundamental goal in understanding human evolution is to determine how human-specific genetic changes influenced developmental programs, including changes in developmental timing. Human accelerated regions (HARs) are a class of conserved noncoding elements that accumulated human-specific sequence changes after divergence from other primates and are enriched near genes involved in development^10,14^. One such HAR is *HACNS1*, which encodes a transcriptional enhancer exhibiting human-specific activity in the developing Md, PA2, and limb buds^13,22^ and which may have contributed to changes in craniofacial and limb development. In this study, we investigated the impact of *HACNS1* during embryonic development using a genetically humanized mouse model. We characterized the temporal dynamics of *HACNS1* activity and identified *HACNS1*-driven heterochronic shifts in the development of the Md, PA2, and limb buds, providing insight into developmental processes that have been modified during human evolution. Previous studies in cellular models of neurodevelopment have shown that many HARs target conserved sets of orthologous genes between humans and chimpanzees but drive human-specific changes in the expression level and distribution of those targets^17,20^. Our results build on these findings by demonstrating that *HACNS1* not only alters the level of gene expression but the timing of gene expression as well, suggesting that evolutionary changes in developmental timing represent an additional dimension through which HARs may have modified human development.

Prior work suggested that *HACNS1* may have influenced human evolution by altering expression of the nearby gene *Gbx2*^22^. Our findings support this model and further suggest that *Gbx2* has distinct developmental functions across tissues. In the Md and PA2, the predicted *Gbx2* regulatory network suggests *Gbx2* contributes to neural crest fate decisions by promoting mesenchymal rather than neural fate and maintaining the stemness of neural crest-derived mesenchymal progenitor cells (Fig. 7F). In the limb buds, the predicted *Gbx2* network suggests *Gbx2* promotes chondrocyte differentiation (Fig. 7F). Together, these findings support that *HACNS1*-mediated changes in *Gbx2* expression act through distinct tissue-specific downstream regulatory networks, in which a single upstream gene regulatory change may be amplified through gene regulatory network cascades, resulting in opposite developmental effects in craniofacial and limb mesenchyme.

The opposite heterochronic effects of *HACNS1* in craniofacial and limb mesenchyme suggest how *HACNS1* and other HARs could have contributed to the evolution of human craniofacial and limb morphology. Compared to other great apes, humans have a shorter, flatter face and longer hindlimbs associated with bipedal locomotion^74–77^. Delayed developmental progression in the Md and PA2 could have potentially contributed to more flattened craniofacial structures, while accelerated progression in the limb buds, particularly in the hindlimbs, could have contributed to elongated limbs. The relationship between such transient developmental shifts and human-specific morphology remains to be determined. However, our findings suggest that tissue-specific downstream effects of *HACNS1*-mediated *Gbx2* expression changes may provide a mechanism through which one human-specific regulatory change could influence the evolution of multiple anatomical structures in different ways.

Our analyses of SOX9 expression revealed that *HACNS1* induces transient morphological changes during early cartilage development in both the pharyngeal arches and limb buds, with the most dramatic changes observed at E11.5, then returning to near wild-type morphology by E12.5. Although the clearest transcriptomic differences were observed in PA2 and hindlimb, the Md and forelimb also showed more subtle shifts in gene expression, including cartilage-associated genes such as *Sox5* in the Md and *Runx2* in the forelimb. Despite these relatively modest transcriptomic changes, SOX9 staining at E11.5 revealed morphological differences in both the Md and FL, suggesting that even small changes in gene expression may be sufficient to produce observable heterochronic phenotypes. These findings suggest that *HACNS1* drives more restricted transcriptional effects in the Md and forelimb, rather than global shifts observed in PA2 and hindlimb. The weaker transcriptomic effects in the Md and forelimb may arise because increased chromatin accessibility does not necessarily lead to stronger predicted GBX2 binding in either the Md or the forelimb (Fig. S25E, G), nor to stronger downstream effects in these tissues.

Differences in the magnitude and duration of *HACNS1*-driven *Gbx2* upregulation may also contribute to the tissue-specific effects of *HACNS1*. The proportion of *Gbx2*-positive nuclei in the Md was not significantly increased in *HACNS1* humanized embryos, potentially contributing to weaker downstream transcriptional effects compared to PA2, in which *Gbx2*-positive nuclei were significantly increased at E10.5 (Fig. 2E). In the limb buds, both the forelimb and hindlimb showed a significant increase in *Gbx2*-positive nuclei at E10.5, but this increase persisted at E11.5 only in the hindlimb (Fig. 2E). Thus, the more sustained increase in *Gbx2* expression in the hindlimb may contribute to the stronger heterochronic shifts observed in this tissue. In addition, tissue-specific developmental timing may also affect the downstream effects of *HACNS1*. The Md and forelimb initiate development approximately half a day earlier than the PA2 and hindlimb. As a result, peak *HACNS1* chromatin accessibility around E10.5 may occur in different developmental contexts across tissues. Together, these factors could explain why the strongest global transcriptional shifts were observed in PA2 and the hindlimb.

Our results, together with previous work^22^, indicate that the chimpanzee ortholog of *HACNS1* exhibits weaker enhancer activity than human *HACNS1* but may retain stronger activity than the mouse ortholog. This is consistent with intermediate levels of *Gbx2* expression observed in both previous *in situ* hybridization data in chimpanzee ortholog embryos^22^ and our snRNA-seq analysis. Qualitatively, chimpanzee ortholog embryos also show intermediate morphological changes compared with *HACNS1* humanized and wild-type mouse embryos. These observations suggest that the activity of *HACNS1’s* non-human primate ancestral sequences may have been modified earlier in primate evolution. This further raises the possibility that *HACNS1* might genetically interact with regulatory elements that have primate-specific functions. Indeed, previous studies have shown that unique substitutions in both the human lineage and ancestral lineages within the ape clade, as well as lineage-specific accelerated regions, contribute to the development of unique human traits and to human evolution^78–80^.

A previous study did not detect major changes in skeletal morphology in *HACNS1* humanized mice at E18.5^22^. This may be due to the mechanisms described above. However, there is another possibility which should be considered when using humanized mouse models to identify morphological phenotypes. The development of individual skeletal structures occurs in tandem with other anatomically and functionally related structures. Although the upper and lower jaws have distinct developmental programs, their growth and patterning must remain coordinated to produce functional jaws^81^. In the limb, the size, shape, and arrangement of skeletal elements are coordinated with other components of the musculoskeletal system^82^. The evolution of human-specific morphological traits likely required many lineage-specific regulatory changes that altered these functionally related tissues, a hypothesis supported by several recent studies^83–86^. A transient alteration in the development of an individual structure would be insufficient to produce a morphological change without corresponding changes in related structures, driven by other human-specific genetic changes.

The polygenicity of human skeletal evolution coupled with the limited developmental impact of *HACNS1* alone also explains why no major anatomical changes appeared in our model. In Md, PA2, and limb buds, chondrocyte differentiation remains at the earliest phase of chondrogenesis, mesenchymal condensation, at E11.5^46^. During this phase, chondroprogenitors continue proliferating to establish the shape and size of future cartilage. The subsequent differentiation phase begins after condensation and starts around E12.5 in mouse embryos, at which point the earliest cartilage structures become detectable^46^. However, by this stage *HACNS1* activity has already declined. The lack of a subsequent morphological phenotype is consistent with this temporal pattern, suggesting that *HACNS1* induces transient changes in early mesenchymal differentiation that revert to wild-type development as cartilage differentiation proceeds. Persistent morphological differences may therefore require additional lineage-specific regulatory changes acting downstream of *HACNS1*-mediated *Gbx2* regulation. These changes may act at later developmental stages, after *HACNS1* is no longer functioning, to further shape human jaw and limb morphology. We identified tissue-specific downstream regulatory networks associated with *Gbx2*, raising the possibility that the regulation of genes within these networks may also have been modified during human or primate evolution. Such modifications could extend the transient effects of *HACNS1*-mediated changes in *Gbx2* expression, allowing further heterochronic shifts in mesenchymal differentiation that contribute to morphological differences. Identifying such elements will require *in vivo* studies using mouse models of additional human-specific regulatory innovations.

## Methods

### Mouse breeding and embryo collection

All animal work was performed in accordance with approved Yale Institutional Animal Care and Use Committee (IACUC) protocols (#2025-11167). *HACNS1* humanized mouse lines and chimpanzee ortholog lines were generated previously^22^ and have been maintained on a C57BL/6 genetic background. Mouse and embryo genotyping was performed using a PCR-based approach on genomic DNA extracted from toe or yolk sac tissue. DNA was isolated using the DNeasy Blood and Tissue Kit (QIAGEN, #69506) as described previously^22^. Embryonic age was determined by the day on which a vaginal plug was detected and was designated as embryonic day 0.5 (E0.5).

### Sample and library preparation for assays for transposase-accessible chromatin using high-throughput sequencing (ATAC-seq)

Embryos were collected at E9.5, E10.5, E11.5, and E12.5, and the mandibular process, PA2, forelimb, and hindlimb were dissected in ice-cold 1X PBS. Dissected tissues were blotted dry, flash-frozen, and stored in liquid nitrogen until library generation. For each genotype, tissue, and developmental stage, at least eight embryos from two litters were used to generate libraries. For each developmental stage and genotype, the corresponding tissues were collected from the same set of embryos, and tissues of the same type from multiple embryos were pooled to generate each sequencing library.

ATAC-seq libraries and sample indexing were prepared using the ATAC-Seq Kit (Active Motif, #53150) according to the manufacturer’s instructions. For each tissue and developmental stage, two biological replicate libraries were generated. Paired-end sequencing (2 × 100 bp) was performed on an Illumina NovaSeq 6000 platform (RRID: SCR_016387) by the Yale Center for Genome Analysis.

### Sample and library preparation for single-nucleus RNA sequencing (snRNA-seq)

Samples for snRNA-seq were collected following the same procedures described for ATAC-seq. Nuclei were isolated using the 10x Genomics Nuclei Isolation Kit with RNase Inhibitor (PN-1000494) according to the manufacturer’s instructions. Nuclei integrity, debris contamination, and nuclei concentration were assessed using ReadyCount Green/Red Viability Stain (acridine orange/propidium iodide; Thermo Fisher Scientific, A49905) and a fluorescent automated cell counter (CellDrop FL S-09982).

Nuclei suspensions were loaded into PIP tubes (FB0003914) for single-nucleus capture and barcoding, followed by library preparation and sample indexing using the PIPseq™ T20 3′ Single Cell RNA Kit v4.0 PLUS Kit (Fluent BioSciences, #FBS-SCR-T20-4-V4.05-1). Barcoded snRNA-seq libraries from the same developmental stage and tissue were pooled into a single sequencing lane and sequenced on the Illumina NovaSeq 6000 platform (RRID: SCR_016387).

Approximately 20,000 reads per input nucleus were generated for each library. Sequencing was performed by the Yale Center for Genome Analysis.

### Clear lipid-exchanged acrylamide-hybridized rigid imaging compatible tissue hydrogel (CLARITY)

A simplified CLARITY protocol was used to clear fixed embryos for further visualization^87^. Briefly, embryos were collected at E10.5, E11.5, and E12.5 and fixed in 4% paraformaldehyde for one week. Following fixation, embryos were dehydrated and incubated in 100% methanol at -20 °C for at least one week. Samples were then bleached overnight at room temperature in bleach solution (80% methanol, 10% hydrogen peroxide, and 10% DMSO) to reduce pigmentation-related artifacts. After bleaching, embryos were rehydrated and cleared by incubation in 8% SDS in 1× PBS at 37 °C for 4 days (E10.5 embryos) or 5 days (E11.5 and E12.5 embryos). Detergent was removed through multiple 1X PBS washes, and samples were then used for subsequent immunofluorescence staining and imaging.

### Immunofluorescence staining and imaging

Optically cleared embryos were blocked overnight at room temperature in 1× phosphate-buffered saline containing Tween-20 (PBST), 10% fetal bovine serum (FBS), and 0.01% (w/v) sodium azide. Embryos were then incubated with the primary antibody diluted in blocking solution at room temperature for 4 days. The primary antibody used was anti-SOX9 (Millipore Sigma-Aldrich, #AB5535) at a dilution of 1:400. Samples were washed three times with PBS, followed by an overnight wash.

Embryos were subsequently incubated at room temperature with Alexa Fluor™ 488-conjugated secondary antibody (1:500; Thermo Fisher Scientific, #A-11008) together with DAPI (1:1000; Thermo Fisher Scientific, #62248) for 4 days. After three PBS washes and an additional overnight wash, embryos were transferred into RIMS buffer (1× PBS containing 88% w/v Histodenz and 0.01% w/v sodium azide) until optically transparent. Stained embryos were embedded in 1% low-melting-point agarose prepared in RIMS buffer for imaging. Imaging was performed with a Nikon CSU-W1 SoRa spinning-disk confocal microscope with laser powers of 5% for DAPI and 10% for Alexa Fluor™ 488, exposure times of 50 ms for DAPI and 100 ms for Alexa Fluor™ 488, and a z-step size of 2.5 μm, and three-dimensional images were generated with Imaris 10.1.

### ATAC-seq data analysis

Reads were quality checked with FastQC (v0.12.1) using default settings^88^. Adapter sequences (CTGTCTCTTATACACATCT) were trimmed from both reads using cutadapt (v5.0)^89^. Bowtie2 (v2.5.1) indices were generated by replacing the corresponding mouse genomic (mm10) locus with the human *HACNS1* sequence or the chimpanzee ortholog sequence, the sequences of which were described previously^22^. ATAC-seq reads were aligned to the appropriate genome using Bowtie2^90^ in local alignment mode with high-sensitivity settings (--local --very-sensitive --no-mixed --no-discordant --phred33 -I 10 -X 700). Aligned reads were sorted and indexed using SAMtools^91^ (1.21-GCC-13.3.0). Mitochondrial reads were removed using samtools view, and duplicate reads were removed using Picard^92^ MarkDuplicates (v2.25.6; REMOVE_DUPLICATES=true). Reads with mapping quality scores below 10 were subsequently excluded using samtools view (-q 10). All peak calling was performed using MACS2 (v2.2.9.1) ^93^ with a false discovery rate threshold of q < 0.1 and without duplicate filtering (--keep-dup all), and peaks were called as narrow peaks. Subsequently, differential analysis was performed on these peaks using DiffBind with the default DESeq2^31,32^ pipeline (DESeq2::nbinomWaldTest), with a false discovery rate < 0.05.

### snRNA-seq read alignment

Barcode whitelists for each dataset were generated using pipseeker^94^ (v2.1.4) with the barcode function (--chemistry v4) and were subsequently used for read alignment. Reads from all single-nucleus RNA-sequencing experiments were aligned to the mouse reference genome (mm10) using STAR^95^ (v2.7.11a) (parameters: --outSAMtype BAM Unsorted --outSAMattributes NH HI AS nM jM jI CR CY UR UY GX GN --outSAMprimaryFlag AllBestScore --outSAMmultNmax 10 --outBAMcompression 10). STARsolo^96^ was used to generate count matrices for full-length gene, splice junction, and velocyto features and select cells for downstream analysis (implemented in STAR v2.7.11a; parameters: --soloFeatures Gene GeneFull SJ Velocyto --soloType CB_UMI_Simple --soloCBwhitelist barcode_whitelist.txt --soloCBstart 1 --soloCBlen 16 --soloUMIstart 17 --soloUMIlen 12 --soloBarcodeReadLength 1 --soloCBmatchWLtype 1MM_multi --soloUMIdedup 1MM_All --soloUMIfiltering ---soloCellFilter CellRanger2.2). Specifically, both pre-mRNA and spliced mRNA transcripts were captured for single-nucleus RNA sequencing by counting reads aligned across the entire gene body, including intronic, exonic, and exon-exon junction regions, using the --soloFeatures GeneFull option.

### Quality control, cell filtering, normalization, and cluster analysis

snRNA-seq data were initially processed using Seurat^97^ v5.1.0. We evaluated standard quality control metrics, including total feature counts (nCount_RNA), the number of detected genes (nFeature_RNA), and the percentage of reads derived from mitochondrial RNA (percent.mt). Because single-nucleus RNA-sequencing is expected to contain minimal mitochondrial transcripts, nuclei with greater than 1% mitochondrial content were excluded. Nuclei with fewer than 200 detected genes or more than 5,000 detected genes were also removed.

For initial preprocessing, expression values from each sample were normalized based on library size using the NormalizeData function (normalization.method = "LogNormalize", scale.factor = 10000). To reduce variation due to cell cycle effects while preserving differences between cycling and non-cycling cells, cell cycle scores were calculated using the CellCycleScoring function in Seurat, and the difference between G2M and S phase scores was regressed out. To further minimize technical artifacts caused by doublets, doublets were identified and removed using DoubletFinder^98^ with default parameters (PCs = 1:30, pN = 0.25, pK = 0.07-0.29). Finally, the data were normalized using SCTransform, with mitochondrial content regressed out (vars.to.regress = "percent.mt").

Next, samples from the same tissue and developmental stage were integrated across genotypes using the RunHarmony function in Seurat v5 with default parameters (group.by.vars = "genotype", project.dim = FALSE). For each integrated dataset, principal component analysis (PCA) was performed. Nuclei were then embedded using uniform manifold approximation and projection (UMAP) by applying the RunPCA function (npcs = 50, verbose = FALSE) followed by the RunUMAP function (reduction = "harmony", dims = 1:30, verbose = FALSE) implemented in Seurat.

We then identified nuclei clusters using the Louvain clustering algorithm as implemented in Seurat. Briefly, a shared nearest neighbor graph was constructed in PCA space using the FindNeighbors function (reduction = "pca", dims = 1:30, verbose = FALSE), followed by application of the Louvain algorithm to define clusters using the FindClusters function (resolution = 0.5). Marker genes for each cluster were identified using the FindAllMarkers function (logfc.threshold = 0.25, min.pct = 0.1, min.diff.pct = -Inf, only.pos = TRUE).

Finally, differences in the proportion of *Gbx2*-positive nuclei between genotypes were assessed using two-tailed Fisher’s exact test. P values were adjusted for multiple testing using the Benjamini-Hochberg method.

### Integration and PHATE embedding

Mesenchymal nuclei from all developmental stages within each tissue were integrated using the RunHarmony function in Seurat v5 (group.by.vars = "stage", project.dim = FALSE). To minimize potential artifacts arising from dissection variability, mesenchymal nuclei from the most proximal regions of the limb buds, including the trunk cluster and the scapula/iliac blade cluster, were excluded from downstream analyses. The integrated mesenchymal cells were then visualized using potential of heat-diffusion affinity-based transition embedding (PHATE)^41^ to capture global structure and developmental patterns (PAs: knn = 3, n_pca=200, decay = 15, t = 35, gamma = 0; limb buds: knn = 3, n_pca=100, decay = 15, t = 35, gamma = 0). The embeddings were then visualized in three dimensions using the phate.plot.rotate_scatter3d function. Finally, the initial clustering results were merged into broader cell type clusters for better visualization purposes.

### Reconstructing developmental trajectories

To identify developmental trajectories within the integrated mesenchymal cell populations for each tissue, we first applied URD to the data^43^. Briefly, a diffusion map was constructed, and transition probabilities between cells were calculated using the calcDM function (knn = 200, sigma.use = 8). The resulting transitions were visualized and evaluated by plotting multiple pairs of diffusion map dimensions, as well as by projecting them onto a t-SNE embedding using the plotDim function. Pseudotime scores were then calculated using the floodPseudotime function (n = 150, minimum.cells.flooded = 2) and the floodPseudotimeProcess function. These pseudotime scores were used only for biased random walks. For the PAs, root cells were defined as the proximal mesenchymal populations from either the mandibular process or PA2 at E9.5. For the forelimb, root cells were defined as the progress zone population from the E9.5 sample, whereas for the hindlimb, root cells were defined as the progress zone population from the E10.5 sample. Terminal cell populations were defined as early chondroblast or early osteoblast populations from E12.5 for the PAs, and as chondroprogenitor populations from E12.5 for both the forelimb and hindlimb. Biased random walks were then performed 25000 times for each terminal tip using the simulateRandomWalksFromTips function (transition.matrix = axial.biased.tm, n.per.tip = 25000, root.visits = 1, max.steps = 5000, verbose = FALSE). The resulting random walks were subsequently processed into visitation frequencies using the processRandomWalksFromTips function.

Because the biased random walks alone were insufficient to resolve the autopod, zeugopod, and stylopod lineages in the limb buds or to distinguish chondroblast and osteoblast lineages in the PAs, we further subset cells with visitation frequencies greater than 1. These cells were then embedded using two-dimensional PHATE (knn = 3, n_pca = 50, decay = 50, t = 50, gamma = 0). This additional embedding step enabled clear separation of the corresponding lineages. Lineage trajectories and pseudotime were then inferred using pyslingshot^44^ on the two-dimensional PHATE embedding with default parameters. Because PA2 contributes only a small portion of the mandibular region at E12.5, trajectories for PA2 were constructed using samples from E9.5 to E11.5 only.

### Identification of genes with trajectory-dependent expression changes

To identify genes with correlated expression patterns along developmental trajectories, we used a spatial autocorrelation test using the graph_test function in Monocle3^99^. Genes with q-values < 0.05 were retained, and the filtered genes were subsequently clustered into gene modules using the find_gene_modules function with Louvain clustering. A resolution of 0.01 was used, resulting in five to six gene modules for each lineage.

To illustrate dynamic gene expression patterns within each module, we calculated z-scored expression values for each gene and visualized them using the pheatmap package^100^. For individual genes, we fitted smooth curves relating average gene expression to pseudotime using locally weighted regression (LOESS). The square root of the resulting local variance estimates was used to compute standard deviation envelopes around the mean expression trajectory.

### Alignment of developmental trajectories

To quantify developmental trajectories and compare expression dynamics across genotypes, we performed dynamic time warping (DTW) of pseudotime trajectories using CellAlign^53^. Trajectories for each genotype were interpolated to 200 equally spaced points along pseudotime. Global alignment was performed using the globalAlign function to align the entire trajectories of both genotypes. Pairwise Euclidean distances between ordered points were computed using the shared 3,000 highly variable genes. Gaussian weighting (winSz = 0.1) was applied to reduce noise and smooth the alignment.

### Trajectory-dependent differential gene expression analysis

To identify genotype-specific transcriptional differences along developmental trajectories, we used tradeSeq to fit generalized additive models (GAMs) for each trajectory^54^. Raw UMI count matrices for genes within each module were used as input to the fitGAM function. Genotype-dependent differential expression along pseudotime was evaluated with a condition-based test using the conditionTest function, and genes with p-values ≤ 0.05 were retained as significantly differentially expressed.

To group genes exhibiting similar temporal expression shift patterns between genotypes, we performed pseudotime-based clustering on significantly differentially expressed genes for each trajectory using the pseudotimeClust function in CellAlign. Gene clusters were further filtered based on the magnitude of the pseudotime shift (ptShift) and whether the shift occurred after gene activation, as differences detected at low expression levels before gene activation are more likely to reflect noise than true biological differences.

### Gene ontology enrichment analysis

Gene Ontology (GO) enrichment analysis was performed for all gene sets analyzed in this study using the enrichGO function from the clusterProfiler package^101^ with the mouse annotation database (org.Mm.eg.db). For each analysis, the background gene set was defined as all genes detected in the corresponding snRNA-seq dataset. Enrichment was assessed across all GO ontologies, including Biological Process, Molecular Function, and Cellular Component (ont = "ALL"). Multiple testing correction was performed using the Benjamini-Hochberg method, and GO terms with adjusted q-values ≤ 0.05 (pAdjustMethod = "BH", qvalueCutoff = 0.05) were considered significant.

### Footprinting analysis

To quantify differences in transcription factor binding between genotypes, we re-mapped E10.5 ATAC-seq data from all tissues to the mouse reference genome (mm10) and analyzed the data using the TOBIAS framework^102^. For each dataset, BAM files were first processed with TOBIAS-ATACorrect to correct for Tn5 transposase insertion bias, generating corrected bigWig files representing the ATAC-seq signal. For each tissue, a merged set of peaks was generated, and this consensus peak set was used as input to TOBIAS-ScoreBigwig to compute base-pair footprint scores across the corresponding genomic regions. TOBIAS-BINDetect was then applied to identify motif occurrences within the merged peak set and to calculate associated footprint scores for each genotype using vertebrate transcription factor motifs from the JASPAR2020 database. This step also quantified genotype-specific differences in putative transcription factor binding across the shared peak set. Finally, TOBIAS-PlotAggregate was used to generate aggregate footprint profiles by averaging corrected ATAC-seq signal within ±100 bp windows surrounding motif occurrences.

### Modeling tissue-specific gene regulatory networks

We used the combined pool of binding sites generated from footprinting analysis to unravel transcription factor binding networks. First, ATAC-seq peaks at promoter regions were annotated with respect to transcription start sites using ChIPseeker^103^, with promoters defined as regions spanning 2 kb upstream to 500 bp downstream of transcription start sites.

Then, in order to identify active enhancers in each tissue at E10.5, we analyzed published H3K27ac ChIP-seq datasets^66,67^. FASTQ files were downloaded from the Sequence Read Archive for the mandibular process (SRR4835044), PA2 (SRR4835045), forelimb (SRR6294689, SRR6294690), and hindlimb (SRR3950341, SRR3950342). Reads were aligned to the mouse reference genome (mm10) using Bowtie2 (--local --very-sensitive --no-mixed --no-discordant --phred33 -I 10 -X 700). Aligned reads were sorted, duplicate reads were removed, and mitochondrial reads were excluded before peak calling. Narrow peaks were called using MACS2 (--q 0.1 --keep-dup all). Active enhancers were defined by intersecting H3K27ac peaks with ATAC-seq-defined open chromatin regions using the intersect function in BEDTools^104^ (2.31.1-GCC-13.3.0).

To identify target genes of active enhancers, we used published high-confidence enhancer-promoter interactions from the E10.5 mandibular process and PA2, identified using PCHi-C^68^. ATAC-seq peaks at active enhancer regions were subsequently annotated based on their associated target genes. Because PCHi-C data are not available for the limb buds, enhancer-promoter interactions identified in the PAs were used for enhancer-target gene annotation in the limb buds. Next, annotated ATAC-seq peaks and transcription factor binding files were used to construct transcription factor binding networks for each tissue using the CreateNetwork function in TOBIAS, resulting in tissue-specific direct binding networks.

To evaluate transcription factor binding activity, we inferred coexpression relationships between transcription factors and candidate target genes using our snRNA-seq dataset. Because snRNA-seq data often contain many sources of technical noise, such as mRNA undersampling, which can obscure gene-gene relationships, we first denoised gene expression values using MAGIC^105^ with default parameters. Coexpression modules were then inferred using GRNBoost2, which was implemented in pySCENIC^106^. Because GRNBoost2 is computationally intensive when applied to the whole datasets and because the GRN step is stochastic, snRNA-seq datasets were randomly downsampled to 3,000 cells per dataset using NumPy^107^ (np.random.choice). This procedure was repeated 40 times, and target genes identified in more than 95% of runs were considered high-confidence coexpression targets. Tissue-specific transcription factor direct binding networks were then filtered using coexpression modules to retain only those interactions in which the target gene was directly bound by the transcription factor and the transcription factor exerted a positive regulatory contribution to the target gene’s expression. Finally, gene regulatory networks were visualized using Cytoscape with edge-weighted force-directed layouts, in which edge lengths reflect the average importance score of each transcription factor-target interaction across the 40 runs.

To test whether shifted genes were enriched at specific distances from *Gbx2* within the inferred GRNs, all genes tested for pseudotime shifts were used as the background. GRNs were converted into directed graphs in R using the igraph package^108^, with edges oriented from transcription factors to target genes. For each tissue, the shortest downstream path length from *Gbx2* to each gene was calculated using igraph::distances with mode = "out". Genes were then assigned to one of four categories: one step, two steps, three steps, or not within three steps of *Gbx2*. For each tissue, we performed two-tailed permutation testing. We randomly sampled the same number of genes as the observed shifted gene set from the corresponding background gene set 20,000 times. For each permutation, we counted the number of sampled genes assigned to each *Gbx2* step, generating an expected distribution for each step. The observed number of shifted genes in each step was then compared with this expected distribution. Enrichment or depletion was quantified as the ratio of the observed count to the mean expected count across permutations. The empirical two-sided p-values were calculated by doubling the smaller of the proportions of trials greater than or less than the observed number of shifted genes, followed by Benjamini-Hochberg correction.

## Acknowledgements

We thank S. Mane, C. Castaldi and colleagues at the Yale Center for Genome Analysis for generating the sequencing data used in this study, S. Wilson and colleagues in the Yale Neuroscience Imaging Core Facility for assistance with microscopy, the Yale Animal Resources Center for assistance with mouse husbandry, and the members of the Noonan laboratory for their input on the manuscript. This work was supported by an award from the Eunice Kennedy Shriver National Institute of Child Health and Human Development (NICHD; R01 HD102030 to J.P.N), an NSF Graduate Research Fellowship (to M.M.), an NICHD F32 Postdoctoral Fellowship (F32 HD108935 to M.B) and funds from the Yale School of Medicine. This research program and related results were also made possible by the support of the NOMIS Foundation (to J.P.N.).

## Author Contributions

Y.J. and J.P.N. conceived and designed the study. Y.J. designed and performed ATAC-seq and snRNA-seq experiments, SOX9 whole mount immunofluorescence staining, animal husbandry, and embryo collection, staging, and dissections. Y.J. also carried out all computational analyses with advice from M.M. M.B. provided technical input on the preparation of PIPseq snRNA-seq libraries. Y.J. and J.P.N. wrote the manuscript with input from all authors.

## Data and Code Availability

ATAC-seq sequencing data and peak calls are available from the Gene Expression Omnibus (GEO) under accession number GSE343875. PIPseq snRNA-seq reads, barcodes and count matrices are available at GEO under accession number GSE343880. Imaging data are available through Mendeley at doi.org/10.17632/8mjmc2s3h6.1. Code associated with the analyses performed in this paper is available at GitHub: https://github.com/NoonanLab/Ji_et_al_HACNS1 and Zenodo: doi.org/10.5281/zenodo.22348376.

## Supplemental Figures

**Fig. S1.**
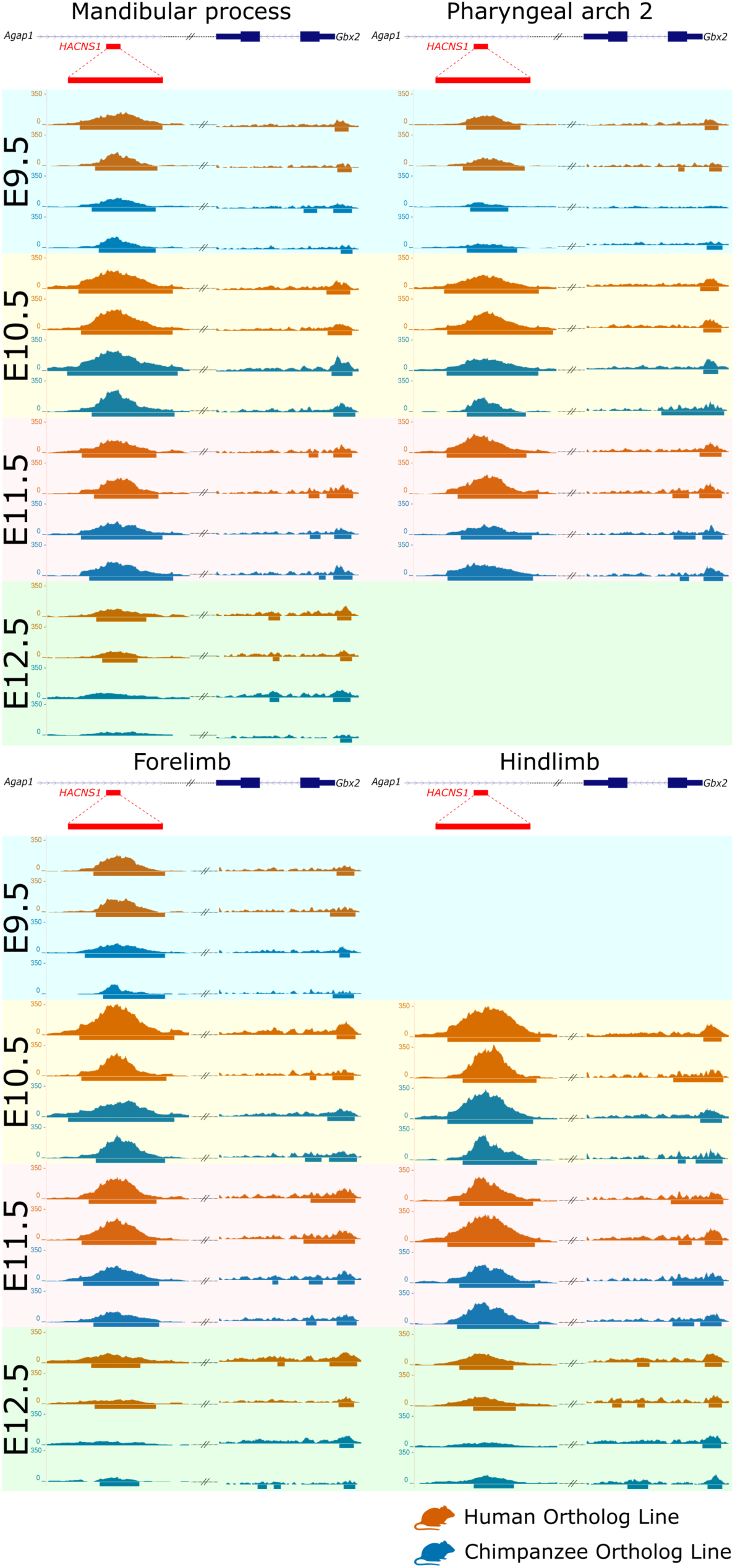
Chromatin accessibility at the *HACNS1* locus and the *Gbx2* promoter. ATAC-seq signal at the *HACNS1* locus (left) and the *Gbx2* promoter (right) for the *HACNS1* humanized line (dark orange) and the chimpanzee ortholog line (blue) in the mandibular process, second pharyngeal arch, forelimb, and hindlimb at E9.5 (light cyan), E10.5 (light yellow), E11.5 (light pink), and E12.5 (light green). Two independent biological replicates are shown for each genotype, tissue, and developmental stage. The positions of the humanized *HACNS1* locus (in red) and the *Gbx2* gene (in blue) in the *HACNS1* line are indicated above the tracks. Peak calls indicating significantly accessible regions are shown as bars below each signal track. See Methods for details on the genome assembly used for alignment and peak calling.

**Fig. S2.**
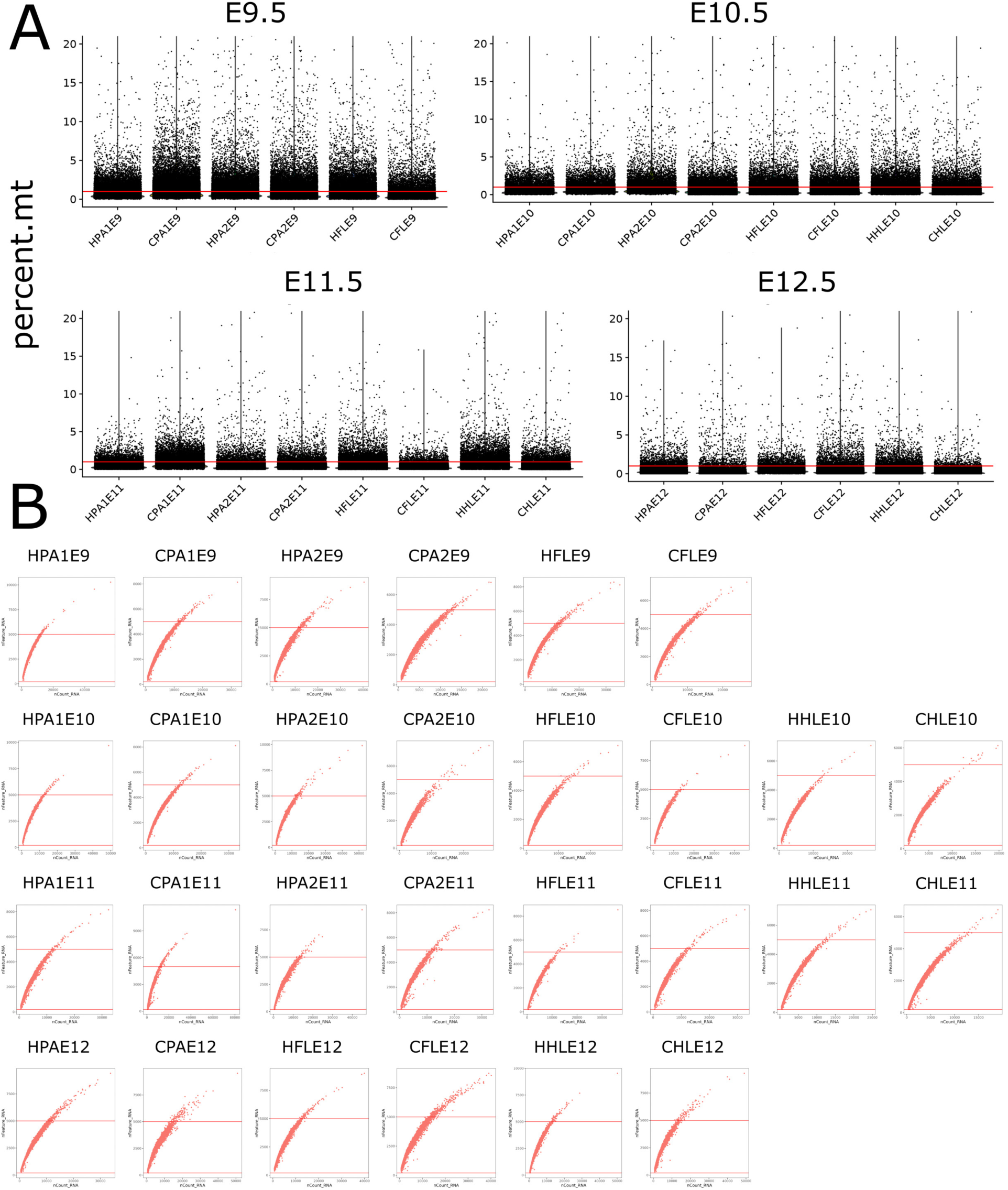
Quality control filtering of single-nucleus RNA-seq data. **A:** The percentage of mitochondrial RNA (percent.mt, vertical axis) per nucleus is shown for each indicated sample (horizontal axis) from each time point. Red lines indicate the 1% threshold used for filtering. **B:** Total RNA counts per nucleus (nCount_RNA, horizontal axis) and the number of genes detected per nucleus (nFeature_RNA, vertical axis) for each sample. Red lines indicate the lower and upper nFeature_RNA thresholds of 200 and 5,000, respectively, used for filtering. Sample IDs indicate genotype, tissue, and developmental stage: H, *HACNS1* humanized line; C, chimpanzee ortholog line; PA1, mandibular process; PA2, second pharyngeal arch; FL, forelimb; HL, hindlimb; and E9-E12, E9.5-E12.5.

**Fig. S3.**
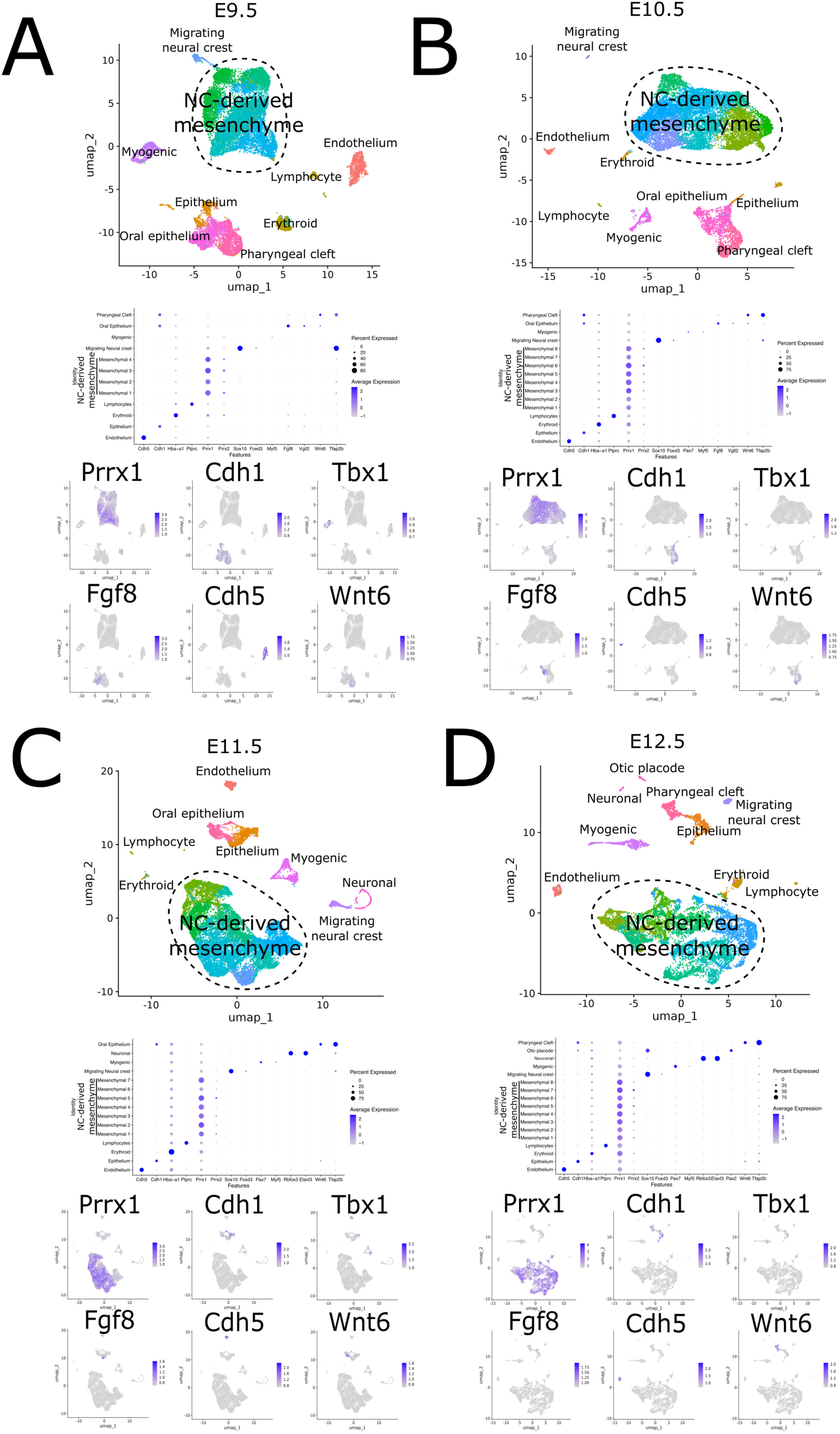
Cell types identified in the mandibular process from snRNA-seq data. Major cell types identified in integrated mandibular process snRNA-seq datasets from *HACNS1* humanized and chimpanzee ortholog embryos at E9.5 **(A)**, E10.5 **(B)**, E11.5 **(C)**, and E12.5 **(D)**. The top panels for each main figure panel show UMAP embeddings colored by cell type assignments. Neural crest-derived mesenchymal populations are outlined by dashed lines. The middle panels show dot plots illustrating the expression of marker genes across annotated cell types. The dot size represents the percentage of nuclei expressing each gene, and the color intensity represents the average scaled expression of the gene within each cell type (see the scales to the right of each panel). The bottom panels show normalized expression of representative cell type marker genes in UMAP embedding, with increasing color intensity indicating higher expression (see scale bars to the right of each panel).

**Fig. S4.**
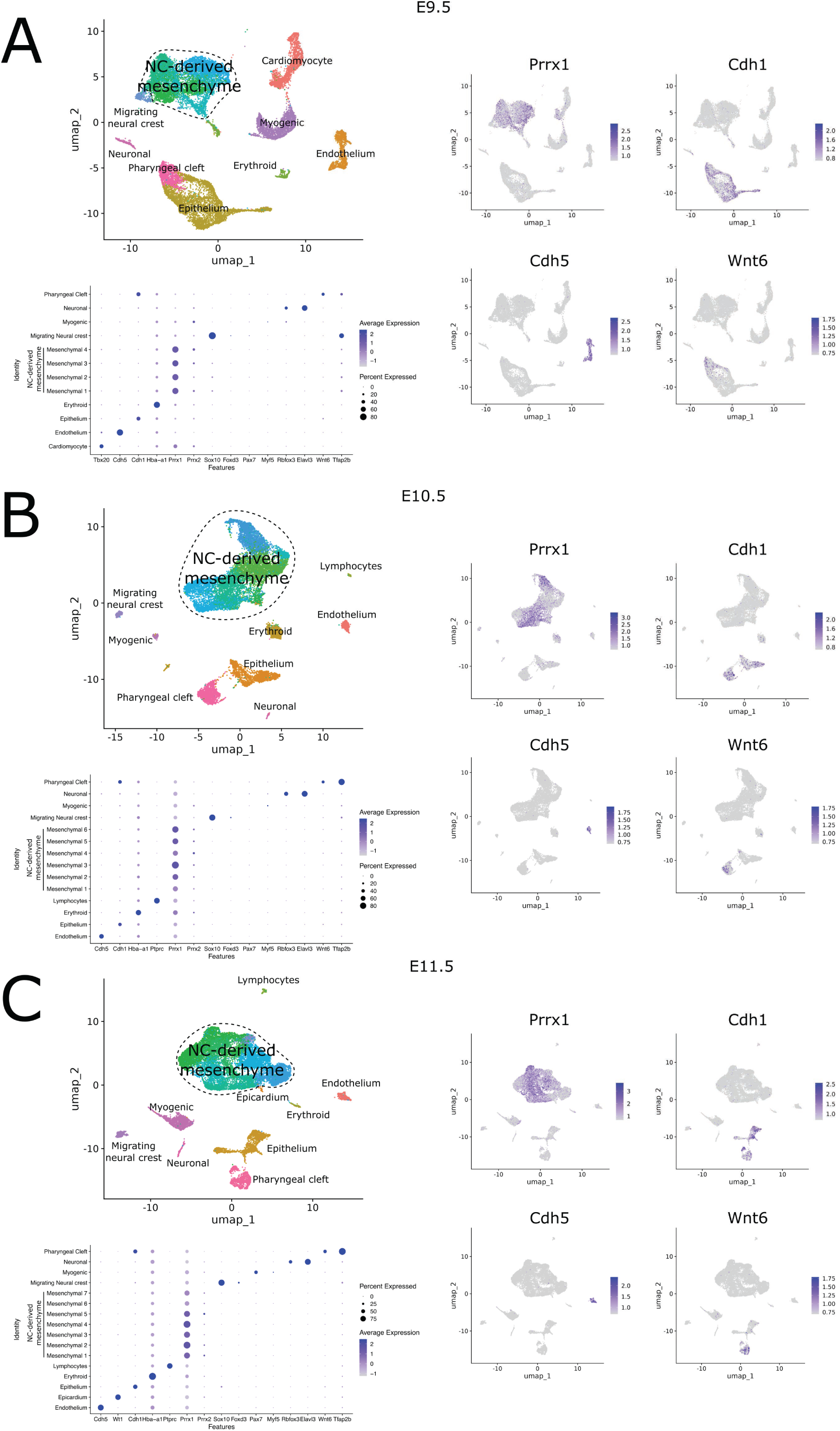
Cell types identified in the pharyngeal arch 2 from snRNA-seq data. Major cell types identified in integrated second pharyngeal arch snRNA-seq datasets from *HACNS1* humanized and chimpanzee ortholog embryos at E9.5 **(A)**, E10.5 **(B)**, and E11.5 **(C)**. The top-left panels for each main figure panel show UMAP embeddings colored by cell type assignments. Neural crest-derived mesenchymal populations are outlined by dashed lines. The bottom-left panels show dot plots illustrating the expression of marker genes across annotated cell types. The dot size represents the percentage of nuclei expressing each gene, and color intensity represents the average scaled expression of the gene within each cell type (see the scales to the right of each panel). The right panels show normalized expression of representative cell type marker genes in UMAP embedding, with increasing color intensity indicating higher expression (see scale bars to the right of each panel).

**Fig. S5.**
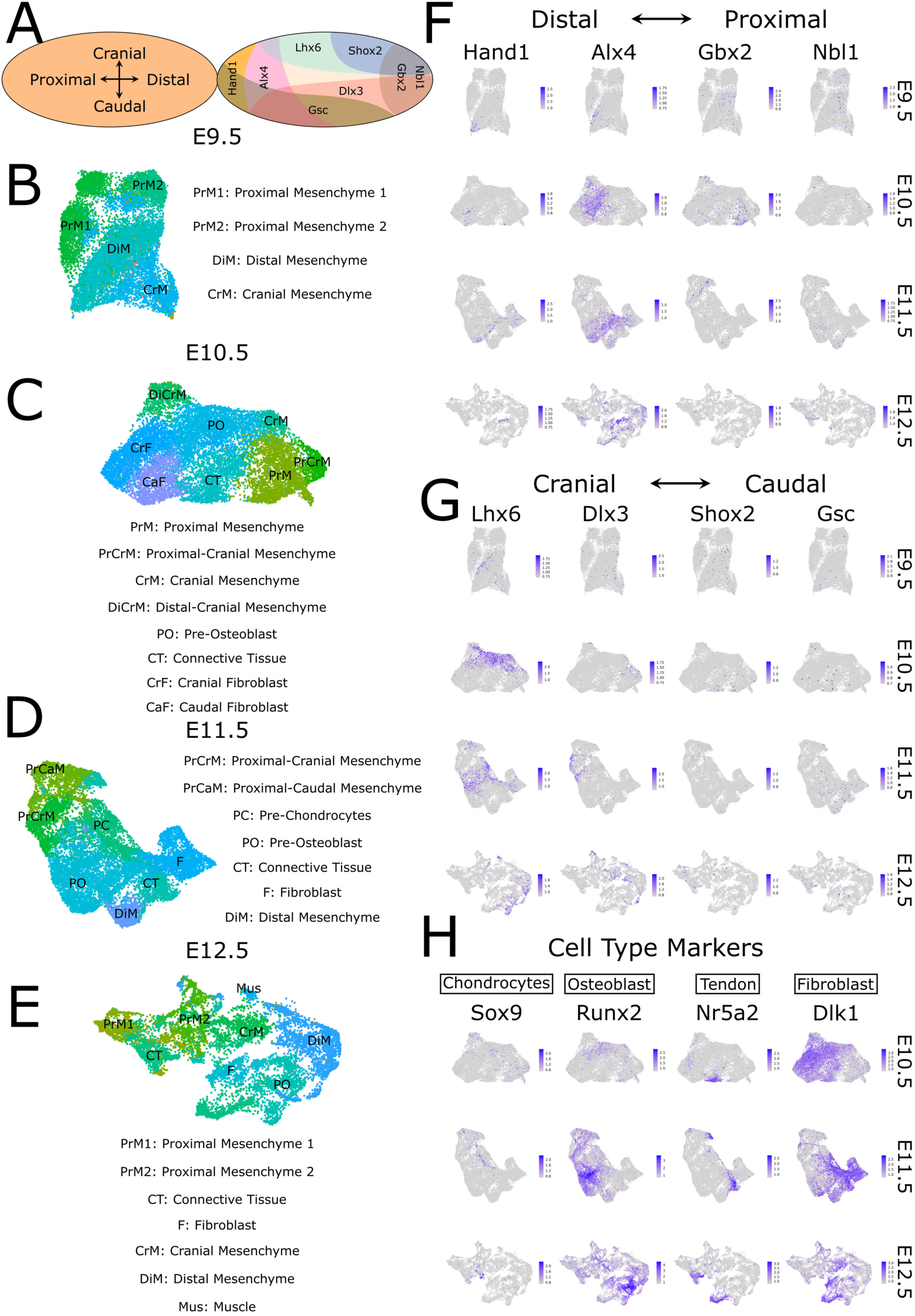
Mesenchymal cell type assignments in the mandibular process. **A:** Schematic of spatial patterning of neural crest-derived mesenchymal cells within the mandibular process. Colored regions indicate overlapping expression domains of marker genes used to define mesenchymal populations along the proximal-distal and cranial-caudal axes. *Hand1* and *Alx4* mark distal domains, whereas *Gbx2* and *Nbl1* mark proximal domains; *Lhx6* and *Shox2* mark cranial domains, and *Dlx3* and *Gsc* mark caudal domains. **B-E:** UMAP embeddings of integrated mandibular process mesenchymal nuclei from *HACNS1* humanized and chimpanzee ortholog embryos colored by cell type assignment at E9.5 **(B)**, E10.5 **(C)**, E11.5 **(D)**, and E12.5 **(E)**. Mesenchymal cell types for each stage are indicated next to their respective UMAP embeddings. **F-G:** UMAP embeddings showing normalized expression of marker genes used to define mesenchymal populations along the distal-proximal **(F)** and cranial-caudal **(G)** axes at E9.5-E12.5. Scales of normalized expression are shown at the right of each plot. **H:** UMAP embeddings showing normalized expression of markers of mesenchymal cell types at E10.5-E12.5, with increasing color intensity indicating higher expression: *Sox9*, chondrocytes; *Runx2*, osteoblasts; *Nr5a2*, tendon; and *Dlk1*, fibroblasts. Scales of normalized expression are shown at the right of each plot.

**Fig. S6.**
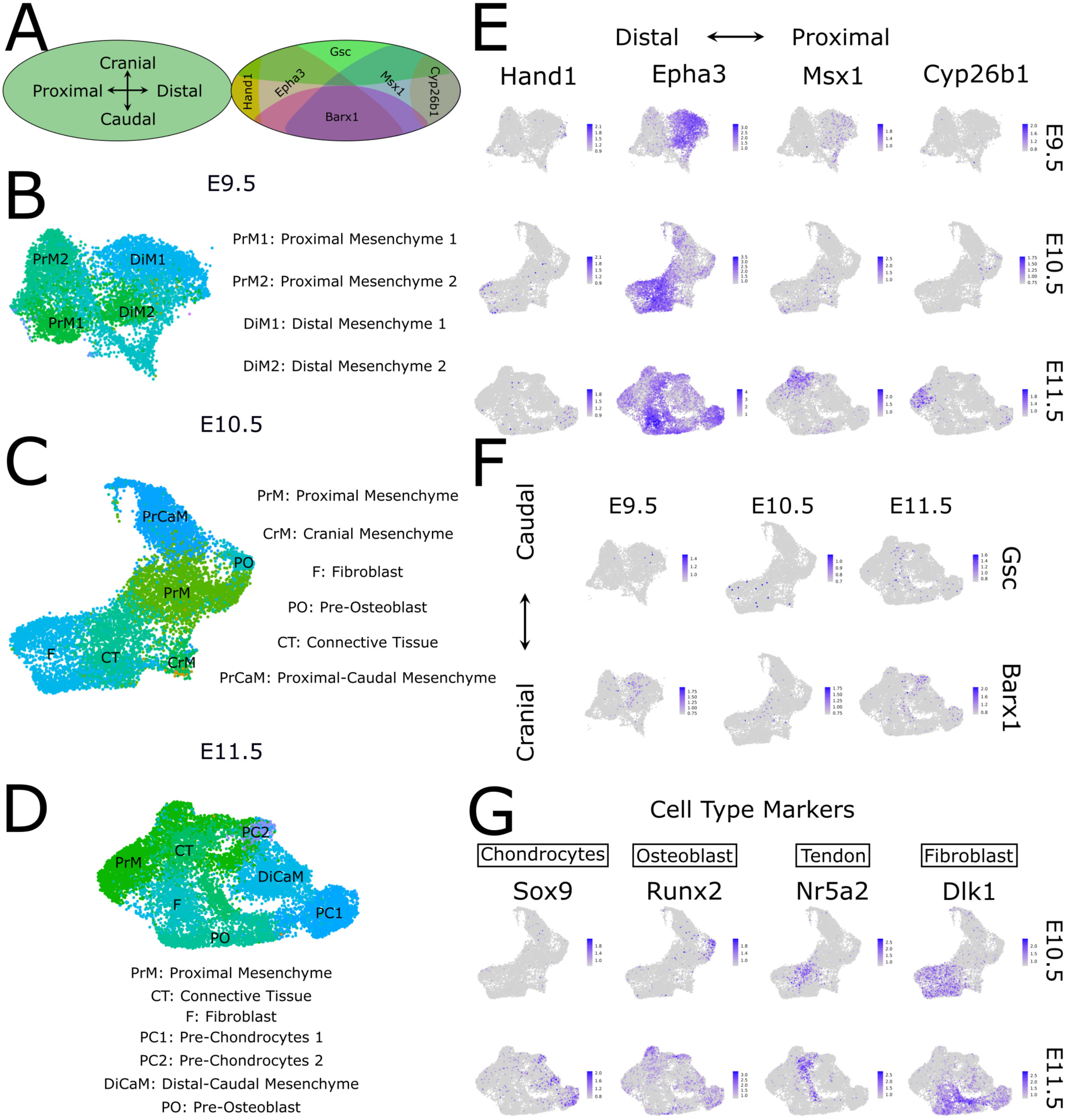
Mesenchymal cell type assignments in the pharyngeal arch 2. **A:** Schematic of spatial patterning of neural crest-derived mesenchymal cells within the second pharyngeal arch. Colored regions indicate overlapping expression domains of marker genes used to define mesenchymal populations along the proximal-distal and cranial-caudal axes. *Hand1* and *Epha3* mark distal domains, whereas *Cyp26b1* and *Msx1* mark proximal domains; *Gsc* marks cranial domains, and *Barx1* marks caudal domains. **B-D:** UMAP embeddings of integrated second pharyngeal arch mesenchymal nuclei from *HACNS1* humanized and chimpanzee ortholog embryos colored by cell type assignment at E9.5 **(B)**, E10.5 **(C)**, and E11.5 **(D)**. Mesenchymal cell types for each stage are indicated next to their respective UMAP embeddings. **E-F:** UMAP embeddings showing normalized expression of marker genes used to define mesenchymal populations along the distal-proximal **(E)** and cranial-caudal **(F)** axes at E9.5-E11.5. Scales of normalized expression are shown at the right of each plot. **G:** UMAP embeddings showing normalized expression of markers of mesenchymal cell types at E10.5-E11.5, with increasing color intensity indicating higher expression: *Sox9*, chondrocytes; *Runx2*, osteoblasts; *Nr5a2*, tendon; and *Dlk1*, fibroblasts. Scales of normalized expression are shown at the right of each plot.

**Fig. S7.**
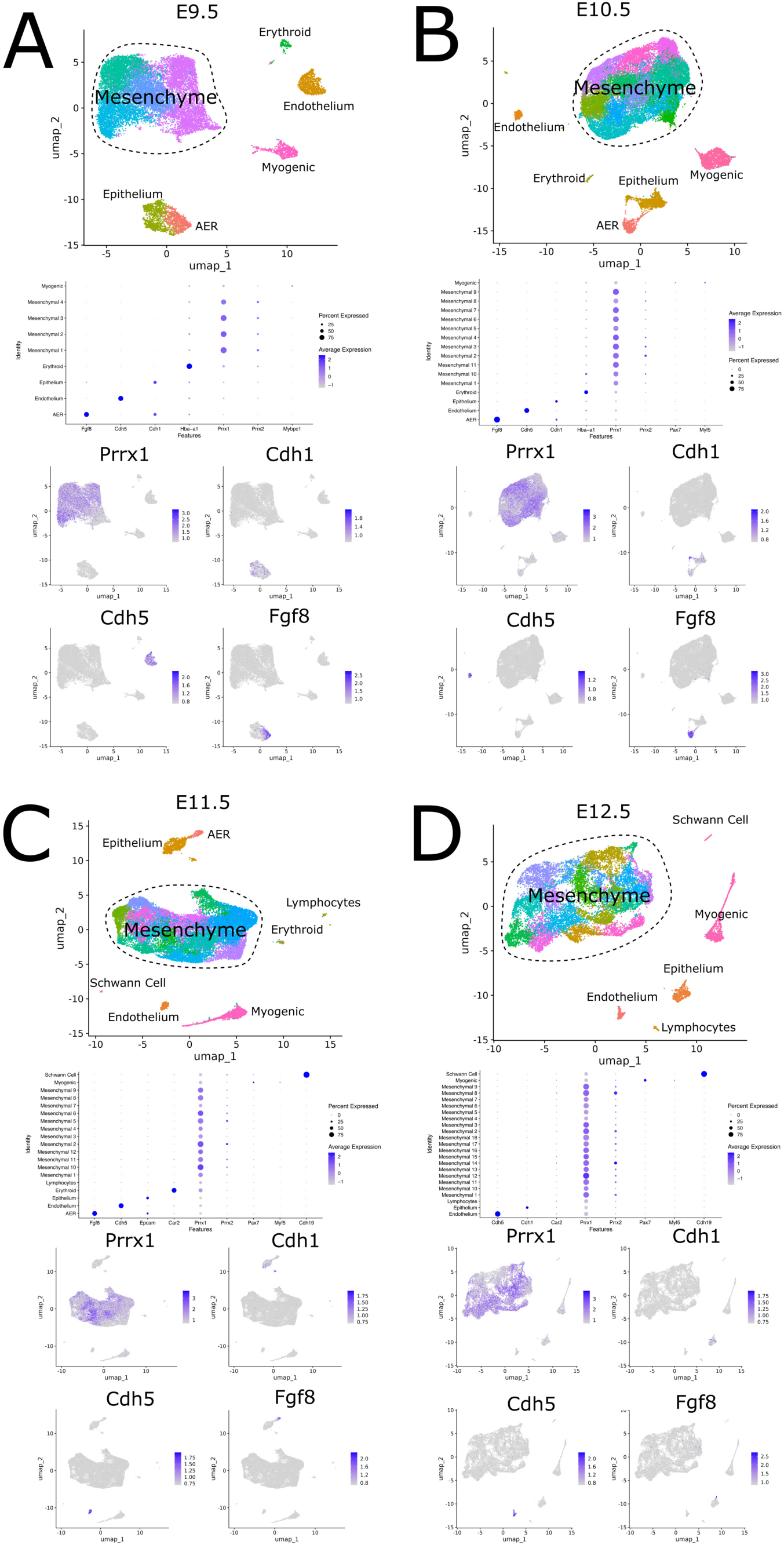
Cell types identified in the forelimb from snRNA-seq data. Major cell types identified in integrated forelimb snRNA-seq datasets from *HACNS1* humanized and chimpanzee ortholog embryos at E9.5 **(A)**, E10.5 **(B)**, E11.5 **(C)**, and E12.5 **(D)**. The top panels for each main figure panel show UMAP embeddings colored by cell type assignments. Mesenchymal populations are outlined by dashed lines. The middle panels show dot plots illustrating the expression of marker genes across annotated cell types. The dot size represents the percentage of nuclei expressing each gene, and color intensity represents the average scaled expression of the gene within each cell type (see the scales to the right of each panel). The bottom panels show normalized expression of representative cell type marker genes in UMAP embedding, with increasing color intensity indicating higher expression (see the scales to the right of each panel).

**Fig. S8.**
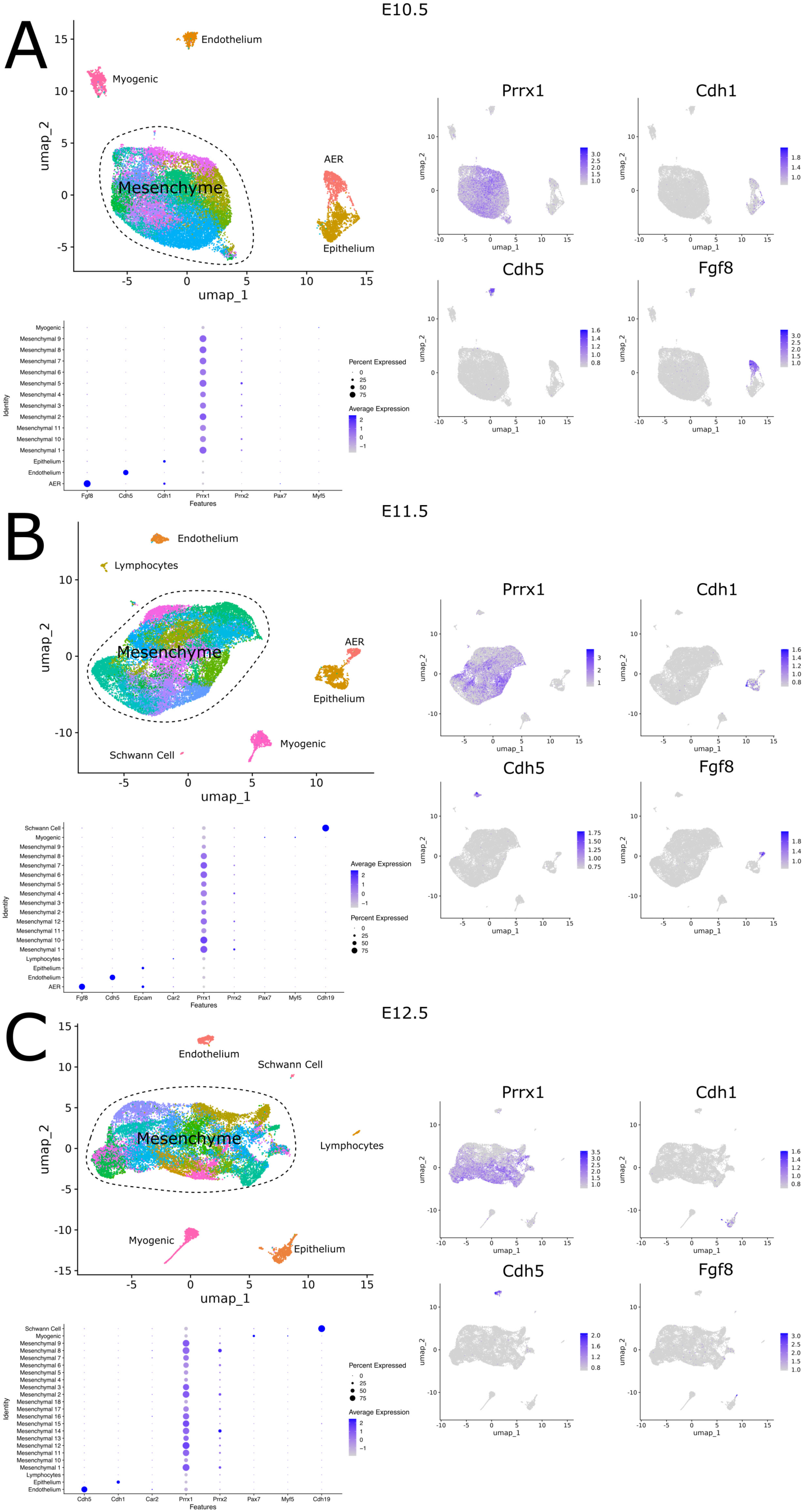
Cell types identified in the hindlimb from snRNA-seq data. Major cell types identified in integrated hindlimb snRNA-seq datasets from *HACNS1* humanized and chimpanzee ortholog embryos at E10.5 **(A)**, E11.5 **(B)**, and E12.5 **(C)**. The top-left panels for each main figure panel show UMAP embeddings colored by cell type assignments. Mesenchymal populations are outlined by dashed lines. The bottom-left panels show dot plots illustrating the expression of selected marker genes across annotated cell types. The dot size represents the percentage of nuclei expressing each gene, and color intensity represents the average scaled expression of the gene within each cell type (see the scales to the right of each panel). The right panels show normalized expression of representative cell type marker genes in UMAP embedding, with increasing color intensity indicating higher expression (see the scales to the right of each panel).

**Fig. S9.**
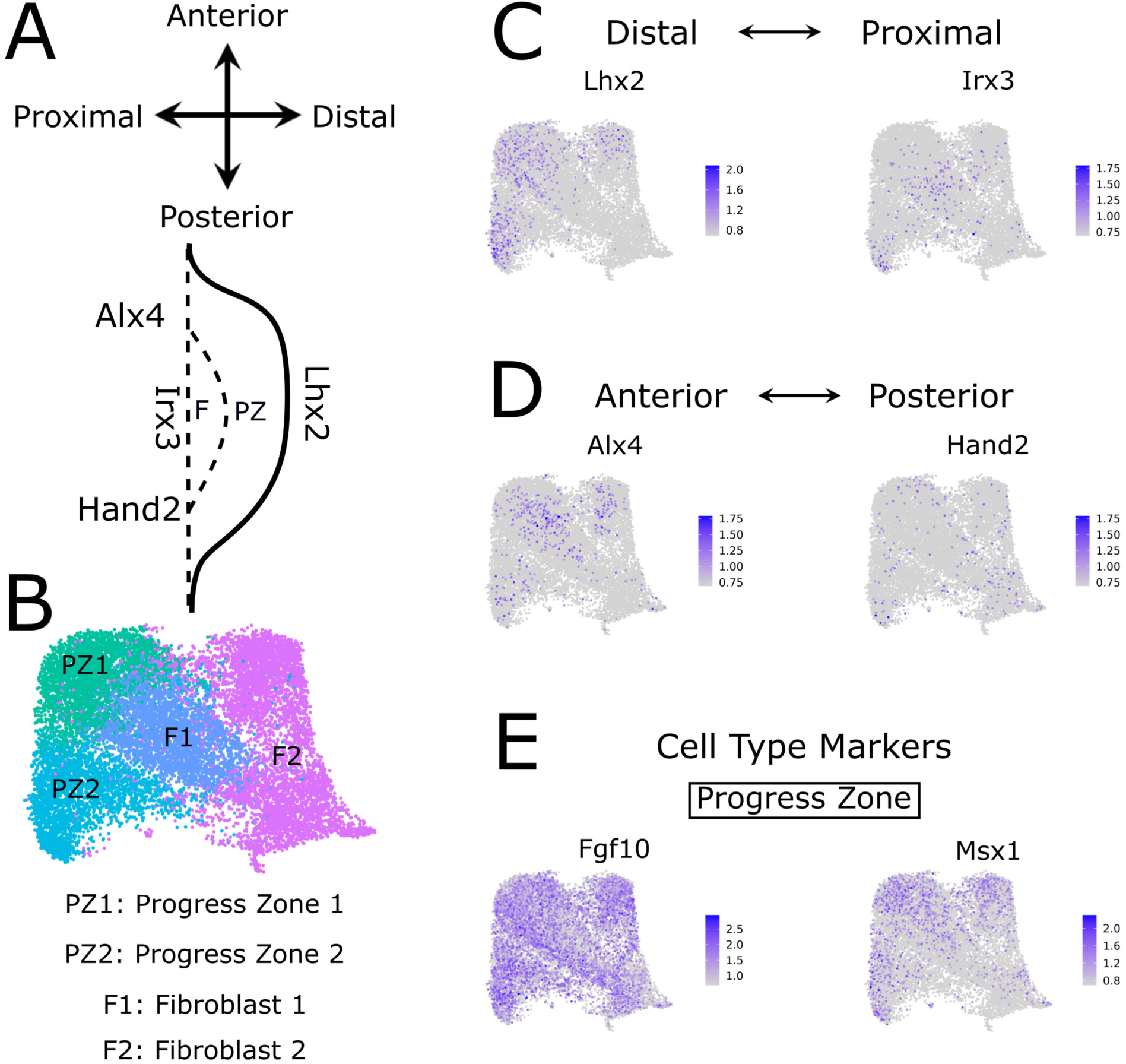
Mesenchymal cell type assignments in E9.5 limb buds. **A:** Schematic of spatial patterning of mesenchymal cells within E9.5 limb buds. *Lhx2* and *Irx3* mark distal and proximal domains, respectively, whereas *Alx4* and *Hand2* mark anterior and posterior domains, respectively. **B:** UMAP embedding of E9.5 limb bud mesenchymal nuclei from *HACNS1* humanized and chimpanzee ortholog embryos colored by cell type assignment. Mesenchymal cell type labels are shown next to the UMAP embedding. **C-D:** UMAP embeddings showing normalized expression of marker genes used to define mesenchymal populations along the distal-proximal **(C)** and anterior-posterior **(D)** axes. Scales of normalized expression are shown at the right of each plot. **E:** UMAP embeddings showing normalized expression of *Fgf10* and *Msx1*, markers of progress zone mesenchymal cell types at E9.5, with increasing color intensity indicating higher expression. Scales of normalized expression are shown at the right of each plot.

**Fig. S10.**
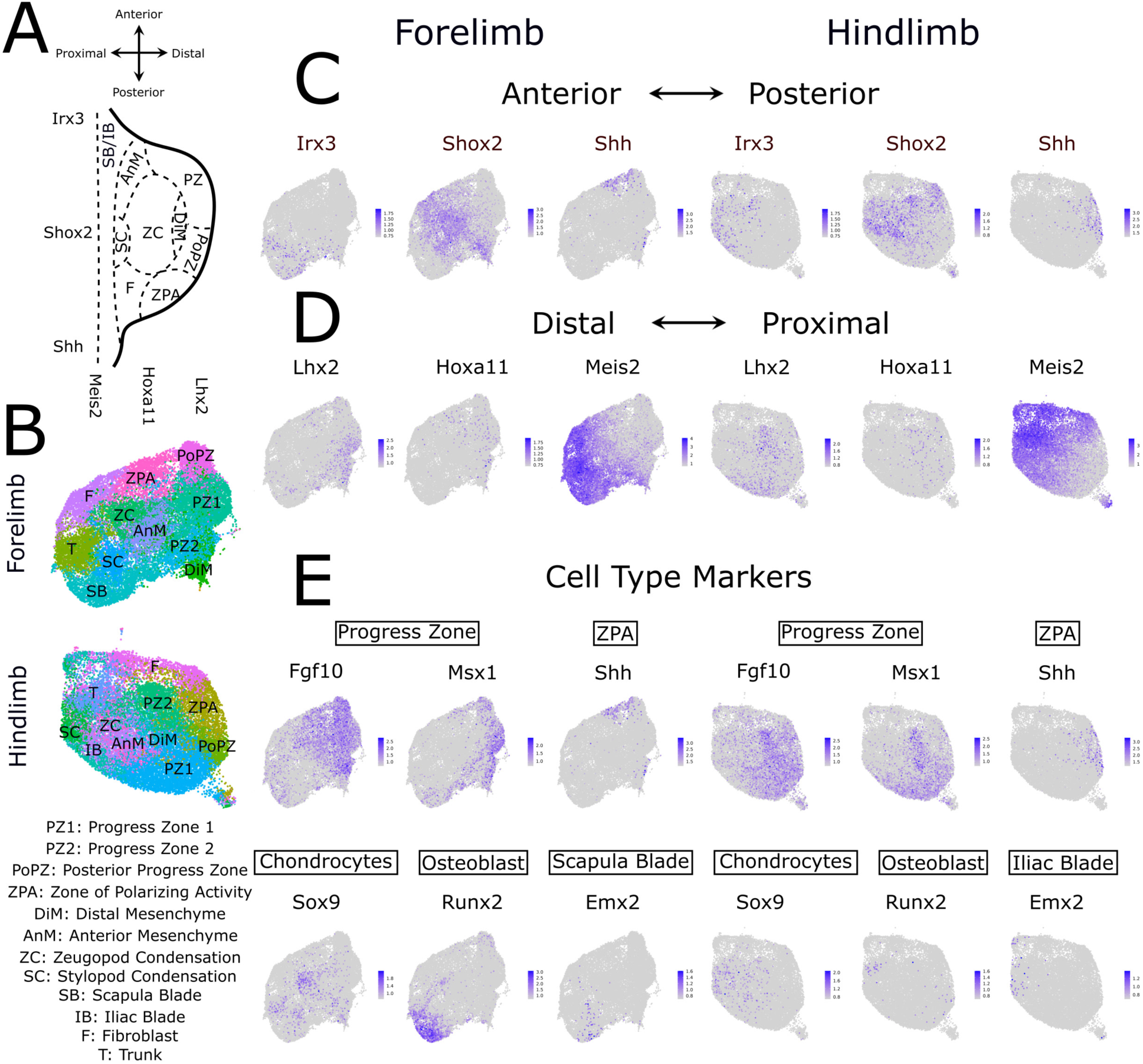
Mesenchymal cell type assignments in E10.5 limb buds. **A:** Schematic of spatial patterning of mesenchymal cells within E10.5 limb buds. *Lhx2*, *Hoxa11*, and *Meis2* mark the distal-proximal axis, whereas *Irx3*, *Shox2*, and *Shh* mark the anterior-posterior axis. **B:** UMAP embeddings of E10.5 forelimb (top) and hindlimb (bottom) mesenchymal nuclei from *HACNS1* humanized and chimpanzee ortholog embryos colored by cell type assignment. Mesenchymal cell type labels are shown next to the UMAP embeddings. **C-D:** UMAP embeddings of E10.5 forelimb (left) and hindlimb (right) showing normalized expression of marker genes used to define mesenchymal populations along the anterior-posterior **(C)** and distal-proximal **(D)** axes. Scales of normalized expression are shown at the right of each plot. **E:** UMAP embeddings showing normalized expression of markers of mesenchymal cell types from forelimb (left) and hindlimb (right), with increasing color intensity indicating higher expression: *Fgf10* and *Msx1*, progress zone; *Shh*, zone of polarizing activity; *Sox9*, chondrocytes; *Runx2*, osteoblasts; and *Emx2*, scapular blade in the forelimb and iliac blade in the hindlimb. Scales of normalized expression are shown at the right of each plot.

**Fig. S11.**
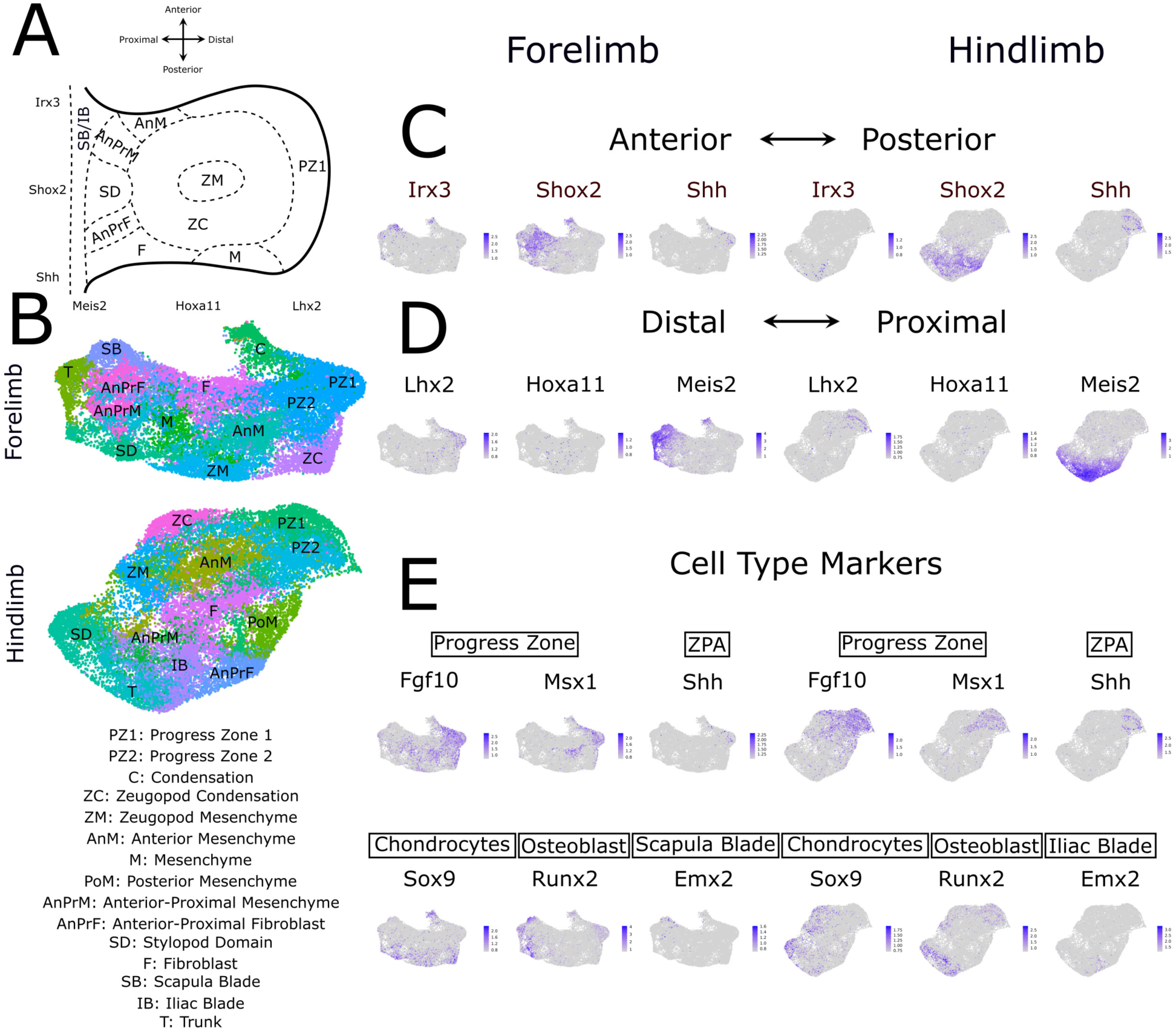
Mesenchymal cell type assignments in E11.5 limb buds. **A:** Schematic of spatial patterning of mesenchymal cells within E11.5 limb buds. *Lhx2*, *Hoxa11*, and *Meis2* mark the distal-proximal axis, whereas *Irx3*, *Shox2*, and *Shh* mark the anterior-posterior axis. **B:** UMAP embeddings of E11.5 forelimb (top) and hindlimb (bottom) mesenchymal nuclei from *HACNS1* humanized and chimpanzee ortholog embryos colored by cell type assignment. Mesenchymal cell type labels are shown next to the UMAP embeddings. **C-D:** UMAP embeddings of E11.5 forelimb (left) and hindlimb (right) showing normalized expression of marker genes used to define mesenchymal populations along the anterior-posterior **(C)** and distal-proximal **(D)** axes. Scales of normalized expression are shown at the right of each plot. **E:** UMAP embeddings showing normalized expression of markers of mesenchymal cell types from forelimb (left) and hindlimb (right), with increasing color intensity indicating higher expression: *Fgf10* and *Msx1*, progress zone; *Shh*, zone of polarizing activity; *Sox9*, chondrocytes; *Runx2*, osteoblasts; and *Emx2*, scapular blade in the forelimb and iliac blade in the hindlimb. Scales of normalized expression are shown at the right of each plot.

**Fig. S12.**
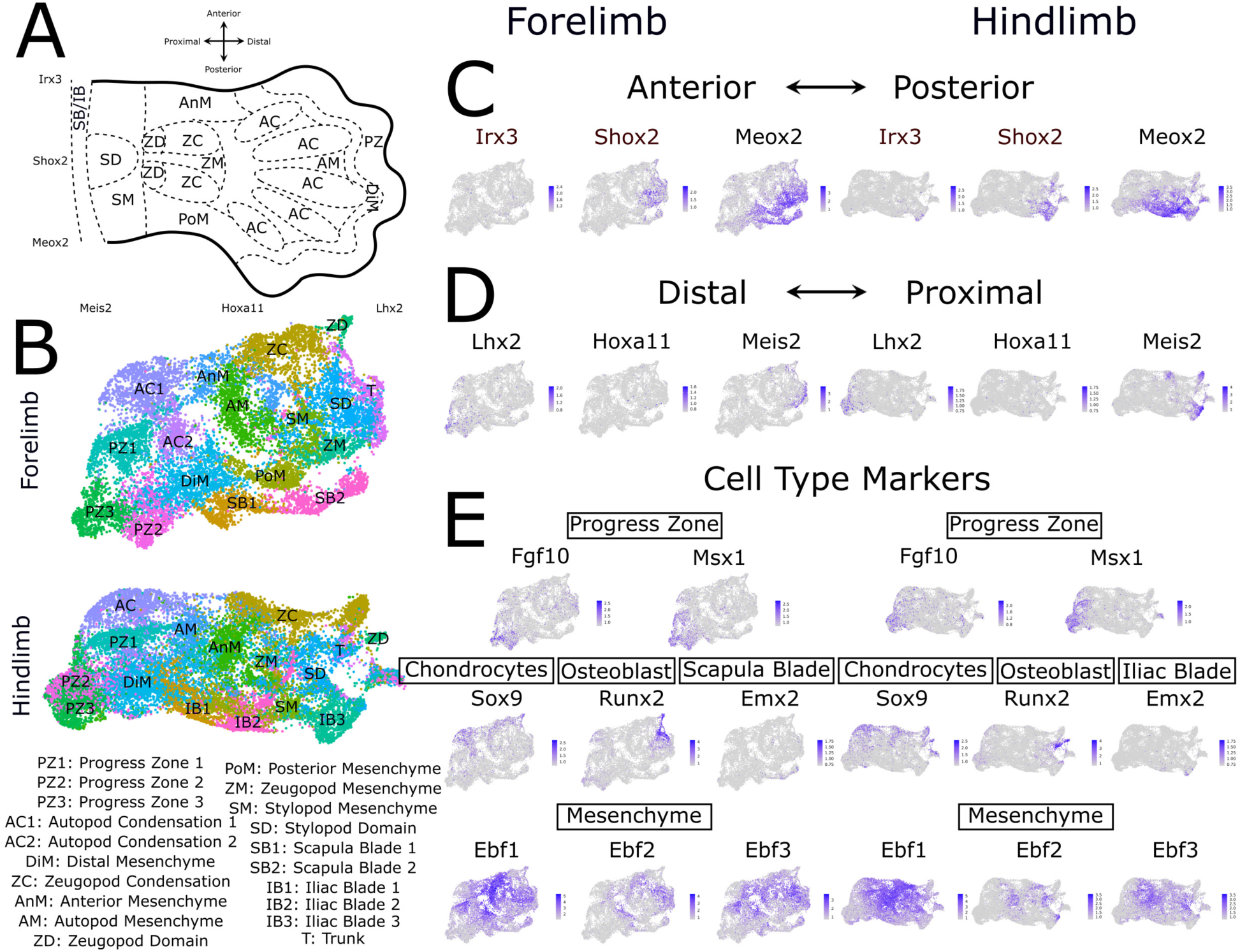
Mesenchymal cell type assignments in E12.5 limb buds. **A:** Schematic of spatial patterning of mesenchymal cells within E12.5 limb buds. *Lhx2*, *Hoxa11*, and *Meis2* mark the distal-proximal axis, whereas *Irx3*, *Shox2*, and *Shh* mark the anterior-posterior axis. **B:** UMAP embeddings of E12.5 forelimb (top) and hindlimb (bottom) mesenchymal nuclei from *HACNS1* humanized and chimpanzee ortholog embryos colored by cell type assignment. Mesenchymal cell type labels are shown next to the UMAP embeddings. **C-D:** UMAP embeddings of E12.5 forelimb (left) and hindlimb (right) showing normalized expression of marker genes used to define mesenchymal populations along the anterior-posterior **(C)** and distal-proximal **(D)** axes. Scales of normalized expression are shown at the right of each plot. **E:** UMAP embeddings showing normalized expression of markers of mesenchymal cell types from forelimb (left) and hindlimb (right), with increasing color intensity indicating higher expression: *Fgf10* and *Msx1*, progress zone; *Shh*, zone of polarizing activity; *Sox9*, chondrocytes; *Runx2*, osteoblasts; and *Emx2*, scapular blade in the forelimb and iliac blade in the hindlimb. Scales of normalized expression are shown at the right of each plot.

**Fig. S13.**
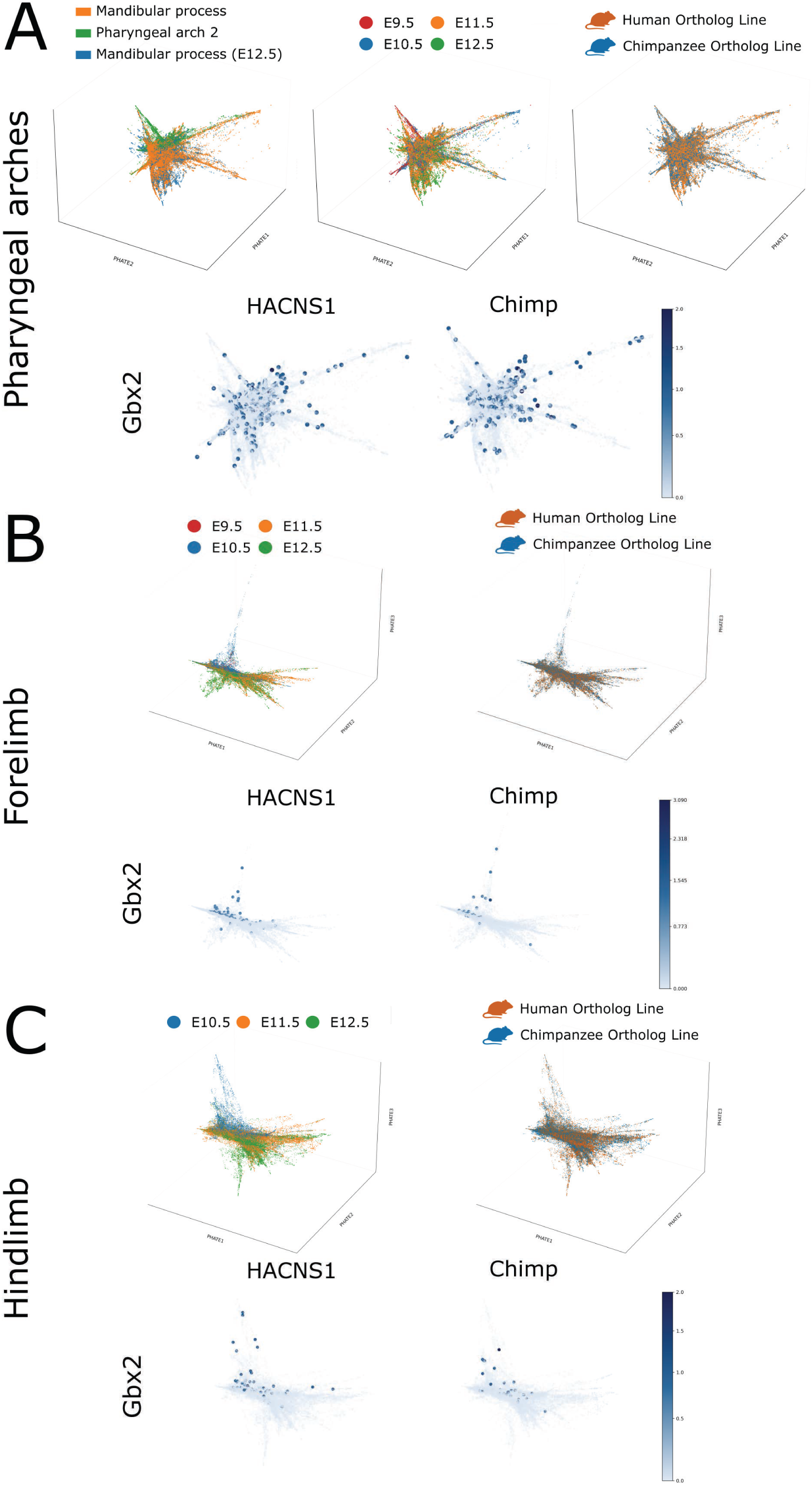
3D PHATE embedding of integrated mesenchymal nuclei. **A-C:** 3D PHATE embeddings of integrated mesenchymal nuclei from the pharyngeal arches **(A)**, forelimb **(B)**, and hindlimb **(C)**. Nuclei from both the *HACNS1* humanized line and the chimpanzee ortholog line, and all developmental stages, are included. In **A**, the top panels show nuclei colored by tissue, developmental stage, and genotype. The bottom panels show 3D PHATE embeddings illustrating normalized *Gbx2* expression for the *HACNS1* humanized (left) and chimpanzee ortholog (right) lines. Scales of normalized expression are shown at the right of each plot. In **B-C**, the top panels show nuclei colored by developmental stage and genotype. The bottom panels show 3D PHATE embeddings illustrating normalized *Gbx2* expression for the *HACNS1* humanized (left) and chimpanzee ortholog (right) lines. Scales of normalized expression are shown at the right of each plot. See also Figure 2.

**Fig. S14.**
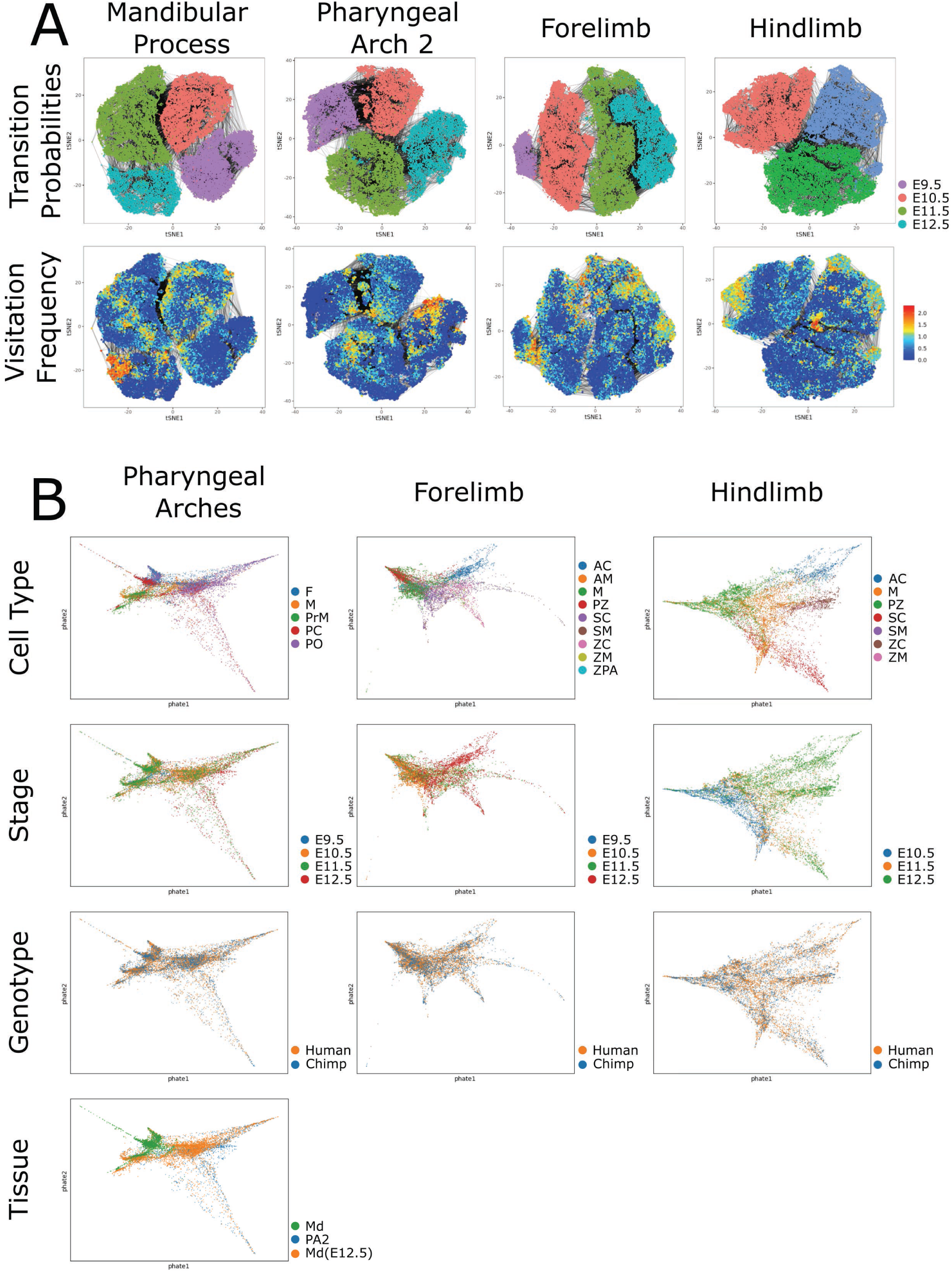
Workflow for reconstructing mesenchymal differentiation trajectories from snRNA-seq data. **A:** Biased random walking analysis of integrated mesenchymal nuclei from both the *HACNS1* humanized line and the chimpanzee ortholog line, and all developmental stages of the mandibular process, second pharyngeal arch, forelimb, and hindlimb using URD (see Methods). The top panels show tSNE embeddings of integrated mesenchymal nuclei colored by developmental stage, with gray connecting lines representing transition probabilities between neighboring nuclei. The bottom panels show visitation frequencies derived from biased random walks, with low frequencies shown in blue and high frequencies in red. The color scale of visitation frequencies is shown on the right. **B:** PHATE embeddings of nuclei with high visitation frequencies associated with chondroblast or osteoblast differentiation trajectories in the pharyngeal arches (left), forelimb (middle), and hindlimb (right). Nuclei are colored by cell type assignment, developmental stage, genotype, and, for the pharyngeal arches, tissue of origin.

**Fig. S15.**
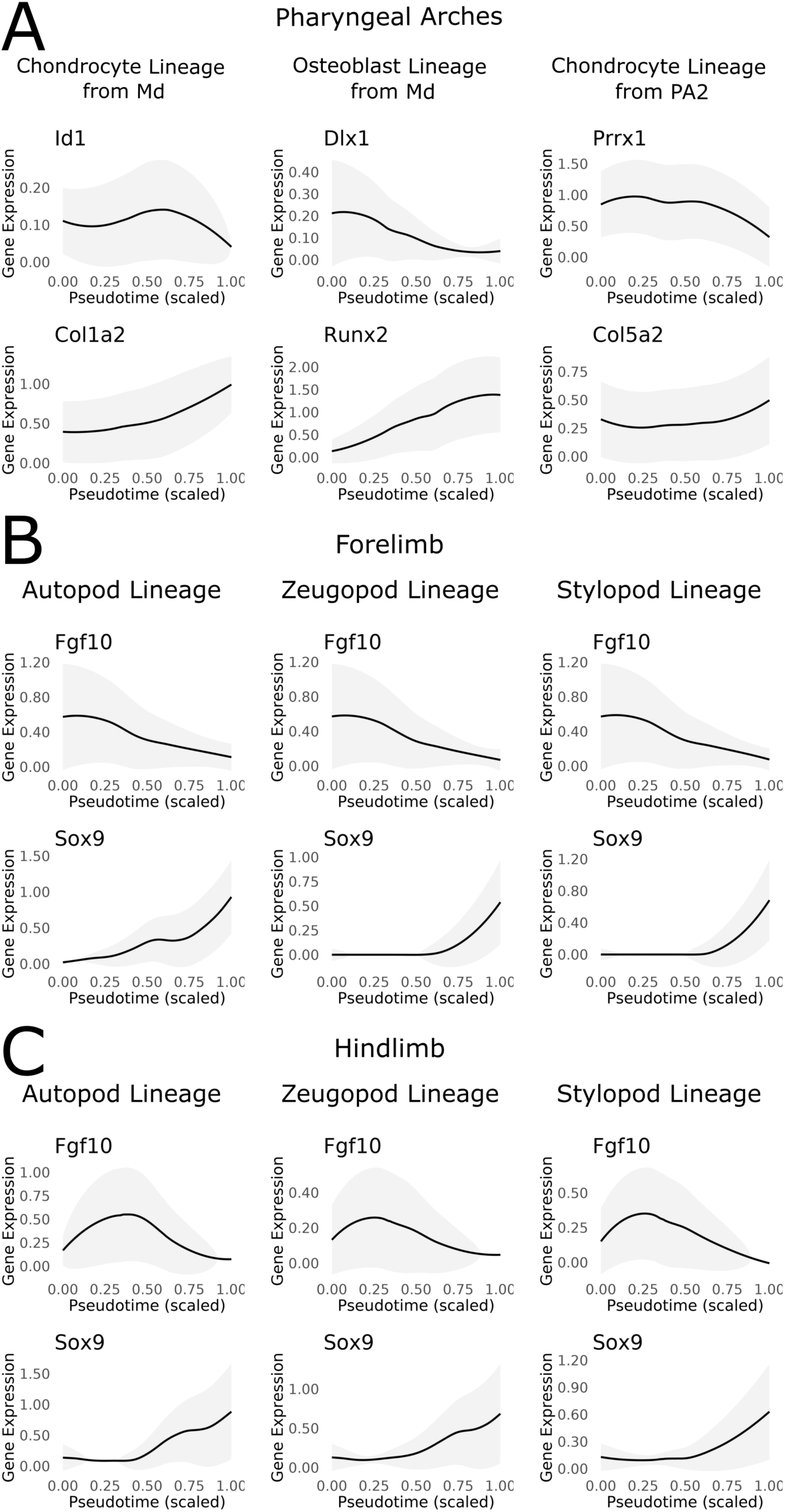
Marker gene expression along mesenchymal differentiation trajectories. **A-C:** Expression of representative marker genes along mesenchymal differentiation trajectories in the pharyngeal arches **(A)**, forelimb **(B)**, and hindlimb **(C)**. Nuclei from both the *HACNS1* humanized and chimpanzee ortholog lines are included in each trajectory. **A:** Chondrocyte and osteoblast lineages originating from the mandibular process (Md) and the chondrocyte lineage originating from the second pharyngeal arch (PA2). **B-C:** Autopod, zeugopod, and stylopod lineages in the forelimb **(B)** and hindlimb **(C)**. For each plot, pseudotime is scaled on the horizontal axis from 0 to 1 for each trajectory. The vertical axis shows normalized gene expression among nuclei assigned to the indicated trajectory. The black line represents average normalized gene expression along pseudotime, and the gray shading indicates standard deviation. See also Figure 3.

**Fig. S16.**
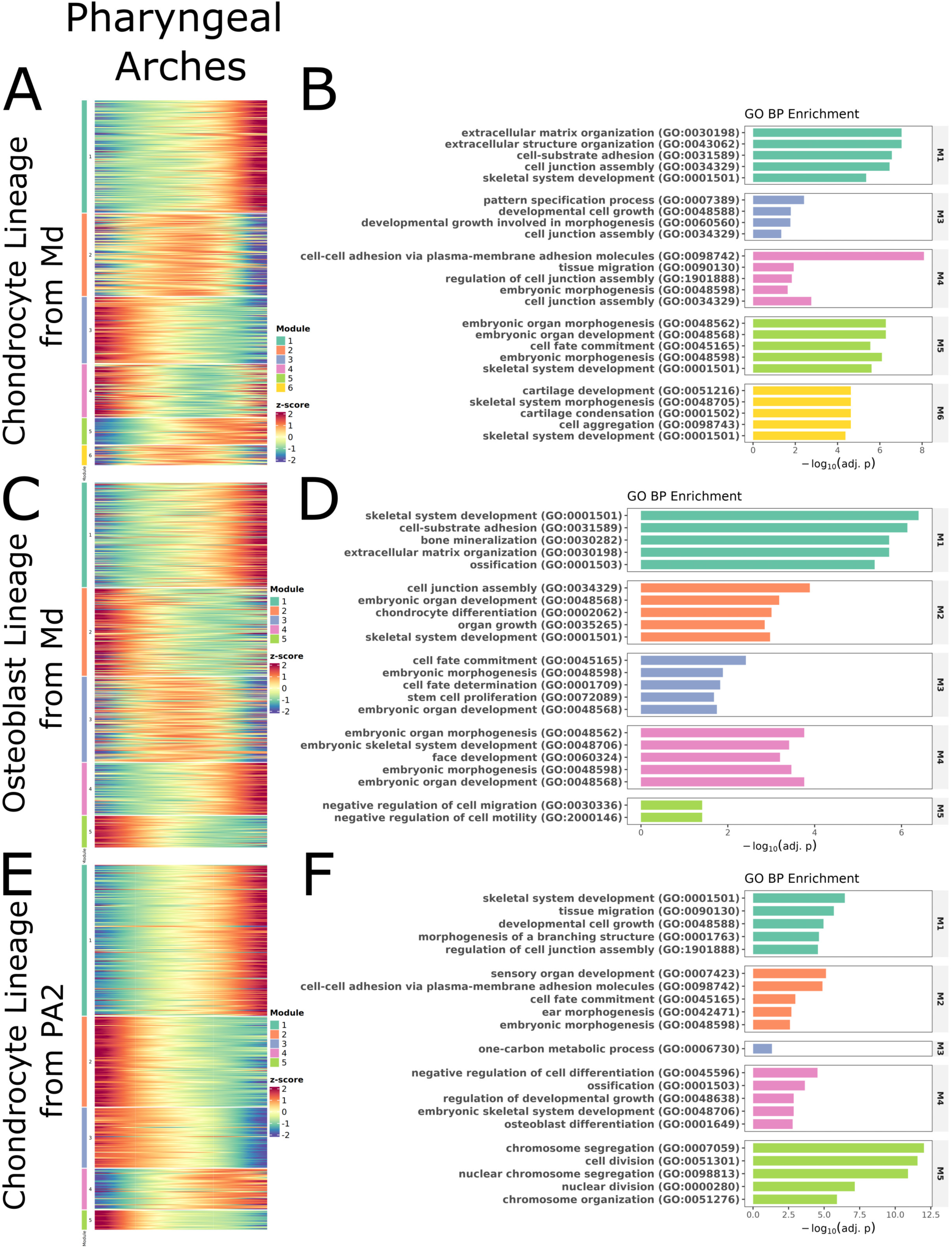
Gene expression modules associated with mesenchymal differentiation trajectories in the pharyngeal arches. **A, C, E:** Heatmaps of genes whose expression is significantly associated with pseudotime along the developmental trajectory for the chondrocyte lineage from the mandibular process **(A)**, the osteoblast lineage from the mandibular process **(C)**, and the chondrocyte lineage from PA2 **(E)**. Nuclei from both the *HACNS1* humanized and chimpanzee ortholog lines are ordered by pseudotime from left to right in each heatmap. Gene expression values are scaled using z-scores, with red indicating high expression and blue indicating low expression. Scales of z-scores are shown at the right of each heatmap. Genes are clustered into modules based on shared expression dynamics along each trajectory. The colored bar to the left of each heatmap indicates the module assigned to each gene, with the module numbers and corresponding colors defined in the Module legend at the right of each heatmap. **B, D, F:** GO enrichment analysis of shifted genes for each module in the chondrocyte lineage from the mandibular process **(B)**, the osteoblast lineage from the mandibular process **(D)**, and the chondrocyte lineage from PA2 **(F)**. The bar plots show significantly enriched GO terms. The horizontal axis shows enrichment significance as −log10(adjusted P value), and the bars are colored by module assignment.

**Fig. S17.**
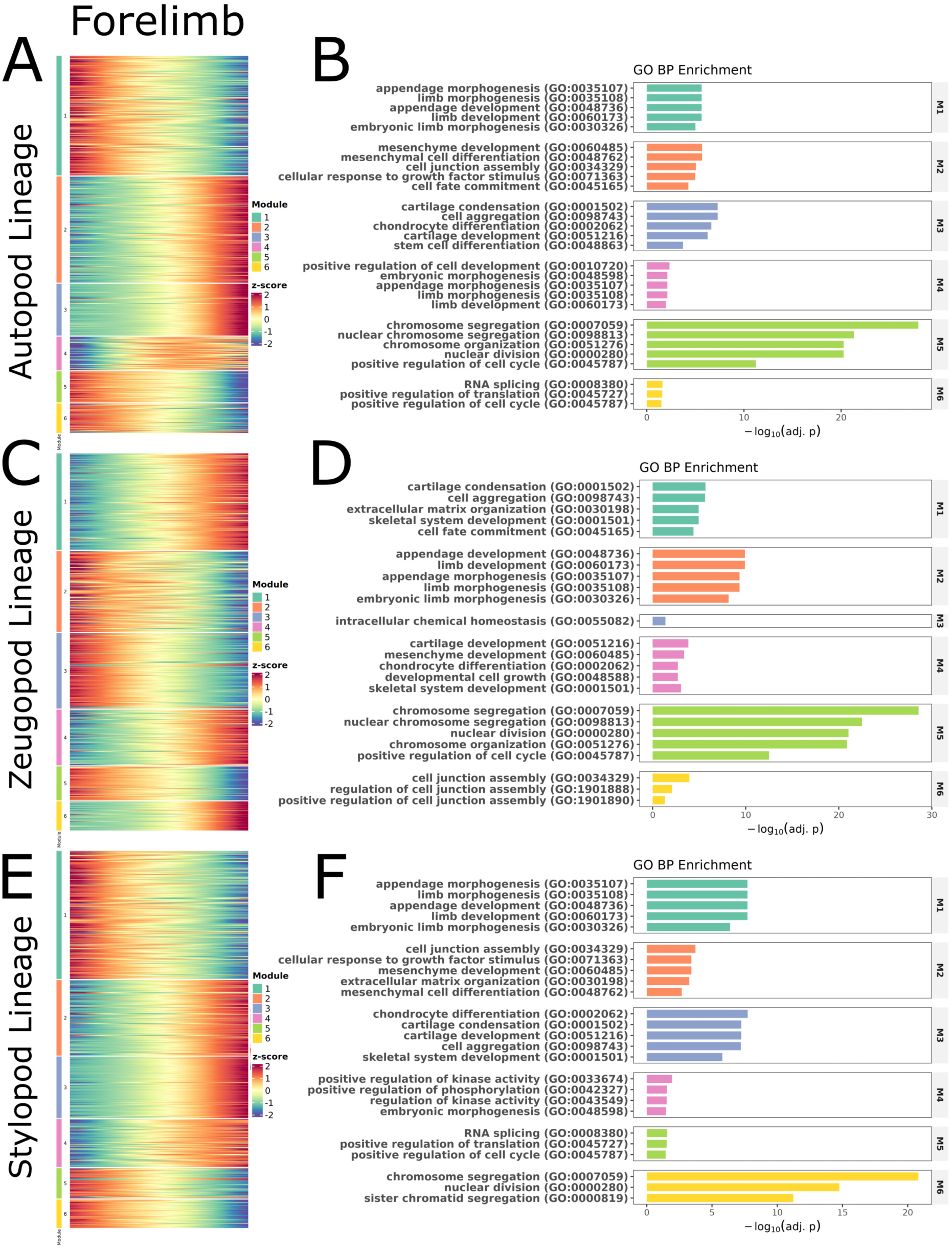
Gene expression modules associated with mesenchymal differentiation trajectories in the forelimb. **A, C, E:** Heatmaps of genes whose expression is significantly associated with pseudotime along the developmental trajectory for the autopod lineage **(A)**, the zeugopod lineage **(C)**, and the stylopod lineage **(E)** in the forelimb. Nuclei from both the *HACNS1* humanized and chimpanzee ortholog lines are ordered by pseudotime from left to right in each heatmap. Gene expression values are scaled using z-scores, with red indicating high expression and blue indicating low expression. Scales of z-scores are shown at the right of each heatmap. Genes are clustered into modules based on shared expression dynamics along each trajectory. The colored bar to the left of each heatmap indicates the module assigned to each gene, with the module numbers and corresponding colors defined in the Module legend at the right of each heatmap. **B, D, F:** GO enrichment analysis of shifted genes for each module in the chondrocyte lineage from the autopod lineage **(B)**, the zeugopod lineage **(D)**, and the stylopod lineage **(F)** in the forelimb. The bar plots show significantly enriched GO terms. The horizontal axis shows enrichment significance as −log10(adjusted P value), and the bars are colored by module assignment.

**Fig. S18.**
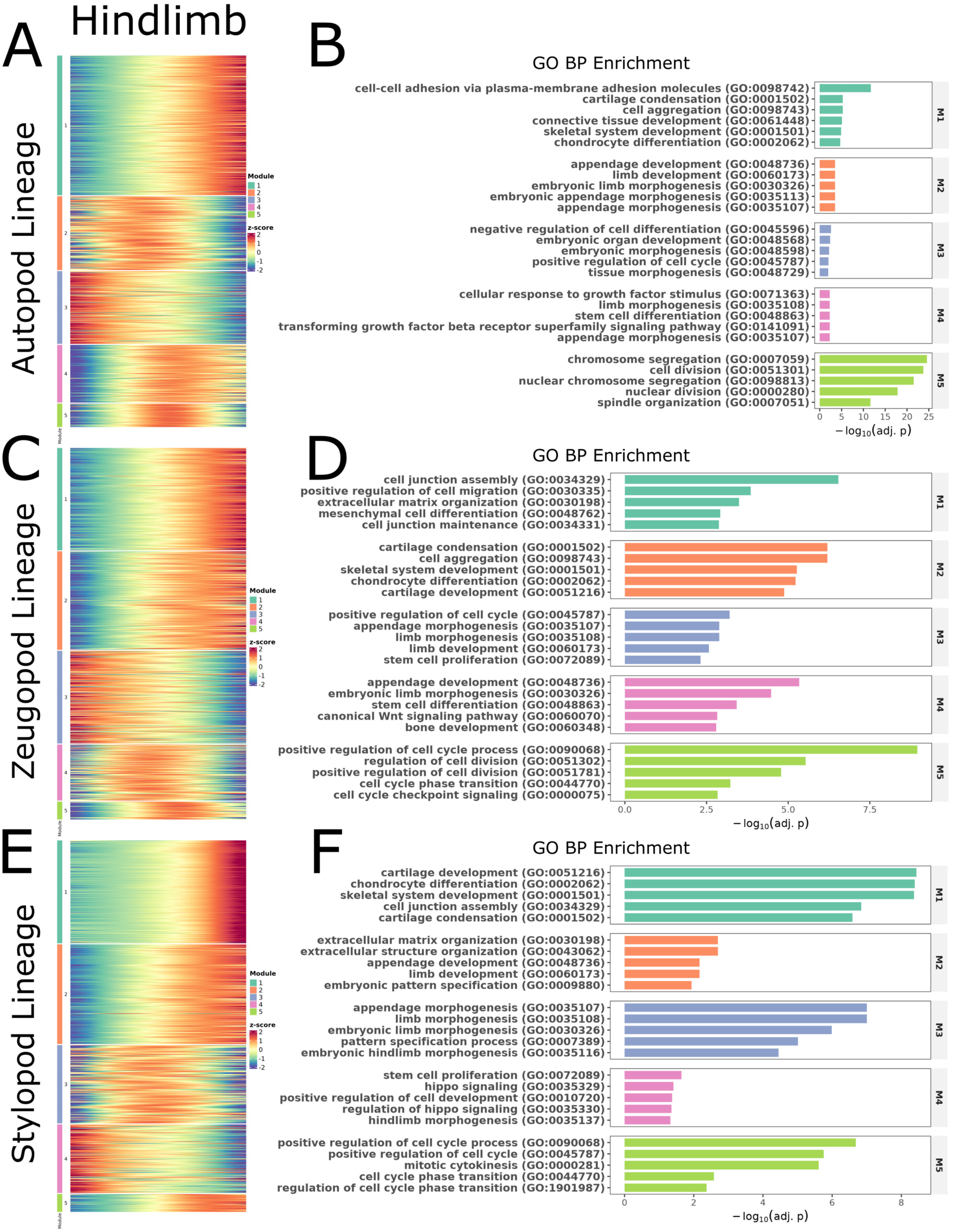
Gene expression modules associated with mesenchymal differentiation trajectories in the hindlimb. **A, C, E:** Heatmaps of genes whose expression is significantly associated with pseudotime along the developmental trajectory for the autopod lineage **(A)**, the zeugopod lineage **(C)**, and the stylopod lineage **(E)** in the hindlimb. Nuclei from both the *HACNS1* humanized and chimpanzee ortholog lines are ordered by pseudotime from left to right in each heatmap. Gene expression values are scaled using z-scores, with red indicating high expression and blue indicating low expression. Scales of z-scores are shown at the right of each heatmap. Genes are clustered into modules based on shared expression dynamics along each trajectory. The colored bar to the left of each heatmap indicates the module assigned to each gene, with the module numbers and corresponding colors defined in the Module legend at the right of each heatmap. **B, D, F:** GO enrichment analysis of shifted genes for each module in the chondrocyte lineage from the autopod lineage **(B)**, the zeugopod lineage **(D)**, and the stylopod lineage **(F)** in the hindlimb. The bar plots show significantly enriched GO terms. The horizontal axis shows enrichment significance as −log10(adjusted P value), and the bars are colored by module assignment.

**Fig. S19.**
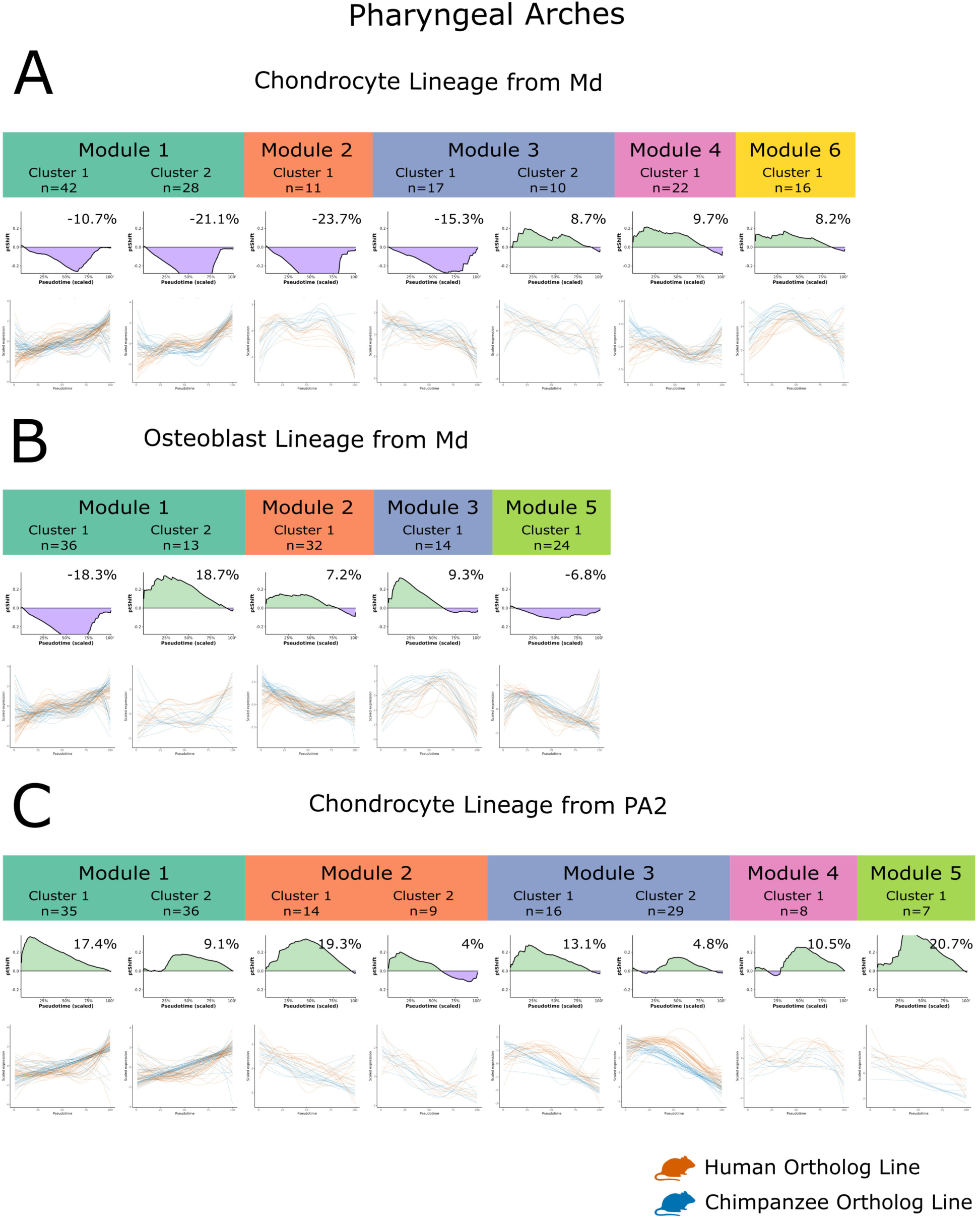
Pseudotime shifts of genes along mesenchymal differentiation trajectories in the pharyngeal arches. **A-C:** Pseudotime shifts of genes from each module along mesenchymal differentiation trajectories for the chondrocyte lineage from the mandibular process **(A)**, the osteoblast lineage from the mandibular process **(B)**, and the chondrocyte lineage from PA2 **(C)**. Within each gene module, genes were further grouped into clusters based on their pseudotime shift (ptShift) similarities. The colored module labels above the plots indicate the module assignment of the clusters shown below, and the colors correspond to those used in Fig. S16. The cluster label and number of genes in that cluster (n) are shown in the header. The top panels in each main figure panel show ptShift profiles along the developmental trajectory for each gene cluster within the indicated module. The horizontal axis shows scaled pseudotime, and the vertical axis shows ptShift. Positive values indicate relative developmental delay (green), whereas negative values indicate relative developmental acceleration (purple) in the *HACNS1* humanized line compared with the chimpanzee control. The percentage above each plot indicates the mean pseudotime shift for that cluster. The bottom panels in each main figure panel show normalized gene expression for all genes within each cluster plotted along pseudotime. Each line represents one gene; orange lines indicate the *HACNS1* humanized line and blue lines indicate the chimpanzee ortholog line. The horizontal axis shows pseudotime, and the vertical axis shows scaled gene expression.

**Fig. S20.**
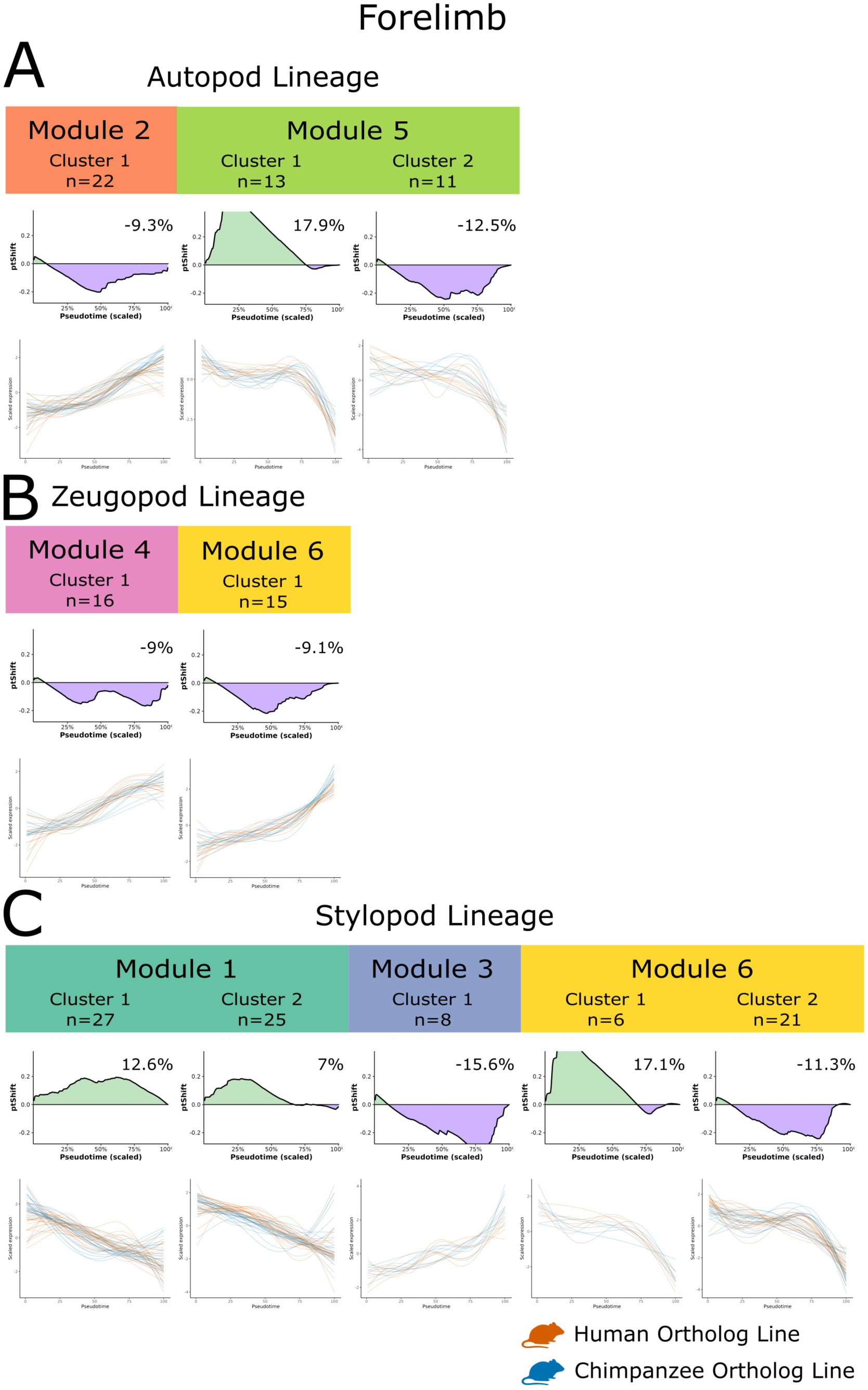
Pseudotime shifts of genes along mesenchymal differentiation trajectories in the forelimb. **A-C :** Pseudotime shifts of genes from each module along mesenchymal differentiation trajectories for the autopod lineage **(A)**, the zeugopod lineage **(B)**, and the stylopod lineage **(C)**. Within each gene module, genes were further grouped into clusters based on their pseudotime shift (ptShift) similarities. The colored module labels above the plots indicate the module assignment of the clusters shown below, and the colors correspond to those used in Fig. S17. The cluster label and number of genes in that cluster (n) are shown in the header. The top panels in each main figure panel show ptShift profiles along the developmental trajectory for each gene cluster within the indicated module. The horizontal axis shows scaled pseudotime, and the vertical axis shows ptShift. Positive values indicate relative developmental delay (green), whereas negative values indicate relative developmental acceleration (purple) in the *HACNS1* humanized line compared with the chimpanzee control. The percentage above each plot indicates the mean pseudotime shift for that cluster. The bottom panels in each main figure panel show normalized gene expression for all genes within each cluster plotted along pseudotime. Each line represents one gene; orange lines indicate the *HACNS1* humanized line and blue lines indicate the chimpanzee ortholog line. The horizontal axis shows pseudotime, and the vertical axis shows scaled gene expression.

**Fig. S21.**
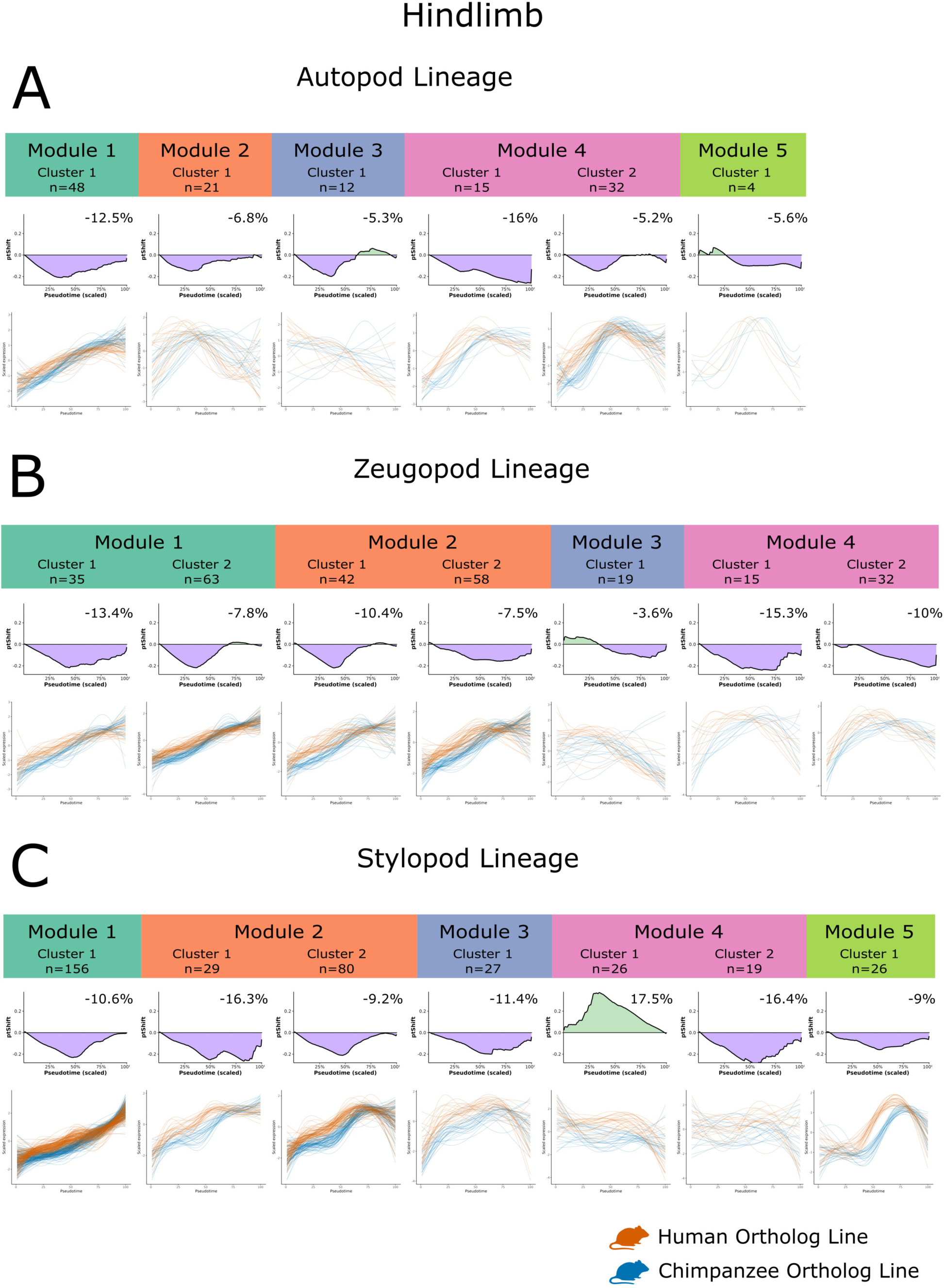
Pseudotime shifts of genes along mesenchymal differentiation trajectories in the hindlimb. **A-C :** Pseudotime shifts of genes from each module along mesenchymal differentiation trajectories for the autopod lineage **(A)**, the zeugopod lineage **(B)**, and the stylopod lineage **(C)**. Within each gene module, genes were further grouped into clusters based on their pseudotime shift (ptShift) similarities. The colored module labels above the plots indicate the module assignment of the clusters shown below, and the colors correspond to those used in Fig. S18. The cluster label and number of genes in that cluster (n) are shown in the header. The top panels in each main figure panel show ptShift profiles along the developmental trajectory for each gene cluster within the indicated module. The horizontal axis shows scaled pseudotime, and the vertical axis shows ptShift. Positive values indicate relative developmental delay (green), whereas negative values indicate relative developmental acceleration (purple) in the *HACNS1* humanized line compared with the chimpanzee control. The percentage above each plot indicates the mean pseudotime shift for that cluster. The bottom panels in each main figure panel show normalized gene expression for all genes within each cluster plotted along pseudotime. Each line represents one gene; orange lines indicate the *HACNS1* humanized line and blue lines indicate the chimpanzee ortholog line. The horizontal axis shows pseudotime, and the vertical axis shows scaled gene expression.

**Fig. S22.**
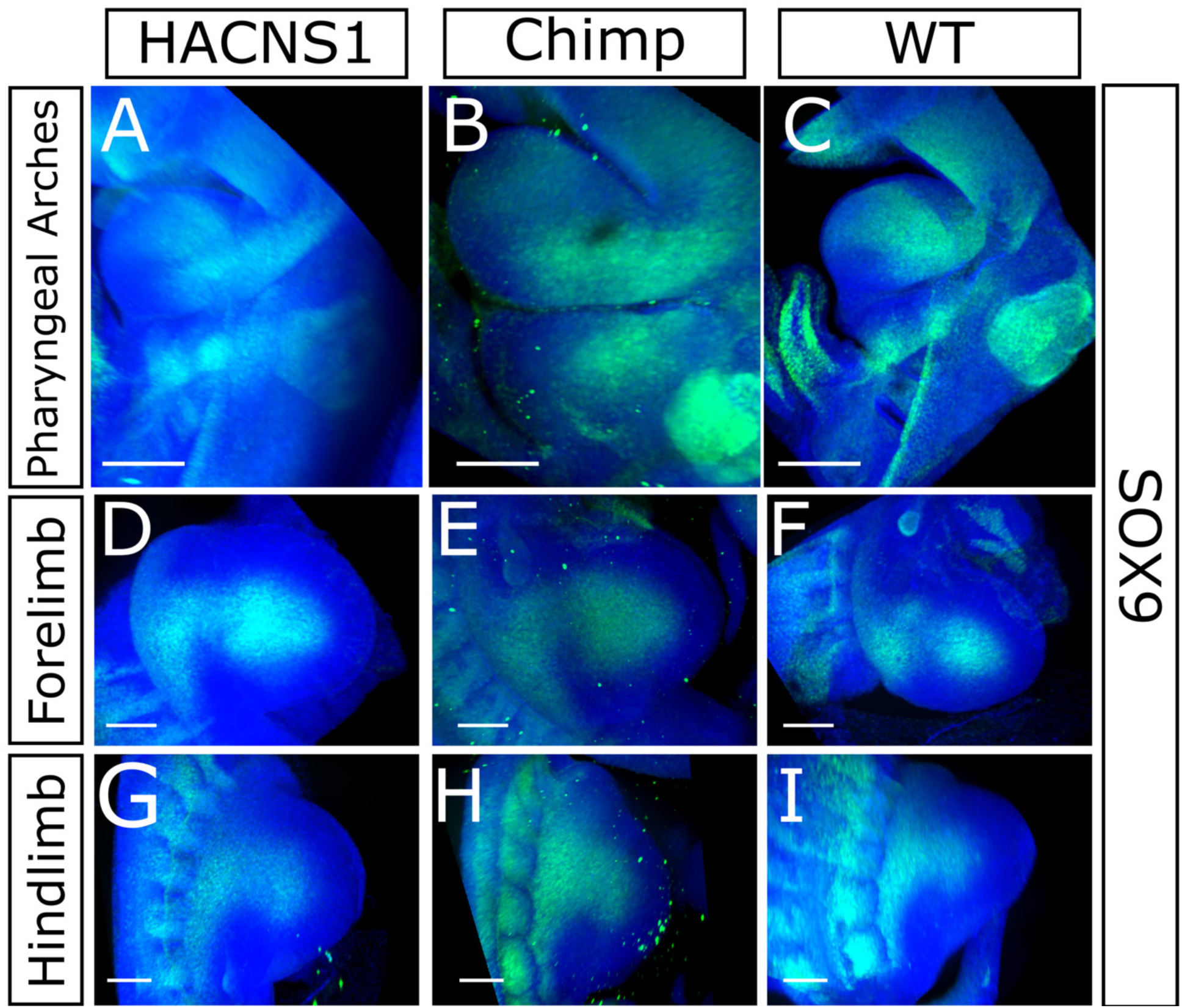
SOX9 expression in E10.5 embryos. Representative immunofluorescence staining for SOX9 in E10.5 *HACNS1* humanized, chimpanzee ortholog, and wild-type control embryos (left, center and right). SOX9 expression is shown in the pharyngeal arches **(A-C)**, forelimb **(D-F)**, and hindlimb **(G-I)** for each indicated genotype. Images were collected from clarified embryos. SOX9 is shown in green, and DAPI nuclear staining is shown in blue. Scale bar, 200 μm; See also Supplemental Movies S4-S12.

**Fig. S23.**
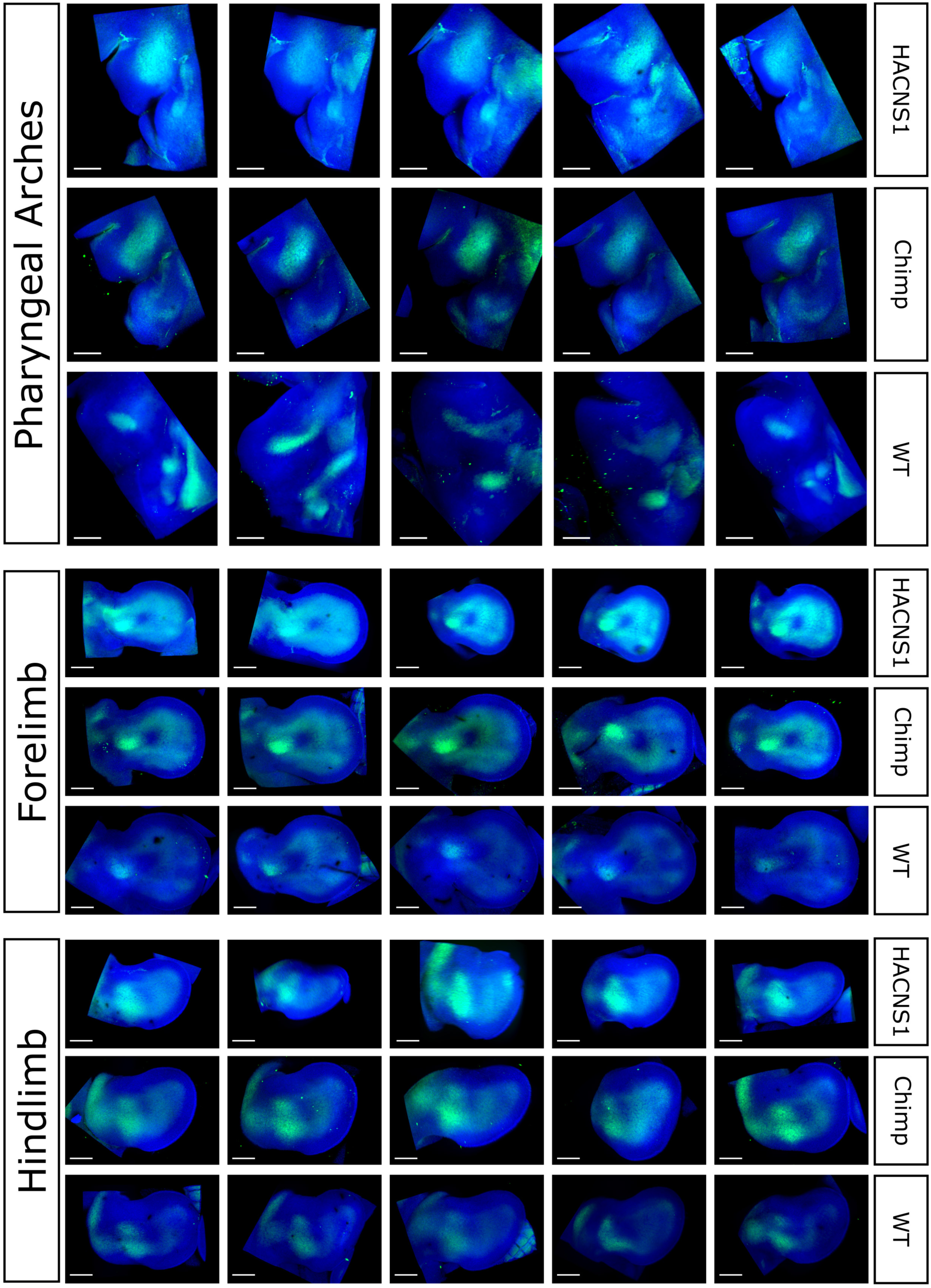
SOX9 expression in E11.5 embryos. All replicates (n=5) of immunofluorescence staining for SOX9 in E11.5 *HACNS1* humanized, chimpanzee ortholog, and wild-type control embryos in the pharyngeal arches (top), forelimb (middle), and hindlimb (bottom). Images were collected from clarified embryos; SOX9 is shown in green, and DAPI nuclear staining is shown in blue. Scale bar, 300 μm. See also Supplemental Movies S13-S21.

**Fig. S24.**
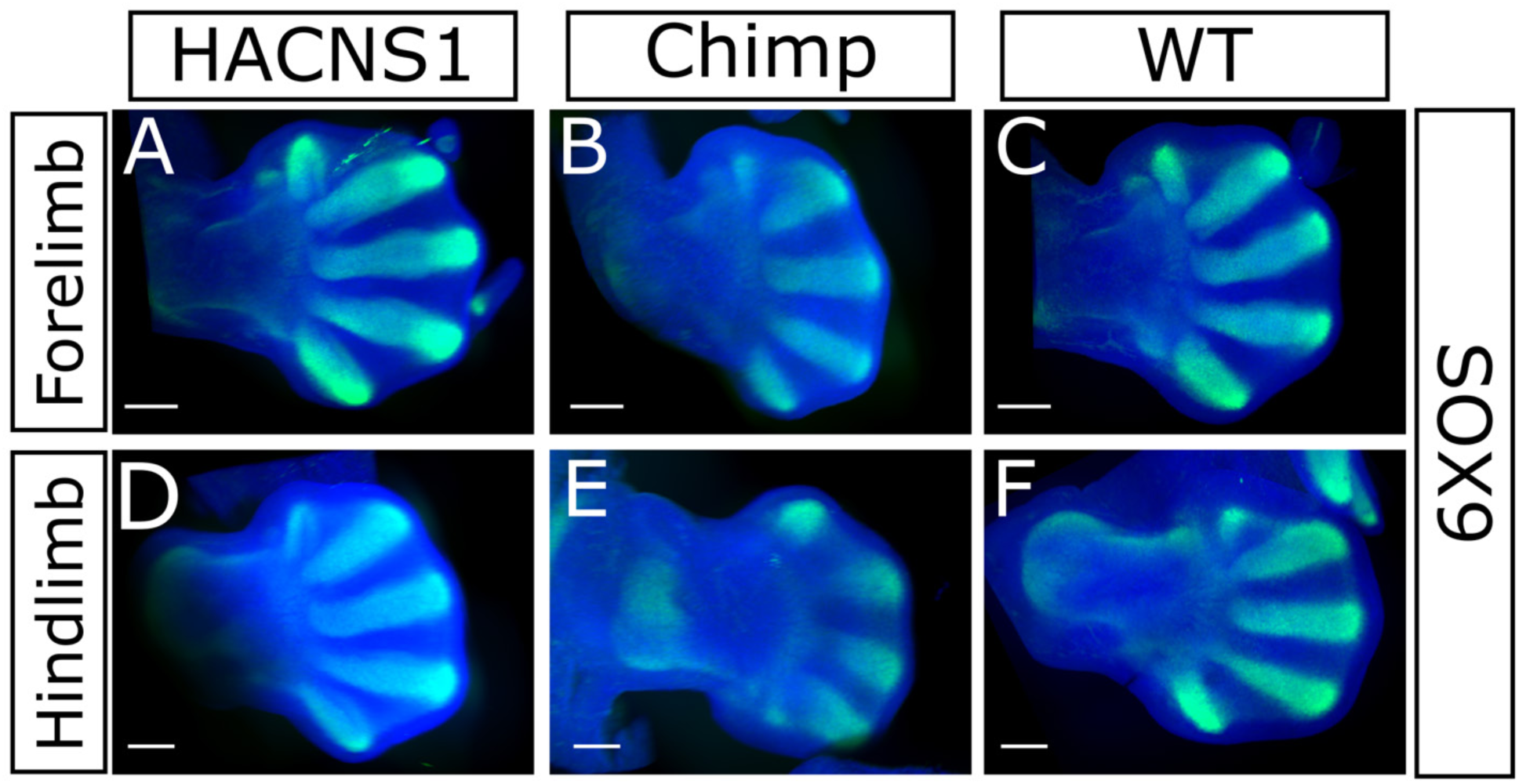
SOX9 expression in E12.5 embryos. Representative immunofluorescence staining for SOX9 in E12.5 *HACNS1* humanized, chimpanzee ortholog, and wild-type forelimb **(A-C)** and hindlimb **(D-F)**. Images were collected from clarified embryos. SOX9 is shown in green, and DAPI nuclear staining is shown in blue. Scale bar, 400 μm; See also Supplemental Movies S22-S27.

**Fig. S25.**
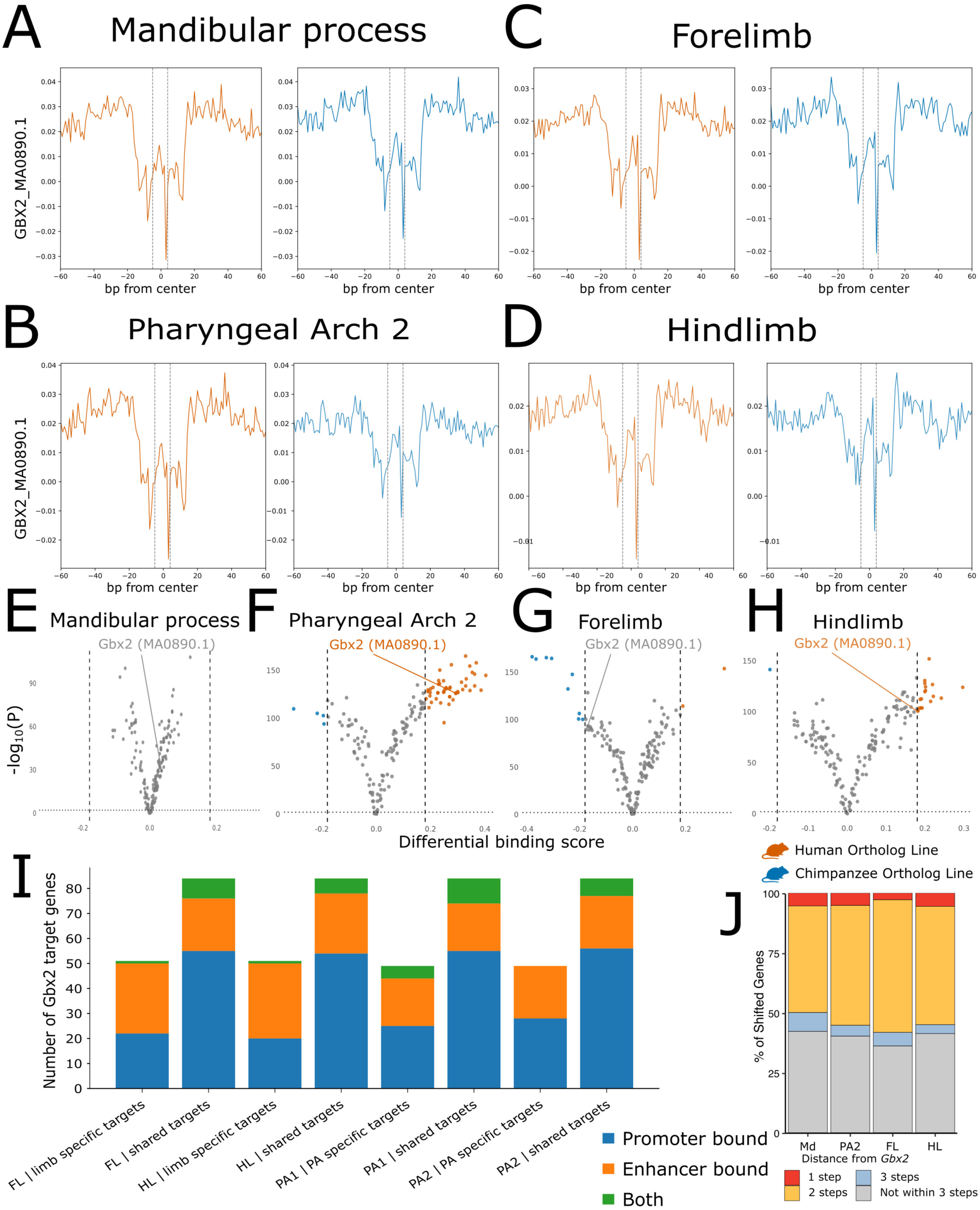
Tissue-specific predicted target genes of *Gbx2* in pharyngeal arches and limb buds. **A-D :** Aggregated footprint plots centered on the predicted binding sites for GBX2 between the *HACNS1* humanized line (dark orange) and the chimpanzee ortholog line (blue) in the Md **(A)**, PA2 **(B)**, forelimb **(C)**, and hindlimb **(D)**. See also Figure 7 and Methods for details. The GBX2_MA0890.1 label to the left of each plot indicates the JASPAR GBX2 motif used to identify predicted GBX2 binding sites for the footprinting analysis (see Methods for more details). The dashed lines represent the edges of the predicted GBX2 binding motif. The horizontal axis shows distance in base pairs from the center of the predicted GBX2 binding site, and the vertical axis in each plot shows the aggregated bias-corrected ATAC-seq signal, and negative values indicate fewer observed cuts than expected after Tn5 bias correction. **E-H:** Pairwise comparison of predicted differential binding between the *HACNS1* humanized line and the chimpanzee ortholog line at E10.5. Volcano plots show differential predicted binding of highly expressed TFs in the Md **(E)**, PA2 **(F)**, forelimb **(G)**, and hindlimb **(H).** See Methods for more details. In each volcano plot, the horizontal axis shows the differential binding score, and the vertical axis shows the −log10(P value) calculated by TOBIAS BINDetect. Orange and blue points indicate motifs with greater predicted binding in the *HACNS1* humanized and chimpanzee ortholog lines, respectively; gray points indicate the remaining motifs. The GBX2 motif is labeled in each plot. **I:** Bar plots showing the number of predicted GBX2 target genes bound at promoters (blue), enhancers (orange), or both promoters and enhancers (green) across tissues. Limb-specific targets were predicted in the limb regulatory networks but not in the PA regulatory networks, whereas PA-specific targets were predicted in the PA regulatory networks but not in the limb regulatory networks. Shared targets were predicted in both. **J:** Stacked bar plots showing the percentage of shifted genes according to their shortest distance in the inferred *Gbx2* regulatory network from *Gbx2* itself. Red, yellow, and blue indicate genes located one, two, or three regulatory steps downstream of *Gbx2*, respectively; gray indicates genes not located within three regulatory steps of *Gbx2*. See also Figure 7 and Methods.

**Fig. S26.**
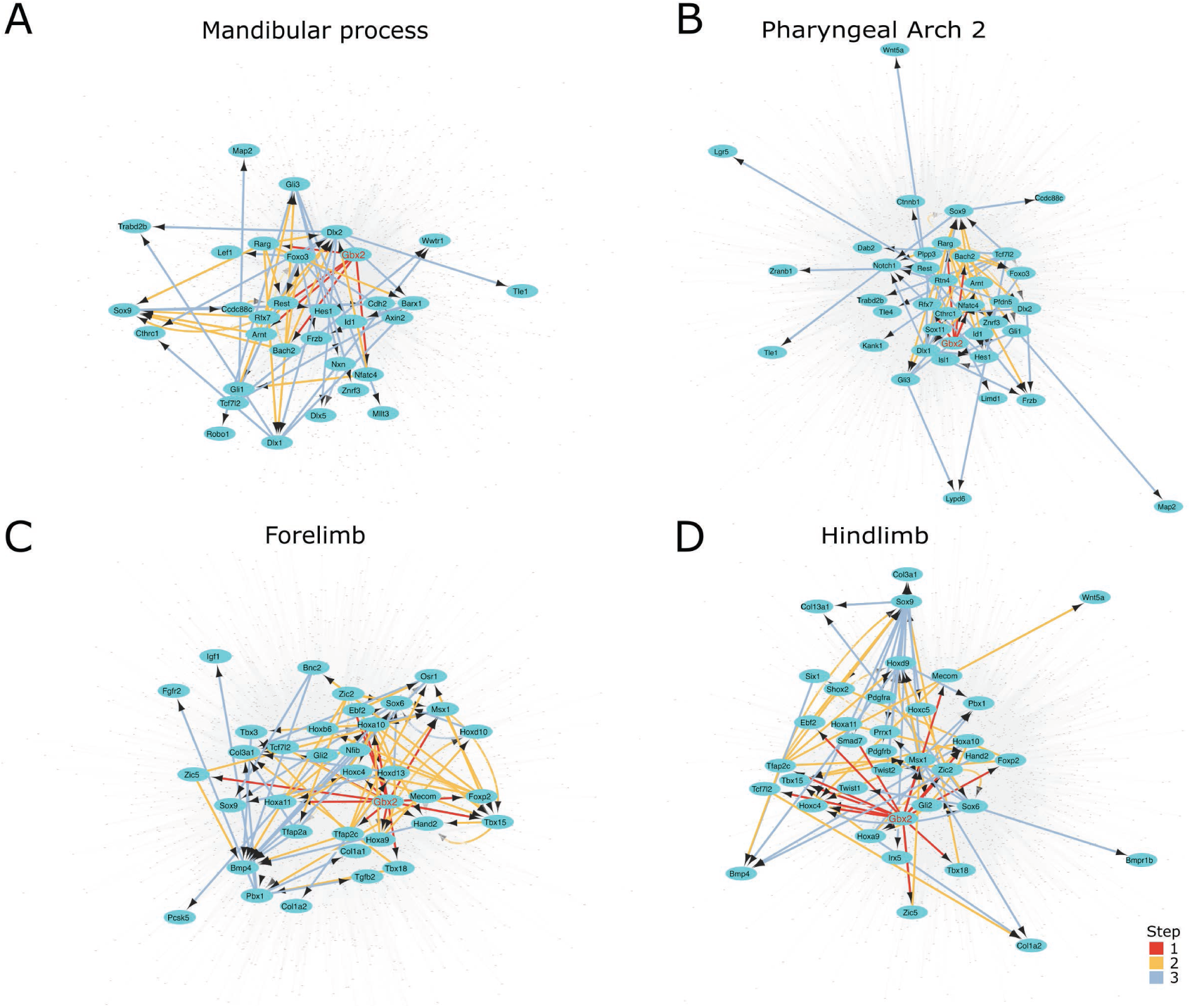
Inferred gene regulatory networks for each tissue. **A-D :** Gene regulatory networks in Md **(A)**, PA2 **(B)**, forelimb **(C)**, and hindlimb **(D)**. Within each gene regulatory network, nodes and edges belonging to the tissue-specific *Gbx2* downstream networks are highlighted, whereas nodes and edges in all other network components are shown in light gray. Nodes represent genes, and arrows point from transcriptional regulators to their target genes. Edges within the highlighted *Gbx2* downstream network are colored according to their distance from *Gbx2*: red, one step; orange, two steps; and blue, three steps. Edge length reflects the GRNBoost2 importance score for each predicted TF-target interaction calculated by pySCENIC GRN, with shorter edges indicating stronger regulatory influence.

## Supplemental Movies

**Movie S1. Three-dimensional PHATE embeddings of mesenchymal cells from the pharyngeal arches.**

Nuclei are colored by cell type, tissue of origin, developmental stage, and genotype.

**Movie S2. Three-dimensional PHATE embeddings of mesenchymal cells from the forelimb.**

Nuclei are colored by cell type, developmental stage, and genotype.

**Movie S3. Three-dimensional PHATE embeddings of mesenchymal cells from the hindlimb.**

Nuclei are colored by cell type, developmental stage, and genotype.

**Movie S4. Immunofluorescence staining of SOX9 in the pharyngeal arches of E10.5 *HACNS1* humanized embryos.**

SOX9 is shown in green, and DAPI nuclear staining is shown in blue. Scale bar, 200 μm

**Movie S5. Immunofluorescence staining of SOX9 in the forelimb of E10.5 *HACNS1* humanized embryos.**

SOX9 is shown in green, and DAPI nuclear staining is shown in blue. Scale bar, 200 μm

**Movie S6. Immunofluorescence staining of SOX9 in the hindlimb of E10.5 *HACNS1* humanized embryos.**

SOX9 is shown in green, and DAPI nuclear staining is shown in blue. Scale bar, 200 μm

**Movie S7. Immunofluorescence staining of SOX9 in the pharyngeal arches of E10.5 chimpanzee ortholog control embryos.**

SOX9 is shown in green, and DAPI nuclear staining is shown in blue. Scale bar, 200 μm

**Movie S8. Immunofluorescence staining of SOX9 in the forelimb of E10.5 chimpanzee ortholog control embryos.**

SOX9 is shown in green, and DAPI nuclear staining is shown in blue. Scale bar, 200 μm

**Movie S9. Immunofluorescence staining of SOX9 in the hindlimb of E10.5 chimpanzee ortholog control embryos.**

SOX9 is shown in green, and DAPI nuclear staining is shown in blue. Scale bar, 200 μm

**Movie S10. Immunofluorescence staining of SOX9 in the pharyngeal arches of E10.5 wild-type control embryos.**

SOX9 is shown in green, and DAPI nuclear staining is shown in blue. Scale bar, 200 μm

**Movie S11. Immunofluorescence staining of SOX9 in the forelimb of E10.5 wild-type control embryos.**

SOX9 is shown in green, and DAPI nuclear staining is shown in blue. Scale bar, 200 μm

**Movie S12. Immunofluorescence staining of SOX9 in the hindlimb of E10.5 wild-type control embryos.**

SOX9 is shown in green, and DAPI nuclear staining is shown in blue. Scale bar, 200 μm

**Movie S13. Immunofluorescence staining of SOX9 in the pharyngeal arches of E11.5 *HACNS1* humanized embryos.**

SOX9 is shown in green, and DAPI nuclear staining is shown in blue. Scale bar, 300 μm

**Movie S14. Immunofluorescence staining of SOX9 in the forelimb of E11.5 *HACNS1* humanized embryos.**

SOX9 is shown in green, and DAPI nuclear staining is shown in blue. Scale bar, 300 μm

**Movie S15. Immunofluorescence staining of SOX9 in the hindlimb of E11.5 *HACNS1* humanized embryos.**

SOX9 is shown in green, and DAPI nuclear staining is shown in blue. Scale bar, 300 μm

**Movie S16. Immunofluorescence staining of SOX9 in the pharyngeal arches of E11.5 chimpanzee ortholog control embryos.**

SOX9 is shown in green, and DAPI nuclear staining is shown in blue. Scale bar, 300 μm

**Movie S17. Immunofluorescence staining of SOX9 in the forelimb of E11.5 chimpanzee ortholog control embryos.**

SOX9 is shown in green, and DAPI nuclear staining is shown in blue. Scale bar, 300 μm

**Movie S18. Immunofluorescence staining of SOX9 in the hindlimb of E11.5 chimpanzee ortholog control embryos.**

SOX9 is shown in green, and DAPI nuclear staining is shown in blue. Scale bar, 300 μm

**Movie S19. Immunofluorescence staining of SOX9 in the pharyngeal arches of E11.5 wild-type control embryos.**

SOX9 is shown in green, and DAPI nuclear staining is shown in blue. Scale bar, 300 μm

**Movie S20. Immunofluorescence staining of SOX9 in the forelimb of E11.5 wild-type control embryos.**

SOX9 is shown in green, and DAPI nuclear staining is shown in blue. Scale bar, 300 μm

**Movie S21. Immunofluorescence staining of SOX9 in the hindlimb of E11.5 wild-type control embryos.**

SOX9 is shown in green, and DAPI nuclear staining is shown in blue. Scale bar, 300 μm

**Movie S22. Immunofluorescence staining of SOX9 in the forelimb of E12.5 *HACNS1* humanized embryos.**

SOX9 is shown in green, and DAPI nuclear staining is shown in blue. Scale bar, 400 μm

**Movie S23. Immunofluorescence staining of SOX9 in the hindlimb of E12.5 *HACNS1* humanized embryos.**

SOX9 is shown in green, and DAPI nuclear staining is shown in blue. Scale bar, 400 μm

**Movie S24. Immunofluorescence staining of SOX9 in the forelimb of E12.5 chimpanzee ortholog control embryos.**

SOX9 is shown in green, and DAPI nuclear staining is shown in blue. Scale bar, 400 μm

**Movie S25. Immunofluorescence staining of SOX9 in the hindlimb of E12.5 chimpanzee ortholog control embryos.**

SOX9 is shown in green, and DAPI nuclear staining is shown in blue. Scale bar, 400 μm

**Movie S26. Immunofluorescence staining of SOX9 in the forelimb of E12.5 wild-type control embryos.**

SOX9 is shown in green, and DAPI nuclear staining is shown in blue. Scale bar, 400 μm

**Movie S27. Immunofluorescence staining of SOX9 in the hindlimb of E12.5 wild-type control embryos.**

SOX9 is shown in green, and DAPI nuclear staining is shown in blue. Scale bar, 400 μm

